# On why we lack confidence in signal-detection-based analyses of confidence

**DOI:** 10.1101/2022.11.07.515537

**Authors:** Derek H. Arnold, Alan Johnston, Joshua Adie, Kielan Yarrow

## Abstract

Signal-detection theory (SDT) is one of the most popular frameworks for analyzing data from studies of human behavior – including investigations of confidence. SDT-based analyses of confidence deliver both standard estimates of sensitivity (d’), and a second estimate based only on high-confidence decisions – meta d’. The extent to which meta d’ estimates fall short of d’ estimates is regarded as a measure of metacognitive inefficiency, quantifying the contamination of confidence by additional noise. These analyses rely on a key but questionable assumption – that repeated exposures to an input will evoke a normally-shaped distribution of perceptual experiences (the normality assumption). Here we show, via analyses inspired by an experiment and modelling, that when distributions of experiences do not conform with the normality assumption, meta d’ can be systematically underestimated relative to d’. Our data therefore highlight that SDT-based analyses of confidence do not provide a ground truth measure of human metacognitive inefficiency.

**Public Significance Statement:** Signal-detection theory is one of the most popular frameworks for analysing data from experiments of human behaviour – including investigations of confidence. The authors show that the results of these analyses cannot be regarded as ground truth. If a key assumption of the framework is inadvertently violated, analyses can encourage conceptually flawed conclusions.

## Introduction

When people make decisions, we experience a subjective level of confidence regarding the quality of our decisions (Fleming et al., 2012; Yeung & Summerfield, 2012). In perception, these feelings are typically accurate, with higher levels of confidence associated with more accurate decisions (De Martino et al., 2013; Keane et al., 2015; Li et al., 2014; Peters et al., 2017). This capacity – to report on the accuracy of our intrinsic decisional processes, is known as perceptual metacognition.

Historically, quantifying the degree of insight we might have into the internal operations of our minds has been challenging. The recent extrapolation of a standard framework, traditionally used to estimate perceptual sensitivity (Green & Swets, 1966), to appraise confidence has therefore been noteworthy – promising an objective measure of the degree of insight we have into the quality of our perceptual decisions (Fleming & Lau, 2014; Maniscalco & Lau, 2012).

To understand the new approach, we need to start with a reprisal of standard implementations of the SDT framework (Green & Swets, 1966; Yarrow et al., 2011). Popular implementations of the framework commit to a set of assumptions that allow investigators to calculate independent estimates of objective sensitivity (d’), and of the subjective criteria people use when making perceptual decisions. One of the key assumptions is that repeated exposures to a given physical input will be associated with a normally-shaped distribution of different perceptual experiences (Fleming & Lau, 2014; Green & Swets, 1966; Maniscalco & Lau, 2012; Yarrow et al., 2011). For instance, a vertical input might most often be perceived as vertical, but also be seen as differently tilted, left and right of vertical on some trials. It is assumed that the various experiences following exposures to a common physical input will be normally distributed. It is this assumption that allows researchers to make a backwards inference, from behavioral data, to estimate the areal overlap between theoretical distributions that describe the different experiences that have ensued following exposures to a target (signal) and to a non-target (noise) input (see Figure 1).

**Figure 1.**
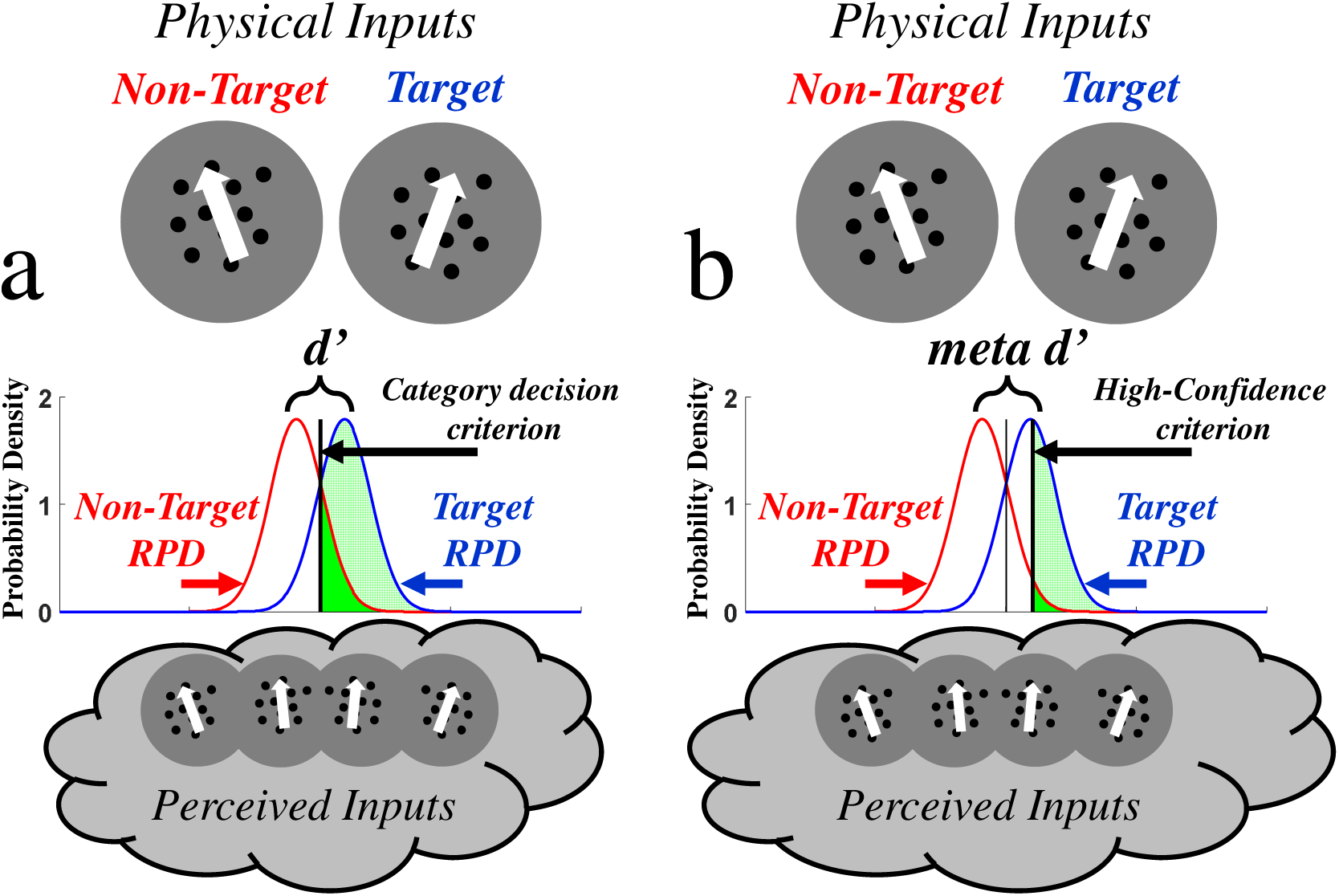
**a)** Graphic depicting popular implementations of the SDT framework. Repeated presentations of a non-target and of a target input (here dots moving left and right of vertical – see top of graphic) will result in different perceptual experiences (different perceived directions – see bottom of graphic). These are thought to be described by normal distributions. It is further assumed that people reference experiences against a category criterion value (vertical black bar) to demark non-target (values to the left of the criterion) from target experiences (values to the right of the criterion). These assumptions allow researchers to backwards infer how separated the target and non-target distributions must be to be consistent with the proportion of non-target presentations categorized as targets (the false alarm rate – see dark green shaded region) and with the proportion of target presentations categorized as targets (the hit rate – see light green shaded region). This estimate – d’ – is regarded as a measure of perceptual sensitivity. **b)** Graphic depicting assumptions underlying popular extensions of the SDT framework, used to analyze confidence. A high-confidence criterion (bold black vertical line) is assumed to demark experiences classified as having been of targets with high confidence (values to the *right* of the high-confidence criterion. This criterion is applied to both non-target (dark green shaded region) and to target (light green shaded region) presentations. Researchers can then backwards infer how separated target and non-target distributions must be, to be consistent with proportions of *correct* (dark green shaded region) and *incorrect* (light green shaded region) high-confidence target categorizations. This estimate is known as meta-d’.

To assess confidence, the standard SDT framework has been extended to include additional ‘confidence’ criteria – values against which experiences are referenced to assign levels of confidence (see Figure 1b). In a minimal case, you need two additional confidence criteria, to demark low-from high-confidence decisions. You could, for instance, have a criterion to demark low-from high-confidence target categorisations (see the bold black vertical bar, Figure 1b). With the addition of these criteria, researchers can backwards infer a second estimate of how separated non-target noise and target signal distributions must be, from an analysis restricted to proportions of correct (Figure 1b, light green shaded area) and incorrect (see Figure 1b, dark green shaded area) high-confidence target categorisations (Fleming & Lau, 2014; Maniscalco & Lau, 2012). In essence, this process estimates the same property as more traditional SDT-based analyses (d’), but analyses are restricted to the right-side tails of distributions – where responses have elicited a high-level of confidence. The key statistic this process delivers is meta d’.

When meta d’ and d’ estimates are equivalent, an observer is said to be metacognitively ‘ideal’, in that their data are consistent with d’ and meta d’ estimates having been informed by a common source of information. The degree to which an observer is said to be metacognitively inefficient is thought to be indicated by the degree to which meta d’ estimates fall short of d’ estimates (Fleming & Lau, 2014; Maniscalco & Lau, 2012). This situation is typically regarded as evidence that estimates of confidence have been contaminated by an additional source(s) of noise, relative to perceptual judgments (Fleming & Lau, 2014; Maniscalco & Lau, 2012). Standard implementations of this framework rest on the assumption that distributions of experiences after repeated exposures to an input are normally distributed, such that areal overlaps of distributions can be inferred from behaviour.

A structural concern for SDT-based analyses of confidence is that we don’t know the precise shape of experiential distributions, and there is good reason in visual perception to suspect these might often deviate from normality. The assumption of normally-shaped experiential distributions presumes that an input is equally likely to be mis-perceived to either side of its veridical position along a psychological dimension. A near vertical input, for instance, should equally be likely to be misperceived as vertical or as further tilted from vertical. Evidence, however, suggests near vertical inputs are more likely to be misperceived as vertical than as further titled from vertical – implying an intrinsic local skew at this part of this particular psychological dimension (Appelle, 1972; Girshick et al., 2011; Storrs & Arnold., 2015). Perceived motion directions are similarly biased toward being seen to move along cardinal (vertical and horizontal) as opposed to oblique angles (Appelle, 1972; Dakin et al., 2005).

In addition to an absence of skew, the assumption of normally-shaped experiential distributions (Fleming & Lau, 2014; Maniscalco & Lau, 2012) presumes that experiential distributions should be mesokurtic, such that the shapes that describe the tails of an experiential distribution should be matched to those that describe the tails of a normal distribution. However, it has been established that detailed datasets describing human decisions can be equally well explained by decisions having been informed by a number of differently shaped experiential distributions (see Rouder et al., 2010). This observation has been advanced as motivation to avoid excessive dependence on assuming a particular shape of experiential distribution within analyses of perceptual decisions (e.g. Kellen & Klauer, 2015; Miyoshi et al., 2022).

Even if inputs were usually mapped onto different perceptual experiences, mappings between inputs and perception can be made to undergo temporary changes, like those invoked by visual adaptation (by prolonged exposure to a given physical input – such as upwards motion). The most obvious changes following visual adaptation are to perception – with a tendency to see similar inputs as repelled from the adaptor (Clifford et al., 2000; Gibson & Radner, 1937; Regan & Beverly, 1985; Webster, 2015). Adaptation to upwards motion, for instance, can make movements to the left and right of vertical seem to be moving in directions more rotated from upwards (Clifford et al., 2000; Webster, 2015). These perceptual changes are accompanied by less obvious changes to the precision of perceptual judgments, with the direction and magnitudes of both types of change being governed by the proximity of inputs to the adaptor within the psychological dimension (Clifford et al., 2001; Regan & Beverly, 1985; Webster, 2015). Recently, it has been shown that visual adaptation can also impact on measures of perceptual confidence (Arnold et al., 2021).

A benefit of visual adaptation is that we can leverage knowledge of the processes underlying human vision to produce biologically-inspired models that detail how changes in perception, in the precision of perceptual decisions, and in confidence, might all have been produced (Clifford et al., 2001; Jin et al., 2005; Kohn & Movshon, 2004; Kohn, 2007; Storrs & Arnold, 2012; Storrs & Arnold, 2015). An important conceptual point is that the success of these models depends on a description of how adaptation might have temporarily changed mappings between inputs and perception, with distinct changes prevailing at different loci along a psychological dimension. Our recent study showed that this approach could describe how tilt adaptation impacts measures of human perception, perceptual precision, and most importantly confidence – specifically the spread of uncertainty (i.e. a lack of confidence). Uncertainty tended to be increased by adaptation, to a greater extent than the accompanying adverse changes to the summary statistics describing the precision of perceptual judgments (Arnold et al., 2021).

An additional advantage of biologically inspired models of perception (Clifford et al., 2001; Jin et al., 2005; Kohn & Movshon, 2004; Kohn, 2007; Storrs & Arnold, 2012; Storrs & Arnold, 2015), untapped until now, is that these can be quizzed to explore what shape experiential distributions might have, before and after visual adaptation.

Here we will report on a study that had four stages. First, we demonstrate the robustness of findings from our past investigation, by assessing the impact of visual adaptation on confidence in a different perceptual dimension (direction perception). To preface these results, confidence is again robustly impacted by adaptation. In a second stage, we show that our behavioural datasets can be similarly accounted for by a biologically-inspired model, which describes how mappings between inputs and aspects of perception might be changed by adaptation. In a third stage, we estimate what shape pre- and post-adaptation experiential distributions might have from modelling. These analyses suggest that experiential distributions could be both skewed, and have excess kurtosis, and that both of these outcomes can undermine SDT-based analyses of confidence.

The final phase of our study is shortest, but in our minds most consequential. We show that the degree to which SDT-based analyses of confidence underestimate meta d’ (relative to d’) can scale with the degree to which the experiential distributions informing these analyses are skewed, or have excess kurtosis. The results of these analyses are not contingent on the veracity of the first three stages of our study, but are rather a direct computational consequence of SDT-based analyses of confidence being informed by probability distributions that are abnormally shaped.

## Methods

Nineteen volunteers participated, 14 female, with a mean age of 23 (S.D. 2.6). All were experienced psychophysical observers by the end of the experiment, but almost all were novice at the beginning. Eighteen of the volunteers participated as partial fulfillment of the requirements of an undergraduate course, and the remaining participant is the third author of the study. All participants completed experimental sessions while seated in a dimly lit room, viewing stimuli from a distance of 57 cm with their head restrained by a chin-rest. The study was approved by The University of Queensland research ethics committee, and was conducted in accordance with the principles of the Declaration of Helsinki. Participation involved ∼11 hours of testing for each observer, split across 11 experimental sessions (usually conducted on different days).

## Stimuli

Stimuli consisted of random dot kinematograms (RDKs), generated using a Cambridge Research Systems ViSaGe stimulus generator driven by custom Matlab R2013b (MathWorks, Natick, MA) software and presented on a gamma-corrected 19 inch Dell P1130 monitor (resolution: 1600 x 1200 pixels; refresh rate: 60 Hz). Each RDK consisted of 100 individual black dots, subtending 0.02 degrees of visual angle (dva) in diameter at the retinae, presented within grey (luminance = 51 cd/m2) circular apertures, with a diameter of 5.5 dva (see Figure 2). The individual lifetime of each dot was 100ms (6 frames), after which it was re-drawn at a random position within the aperture. Each dot was assigned a random initial age (between 0 and 100ms), so when there was no coherent movement in an RDK, dot up-dates created directionless flicker.

**Figure 2.**
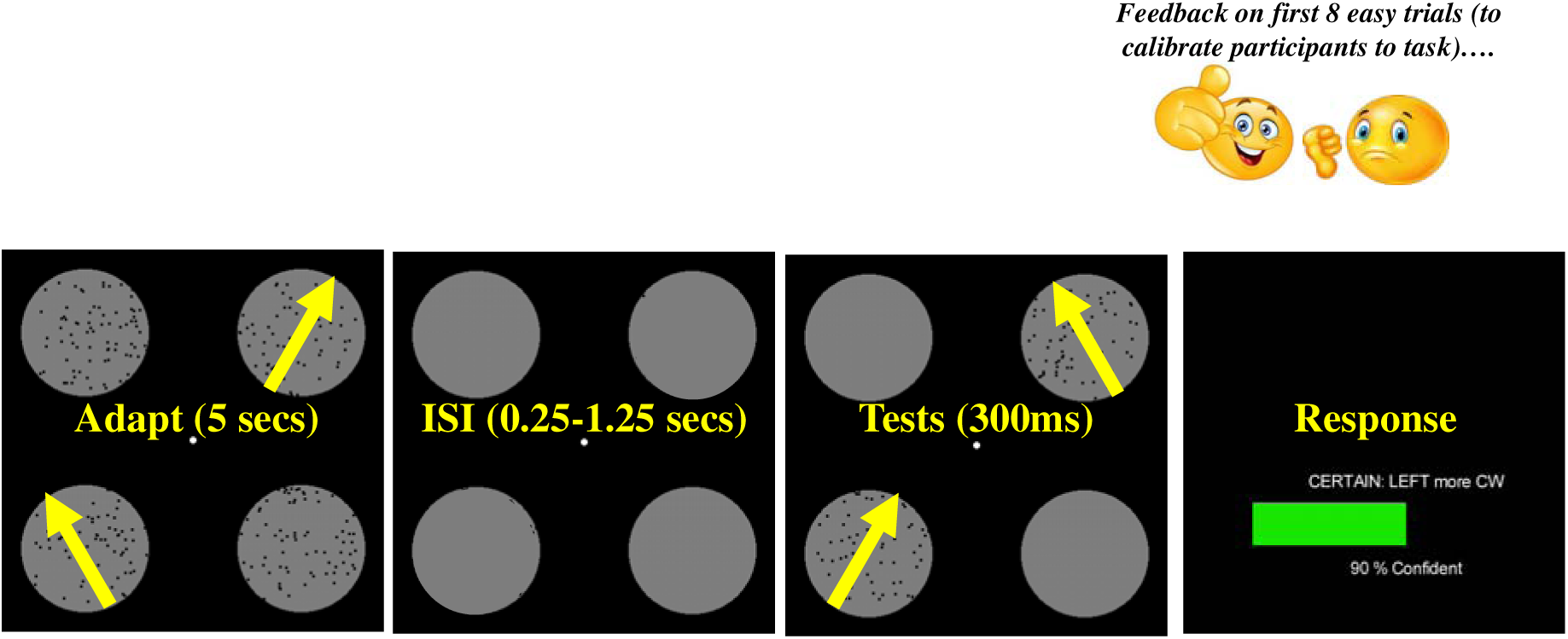
Graphic depicting the experimental protocol. On each trial participants were adapted (for 5 secs) to a pair of RDKs, diagonally positioned relative to the central fixation point (e.g. toward the bottom left and top right). Adapting motion directions were oppositely rotated from vertical, by one of a range of angular magnitudes. A single adapted direction difference prevailed throughout each block of trials. After variable inter-stimulus-intervals, participants viewed a brief pair of test directions, also oppositely rotated from vertical by one of a range of different angular magnitudes. Participants then simultaneously indicated which test had contained a more clockwise direction of motion, and the level of confidence they felt in this decision, by making a setting along a linear scale. Feedback was provided on the first 8 trivially easy trials, to (re)acquaint participants with the experimental task, while avoiding contaminating any intuitive insights into task performance, by training people to recognize when they had made correct or incorrect decisions about ambiguous inputs.

In the center of test displays there was a white (luminance = 102 cd/m2) static disc subtending 0.02 dva in diameter – which served as the fixation point. Participants were asked to fixate on this throughout each experimental session. The display was otherwise black. The 4 apertures that could contain RDKs were centered 6.9 dva to the left, right and above and below the central fixation point (see Figure 2).

## Procedure

All trials had a matched sequence. They began with a 5 second adaptation period, in which two of four RDKs contained dots moving coherently in directions oppositely rotated from upward. The other two contained directionless flicker. There was then a 0.25 to 1.25 second inter-stimulus-interval (ISI), wherein no RDKs were presented, but the grey circular apertures could be seen. Two test RDKs, with dots also moving in oppositely rotated directions from upward, were then shown for 300ms. These were either presented in the same positions as the coherently moving RDKs (Adaptation trials), or in the same positions as flickering RDKs (Baseline trials). Participants were prompted to make a combined response, indicating which test had seemed to contain movement in a more clockwise direction (a categorical perceptual decision), and the degree of confidence they felt in this decision (by making a setting along a linear scale, with a minimal requirement of 5%, up to a maximum of 100%). Feedback was given regarding the accuracy of perceptual decisions on an initial 8 calibration trials in each experimental session. Test differences on these trials were set to the maximal possible test difference, to (re)familiarise participants with the task, while avoiding contaminating intuitive insights into task performance by training people to recognize when they had made a correct or incorrect decision about a perceptually ambiguous input.

Participants were adapted to different magnitudes of direction difference, between the two adapting RDKs, in different experimental sessions. Direction differences of 0 to 360° were sampled, in steps of 40°, with left-side adapting stimulus movements rotated counter-clockwise from upward in steps of 20°, and right-side vice versa, such that at an adapting direction difference of 360°, both adaptors contained downward motion. Each participant completed 10 adaptation conditions, in an individually randomized order.

During each experimental session, the magnitudes of test direction differences were adjusted according to 1-up 1-down staircase procedures (Levitt, 1971), with decisions indicating that the right-side stimulus had moved in a more clockwise direction resulting in subsequent right-side tests moving in a more counter-clockwise direction, and vice versa. Four staircases were interleaved, two for each experimental condition (Baseline and Adaptation). One of these was initiated with right-side tests moving in a maximally clockwise rotated direction (given the range of test directions), and one was initiated with right-side tests moving in a maximally counter-clockwise direction. For most participants, maximal test directions were rotated +/-16° from upward, which was then adjusted in 2° steps. Some participants, however, had more difficulty performing the task, so for these participants maximal test directions were rotated +/-32° from upward, and adjusted in 4° steps.

The first 8 trials of each experimental session were calibration trials, after which staircases were not updated. There were then an additional 100 trials for each experimental condition, all randomly interleaved – for a combined total of 208 individual trials in each experimental session. Staircases were re-set midway through experimental sessions, to ensure large test differences were sampled at least twice.

Individual data for each condition recorded during an experimental session were collated. Confidence ratings were then categorised, as high or low, relative to the median baseline confidence rating during that experimental session. A raised Gaussian function was fit to datasets describing low-confidence decisions as a function of test direction differences. The peaks of these functions were taken as estimates of test directions Perceived to be Subjectively Equivalent (Confidence PSE estimates), and the Full-Width at the Half-Height (FWHH) of these functions were taken as estimates of the range of test directions that had elicited low-confidence decisions (a measure of the spread of uncertainty).

Cumulative Gaussian functions were fit to data describing the proportion of right-side tests seen as having moved in a more clockwise direction, as a function of physical test direction differences. The 50% point of these functions were taken as estimates of test directions Perceived as Subjectively Equal (Perception PSE estimates). Distances between 25 and 75% points of fitted functions were taken as estimates of the Just Noticeable Difference (JND) between test directions. Note that this last statistic is a standard measure of the precision of perceptual judgments.

Experimental sessions were usually completed on separate days, with breaks taken in-between experimental sessions when multiple sessions were completed on the same day. Data informing individual aftereffect functions are generated from 2080 individual trials.

## Data Availability / Pre-Registration

All data and analyses described in this paper are available as supplemental material. This study was not pre-registered.

## Results

### Stage 1: Behavioural Experimental Results

Each experimental session delivered two datasets describing proportions of right-side tests categorised as having moved in a more clockwise direction from vertical, relative to left side tests – in each case as a function of physical direction differences. One was from baseline tests, and one from adapted tests. Cumulative Gaussian functions were fit to each, and 50% points were taken as estimates of Perceptual Points of Subjective Equality (Perceptual PSEs). The distance in-between points on the x-axis, corresponding with the 25 and 75% points on the y-axis, were taken as estimates of Just Noticeable Differences (Perceptual JNDs) between test directions – a standard measure of the precision of perceptual decisions, with smaller JNDs indicating greater precision.

Each experimental session also delivered two datasets describing proportions of low-confidence perceptual decisions, as a function of physical direction differences – one for baseline and one for adapted tests. Raised Gaussian functions were fit to these, and peaks were taken as Confidence PSE estimates (under the assumption that greatest uncertainty should ensue when test directions are perceptually matched). The distance along the x-axis corresponding with the Full Width of fitted functions at their Half Heights (Confidence FWHHs) were taken as estimates of the spread of uncertainty.

Adaptation-induced aftereffects were calculated from each experimental session, by subtracting adapted from unadapted summary statistics (i.e. adapted Perceptual PSEs – baseline Perceptual PSEs, etc…). Aftereffects expressed as a function of adapted direction differences, averaged across all participants, are depicted in Figure 3f-g. Adaptation to directions rotated +/-∼20 to 100° from upward repelled perceived test directions away from adapted directions (see Figure 3f, blue data for Perceptual PSE aftereffects, red data for Confidence PSE aftereffects). It is important to note that these aftereffect functions, calculated from perceptual decisions and from confidence ratings, are very similar – suggesting the two tasks have provided equivalent measures of changes in perceived direction.

**Figure 3.**
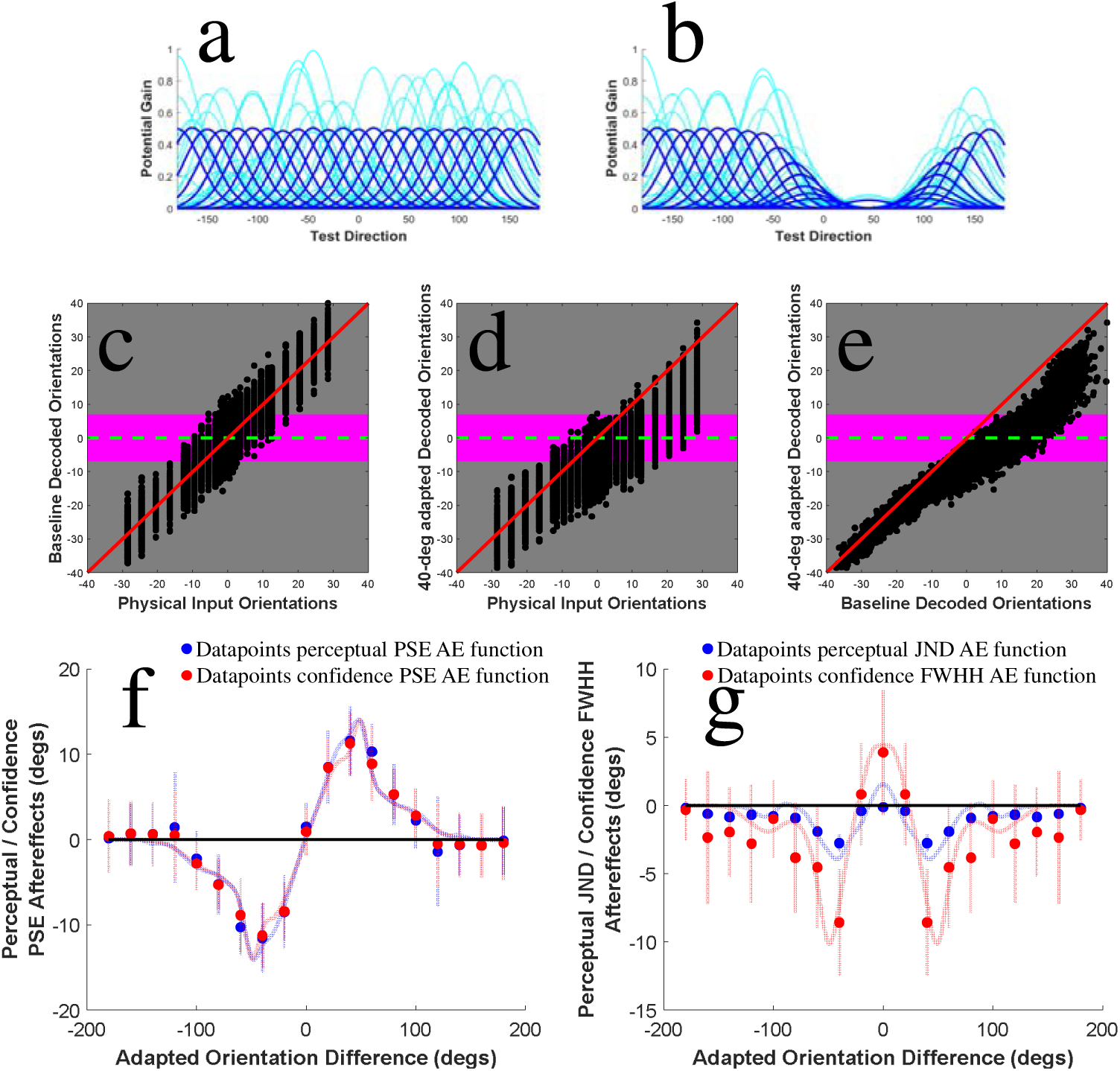
Depictions of **a)** unadapted (Baseline) model channels, and **b)** model channels adapted to a +45° direction. Faint blue lines depict response potentials on 3 simulated trials. Potential channel responses, averaged across 1584 trials, are also depicted (bold blue lines). **c)** X/Y scatter plot of encoded (Y axis) and physical (X axis) directions across 1584 simulated trials by the baseline model. Each physical input is encoded differently on discrete trials, due to simulated encoding noise, but on average the baseline model encodes inputs veridically, so datapoints cluster about the red oblique line (plotting veridical 1:1 input decodings). The horizontal green dotted line depicts the criterion value for perceptual categorizations (0°), and the edges of the horizontal pink rectangle depict the unsigned magnitude criterion for confidence categorizations (as low or high). **d)** Details are as for Figure 3c, but for the same simulated trials decoded by a +45° adapted model (with individual channels subject to the same nominal noise levels). **e)** Differences between perceived directions decoded by baseline (X axis) and +45° adapted (Y axis) models. **f)** Observer model fits (faint lines) to changes in both Perceptual (blue data) and Confidence PSEs (red data). Aftereffects following adaptation to negative direction differences are assumed to be mirror opposites of positive aftereffects. **g)** Observer model fits (faint lines) to changes in both Perceptual JNDs (blue data) and Confidence FWHHs (red data). Again, aftereffects following adaptation to *negative* direction differences are assumed to be mirror opposites of positive aftereffects (see Figure 3d). Model fits capture key qualitative features of all four aftereffect datasets.

Adaptation to tests rotated +/-40° from upward produced an increase in Perceptual JNDs (plotted as negative values, i.e. baseline - adapted JNDs, as this is indicative of reduced precision). This is not obvious for other adaptors (see Figure 3g, blue data). Adaptation-induced reductions were greater in magnitude, and more widespread, for Confidence FWHHs (see Figure 3g, red data). There was also a small benefit to confidence from adapting to the average test value (0° adaptation), which was not apparent for Perceptual JNDs. In sum, the impact of adaptation on estimates of the spread of uncertainty was greater than the impact of adaptation on estimates of the precision of perceptual judgments (see Figure 3g).

### Stage 2: A biologically-inspired model to account for observed behavioural changes

To simultaneously account for the impact of adaptation on 1) direction perception, 2) the precision of direction judgments, and 3) confidence in direction judgments, we created a labelled line observer model (Arnold et al., 2021; Clifford et al., 2001; Jin et al., 2005; Kohn & Movshon, 2004; Kohn, 2007; Storrs & Arnold, 2012; Storrs & Arnold, 2015). This assumes that sensory information is encoded as a pattern of responses to inputs from across a population of ‘channels’, each maximally responsive to a different direction (see Figure 3a-b). In our model the potential response of each channel to inputs is described by a normal distribution, with a standard deviation of 20°. Peak potential responses (channel tunings) are separated by 10°, ranging from ±180° (downward motion) to +170° in 10° steps—so our model has 36 channels.

The neural consequences of visual adaptation include reduced responding to inputs, and changes to both the optimal direction and to the range of directions that elicit responses (Pallier et al., 2002). In our model these effects are operationalized by implementing a reduction in the response potential of model channels (up to a maximum of 95%) in proportion to how responsive each channel is to the adaptor in an unadapted state (Clifford et al., 2000; Jin et al., 2005; Kohn & Movshon, 2004; see Figure 3b), and by applying model channel tuning shifts (up to a maximum of 40°) away from adapted directions (Moreira et al., 2018; Pallier et al., 2002; see Figure 3b), also in proportion to how responsive channels are to the adaptor in an unadapted state. For instance, after adaptation to 40°, channels tuned from - 50 to +130 (in 10° steps) would respectively be subject to a 15, 29, 43, 56, 67, 77, 85, 90, 94, 95, 94, 90, 85, 77, 67, 56, 43, 29 and to a 15% reduction in response potential to inputs. Also after adaptation to 40°, channels tuned to 10, 20 and 30° would be subject to -28, -40 and to - 28° tuning shifts, whereas channels tuned to 50, 60 and to 70° would be subject to tuning shifts of +28, +40 and to +28 degrees. We operationalize trial-by-trial neural encoding noise by applying a random scaling (drawn from a uniform distribution, ranging from 0 to 100%) to the 36 response potentials of all model channels (see Figure 3a-b; model code is provided as Supplemental material code #1).

Our model encodes a perceived direction value on each simulated trial by taking a weighted sum of the direction labels of each channel. Each label is first multiplied by each channel’s response (including noise) then divided by the summed magnitude of all channel responses (including noise). Categorical perceptual decisions (i.e. is the stimulus moving clockwise or counter-clockwise from vertical?) are decoded from adapted and unadapted (baseline) states of the model by comparing the weighted sum of all channel responses (the model response) to a criterion value (0°). Categorical confidence decisions (high / low) are decoded by indexing unsigned model response magnitudes against a criterion value (of 7°, so inputs decoded as moving > +/-7° from vertical elicit a high-confidence rating, and smaller decoded values elicit a low confidence rating). This confidence criterion was set to match half the average baseline Confidence FWHH (14°, S.D. 2.33), which equates to ∼3x the average baseline Perceptual JND (4.5°, S.D. 1.29).

Model data from a simulated experiment are shown in Figure 3c-e. Repeated trials with identical physical inputs result in different encoded values, both at baseline (Figure 3c) and after adaptation to a +40° direction (Figure 3d), in each case due to random trial-by-trial changes in the response potentials of all model channels. Differences between values encoded by the baseline and by the +40° adapted model are also shown (Figure 3e). It is important to note that our model produces human like performance when categorising inputs. In the simulated experiments cumulative Gaussian functions are fit to distributions describing the proportion of inputs encoded by the model as having moved in a clockwise direction relative to upwards (see Figure 4). The fact that these distributions describe probabilistic functions is due to the model, like humans, encoding a given input differently on repeated trials, due to trial-by-trial encoding noise.

**Figure 4.**
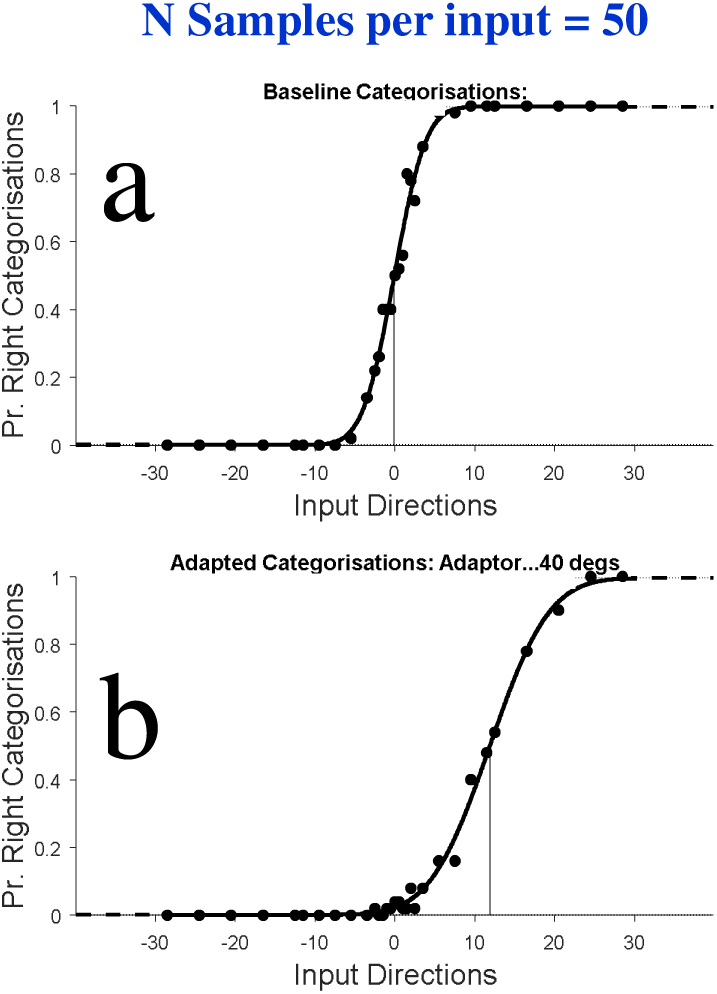
Cumulative Gaussian functions fit to data describing the proportion of inputs encoded by our model as having moved in a clockwise direction relative to vertical as a function of input test values. Functions are fit to data encoded by our **(a)** baseline and by our **(b)** +40° adapted model. Note that both functions are probabilistic, as trial-by-trial encoding noise is intrinsic to the model. The model also captures both changes in perception post +40° adaptation (note the rightward shift of the adapted relative to the baseline function, and compare this to our human behavioral data, depicted in Figure 3f) and changes in the precision of perceptual judgments (note the reduced slope of the adapted relative to the baseline function, which is commensurate with the reduced precision of our human behavioral judgments, depicted in Figure 3g).

Our model provides a good account of the key qualitative features of all four behavioural aftereffect functions, describing the impact of adaptation on perceived direction (Figure 3f), on estimates of the precision of perceptual judgments (Figure 3g – blue data) and on the spread of uncertainty (Figure 3g – red data). All four functions have been fit simultaneously (i.e. with a common set of model parameters).

It is often claimed that labelled line models have few free parameters (e.g. Clifford et al., 2000). The behaviour of this class of models is, however, the product of a complex interplay between many factors. In this case, this has included the number (36) and spacing (10°) of model channels, their tuning bandwidths (20°), the criterion values chosen to classify data (0° for perceptual categorisations, and an absolute, unsigned, value of 7° for high-confidence categorisations), the magnitude of post-adaptation reductions of potential channel responses (up to 95%), the magnitude of post-adaptation tuning shifts (up to 40°), and the spread of the last two types of change across model channels. Of these, post-adaptation reductions in potential channel responses are most important to bias encoded directions away from adapted directions (see Figure 3f), and tuning shifts are most important for generating localized changes in the precision of perceptual judgments and confidence (see Figure 3g). Other factors tend to govern the spread of these changes.

We arrived at the parameter settings for our model via a process of educated adjustments, to achieve good simultaneous fits to all four of our aftereffect functions (which were in each case averaged across all participants). Given the high dimensionality of our model, we would not describe it in any way as optimal, or suggest it is superior to any related model. We would note, however, that other approaches to modelling often achieve a minimum of free parameters by assuming the existence of distributions of a specified shape (e.g. Locke et al., 2022; Maniscalco & Lau, 2016; Shekhar & Rahnev, 2021), and one thing our model demonstrates is that if the neural processes that might contribute to determining the shape of a distribution are made explicit, a host of assumptions might need to be implemented that are otherwise hidden. Regardless, we advance our model simply as an existence proof, that the key qualitative features of our four aftereffect functions can be described by a biologically inspired model. This motivated the next phase of our analyses, to see what shapes would describe the distributions encoded by our model given repeated inputs.

### Stage 3: What shapes describe the experiential distributions output by the model, which successfully describes human behaviour in our experiment?

Stage 2 analyses modelled data for the baseline, and for each of the 10 adaptive conditions sampled in our behavioural experiment. For Stage 3 analyses, we will examine the products of a baseline and a +40° adapted model, as this adaptation condition was associated with the greatest adaptation-driven changes in our behavioural experiment. We simulated SDT experiments using our model to determine if repeated exposures to a given input, before and after adaptation, might result in normally-shaped distributions of different encodings, or in abnormally-shaped distributions, in order to see if any deviance from normality would have a discernible impact on SDT-based analyses of confidence.

We simulated experiments for an unbiased observer, with ‘signal’ and ‘noise’ tests rotated by one of 101 different magnitudes (from +/-1.5 to 2.7°) from a central perceptual category boundary – the direction encoded by the relevant model as upwards (∼0° for our unadapted Baseline model, ∼11.5° for our +40° adapted model). Confidence criteria were set to +/-7° from the central category boundary criterion (note that this confidence criterion is matched to the baseline performance of our human participants). This allows model encodings on each trial to be indexed against a central perceptual category boundary, and against confidence criteria to determine if a trial should be categorised as a noise presentation with high confidence, as a noise presentation with low confidence, as a signal presentation with low confidence, or as a signal presentation with high confidence (see Figure 1b).

This treatment of data allows us to implement a popular SDT-based analysis of confidence (Maniscalco & Lau, 2012) to arrive at d’ and meta d’ estimates (code for this analysis is provided as Supplemental material code #2). Given that our model has no metacognitive bias (i.e. the same encodings inform both d’ and meta d’ calculations), if the model were producing normally-shaped distributions of encoded values, d’ and meta d’ estimates should fall along a line describing a 1:1 slope (marked by green oblique lines in Figure 5).

**Figure 5.**
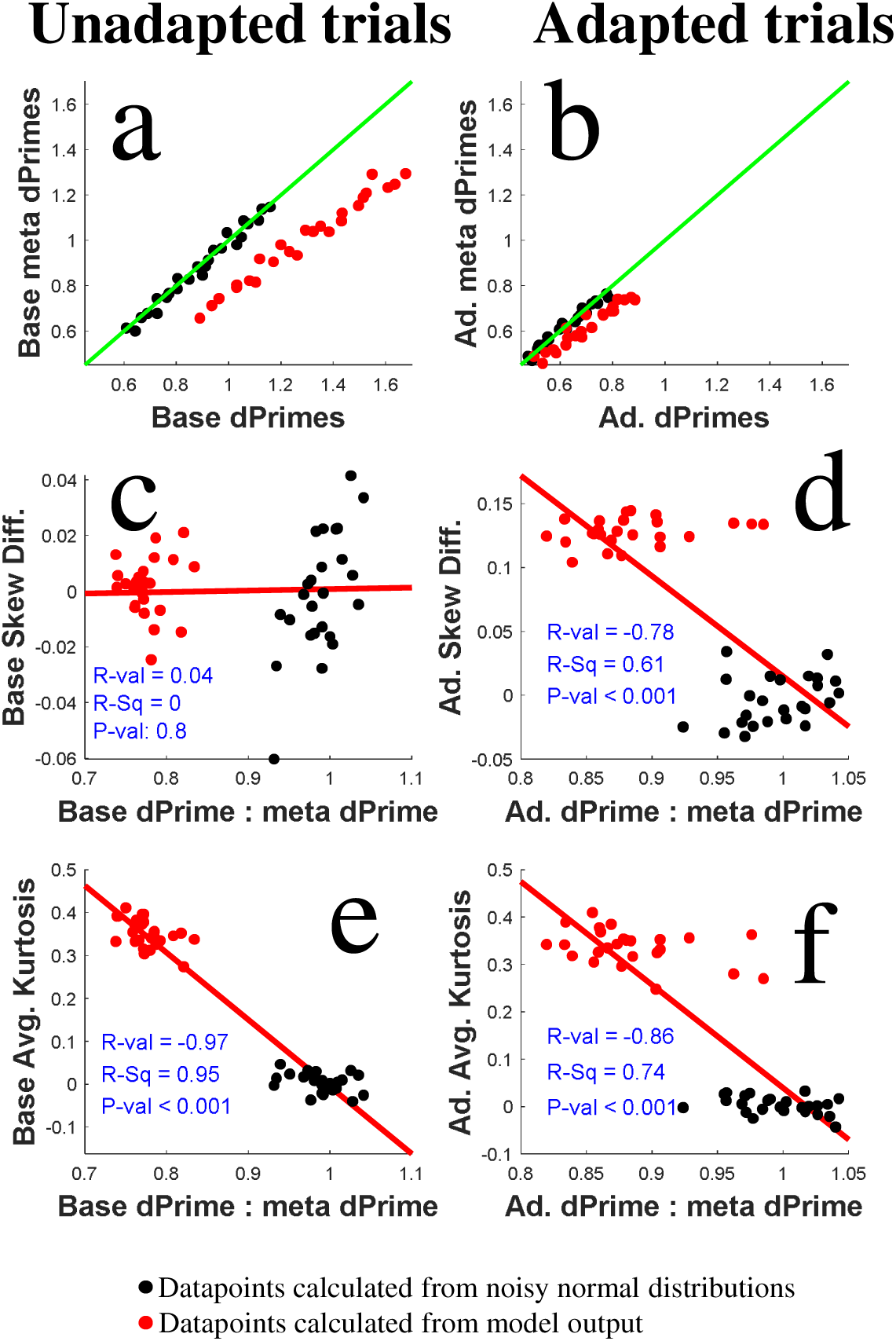
**a)** Scatterplot of d’ (x-axis) and meta d’ (y-axis) estimates calculated from our unadapted model output (red data points) or from values randomly sampled from normally-shaped distributions (black data). For each data point 50000 trials were simulated, 25000 non-target ‘Noise’ and 25000 target ‘Signal’ presentations. All d’ and meta d’ estimates should fall along the horizontal green line if they have been calculated from normally-shaped distributions. **b)** Details are as for Figure 5a, but for (red) data relating to a +40° adapted model. **c)** Scatter plot of Baseline d’ : meta d’ ratios (X-axis) and the average skew of distributions of Baseline model encodings that have informed SDT-based analyses. **d)** Details are as for Figure 5c, but for data relating to +40° adapted models. **e)** Details are as for Figure 5c, but with average baseline distribution excess kurtosis on the Y-axis. **f)** Details are as for Figure 5e, but for data relating to +40° adapted models.

To compare model performance with categorisations based on normally-shaped distributions, we also included conditions where we randomly sample values from normally-shaped distributions. The mean values of these distributions were set to the ‘signal’ and ‘noise’ input values for the relevant simulated experiment (i.e. to one of a range of differences, from +/-1.5 to 2.7° from the direction encoded by the relevant model as upwards). Standard deviations were set to the average JND of our human observers when making perceptual decisions for the same experimental condition (i.e. to 4.5° and to 6.7° respectively for the unadapted Baseline and for the +40° Adaptation conditions).

Results of our second set of simulated experiments are depicted in Figure 5. There are a few important features to note. First, SDT-based analyses informed by both of our model states (Baseline and +40° Adapted) underestimate meta d’ relative to d’ (see red data points in Figure 5a-b, and note that they all fall below the oblique green lines). This would normally be regarded as evidence of metacognitive insensitivity, with confidence judgments assumed to have been impacted by an additional source of noise relative to perceptual decisions (e.g. Fleming & Lau, 2014; Maniscalco & Lau, 2012; Wixted & Stretch, 2004). In our simulated experiments we know this is not true. Second, note that both d’ and meta d’ estimates of sensitivity are reduced for our +40°adapted model, relative to our unadapted Baseline model, capturing the decline in sensitivity displayed by human participants post +40°adaptation (see Figure 3g).

Readers should also note that while SDT-based analyses underestimate meta d’ relative to d’ when analyses are based on model encodings (see red data points, Figure 5a-b), estimates based on values randomly sampled from normal distributions (black data points) cluster about the green line marking a 1:1 ratio for meta d’ : d’ estimates – as predicted by SDT-based analyses of confidence when data are informed by normally-shaped distributions. This can be regarded as a sanity check – signifying that SDT-based analyses of confidence perform as expected when analysis assumptions are met.

To diagnose the cause(s) of the underestimation of meta d’ relative to d’ in analyses informed by our model encodings, we have included scatterplots of meta d’ : d’ ratios (X-axes) and differences in Signal and Noise skews (see Figure 5c-d, Y axes). We have also included scatterplots of meta d’ : d’ ratios (X axes) and excess Kurtosis (averaged across distributions describing both ‘Signal’ and ‘Noise’ test encodings, see Y-axes, Figure 5e-f). For unadapted Baseline model encodings, underestimation of meta d’ relative to d’ is predicted by excess kurtosis – by Signal and Noise distributions having a greater number of extreme encoded values than would be predicted by Normally-shaped distributions (see Figure 5e). There was no discernible relationship with a difference in baseline Signal and Noise distribution skews, as these distributions were effectively un-skewed (i.e. distribution skews were < +/-0.02).

For +40° adapted model encodings, underestimation of meta d’ relative to d’ was again predicted by excess kurtosis (see Figure 5f). However, for +40° adapted model encodings there was an additional robust relationship between underestimations of meta d’ relative to d’ and a difference between Signal and Noise distribution skews (see Figure 5d).

Our analyses suggest that SDT-based analyses of confidence can be undermined by non-normal experiential distributions. A reasonable question would then be whether this observation might matter in practice. To assess this, for our Baseline and +40° adapted model we plot meta d’ : d’ ratios as a function of the magnitude of input differences (see Figure 6). Estimates informed by analyses of values randomly sampled from normally-shaped distributions (black data points) fall within green shaded regions, which mark a ratio of 1:1 +/-5%. Note, however, that ratio estimates informed by values encoded by our two models fall below this region, with meta d’ estimates under-estimated by ∼25% (Baseline model) and by ∼15% (+40° adapted model).

**Figure 6.**
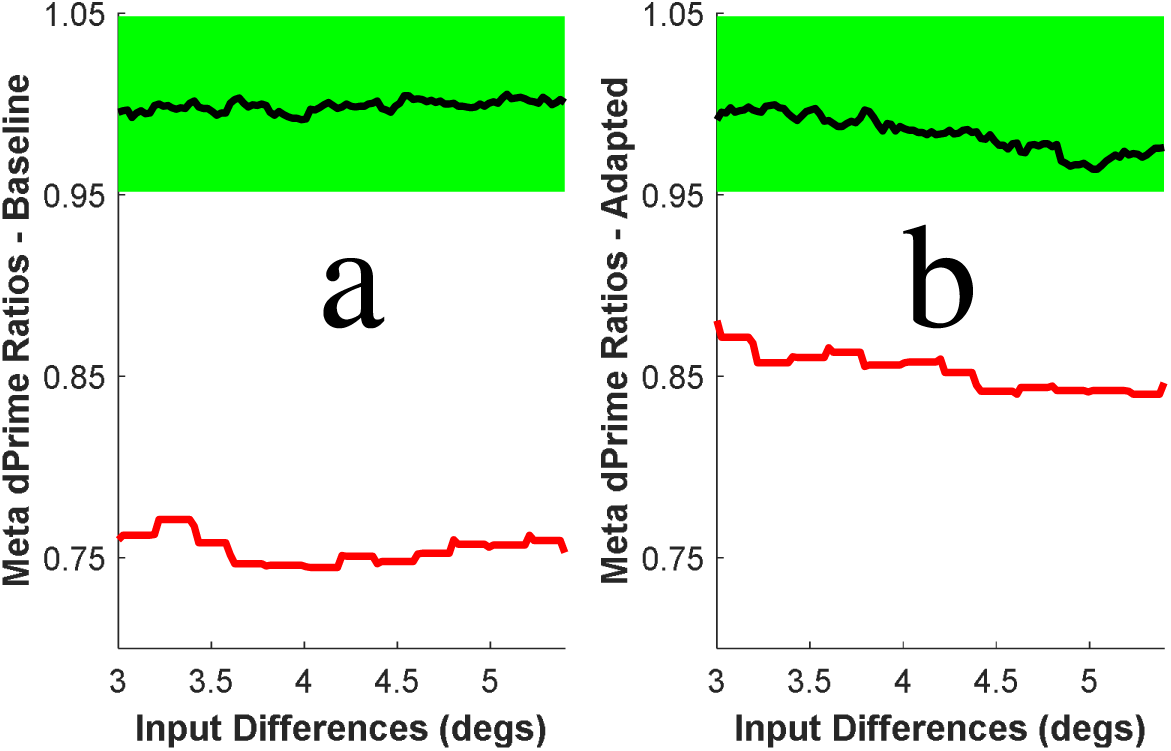
Meta d’ and d’ estimates, expressed as ratios for unadapted Baseline (**a**) and +40° adapted (**b**) conditions, as a function of ‘Signal’ and ‘Noise’ input differences. Data are depicted for analyses based on values randomly sampled from normally-shaped distributions (black data) and from model encodings (red data). Green shaded regions mark +/-5% from a ratio of 1.

Our results to this point can collectively be regarded as an existence proof, that if SDT-based analyses of confidence are applied to datasets that have been informed by non-normal experiential distributions (by Signal and Noise distributions that are either differently skewed, or have excess kurtosis, or both), results can encourage flawed interpretations. How could researchers avoid drawing a false conclusion, that confidence has probably been adversely impacted by some additional source of noise relative to perceptual decisions (e.g. Fleming & Lau, 2014; Maniscalco & Lau, 2012; Wixted & Stretch, 2004), when data has instead been informed by a common source of information characterised by an abnormal shape?

Researchers often do not check for violations of the normality assumption when conducting SDT-based analyses. When they do, the most common check is to plot z-scored hit and false alarm rates, which can be done for target and noise inputs that elicit different levels of performance (e.g. Pastore & Scheirer, 1974) or for inputs that elicit different levels of confidence (e.g. Wixted & Stretch, 2004). When plotted, these should fall along a line with a slope of 1 if they have been calculated from normally-shaped experiential distributions with equal variance (Green & Swets, 1966). In our next set of simulations, we ask if researchers could reliably distinguish between datasets that have been informed by normally- and by non-normal distributions based on the slope associated with X/Y plots of z-scored hit and false alarm rates.

For this set of analyses, we conducted 1000 simulations wherein we randomly select 5 magnitudes of ‘Signal’ and ‘Noise’ test difference (from the 101 levels sampled in our second set of simulated experiments). For each condition we randomly select half the model encodings / values randomly sampled from normal distributions (i.e. data from 30000 simulated trials). From these data, we calculate best-fit linear slopes for sets of 3, 4 or 5 z-scored hit and false alarm rates plotted on an X/Y scatterplot. From these, we calculate threshold criteria, for rejecting slopes as having likely resulted from analyses informed by non-normal distributions (i.e. +/-3 S.D.s from the average slope associated with z-scored hit and false alarm rates for values randomly sampled from normal distributions).

Bar plots depicting proportions of failed ‘slope tests’ are depicted in Figure 7. Note that by definition only a small number of slope tests are ‘failed’ when these criteria are applied to analyses informed by values randomly sampled from normal distributions (see Figure 7a and 7c). This is unsurprising, as these analyses are inherently circular (the test rejection criteria have been calculated from these same data). The point of these analyses is to calculate rejection criterion values that can be applied to slopes calculated for z-scored model encodings. For our Baseline model (which underestimated meta d’ by ∼25%) this resulted in encouraging rejection rates (of ∼95%). However, for our +40° adapted model (which underestimated meta d’ by ∼15%) this resulted in far smaller rejection rates (of ∼5%). This means that experiential distributions that are sufficiently non-normal in order to undermine SDT-based analyses of confidence could not be detected on ∼95% of simulations on the basis of a z-scored slope tests, even when we had detailed knowledge of what tolerance we should adopt so as not to dismiss analyses that were actually informed by normally-shaped distributions (i.e. the average slope +/-3 S.D.s, calculated from 1000 fits to pairs of 3, 4 or 5 z-scored Hit and False Alarm rates calculated from values randomly sampled from normal distributions).

**Figure 7.**
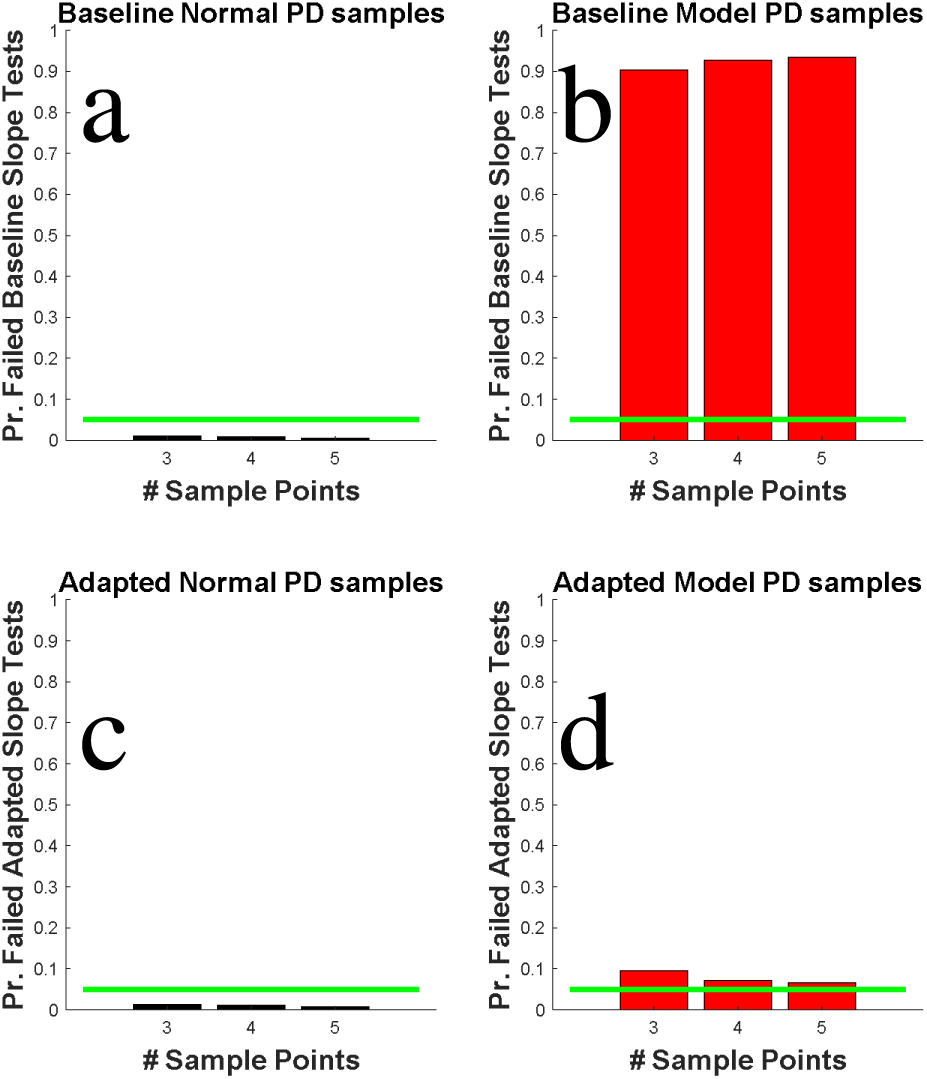
**a)** Bar plot depicting the proportion of 1000 simulations where fitted slopes were > +/-3 S.D.s from the average slope fit to z-scored Hit and False Alarm rates, calculated for random samples taken from normally-shaped distributions (i.e. failed slope tests). These data are circular (as average values and S.D.s were calculated from these data), so by definition there are only a small proportion of failed tests. **b)** Details are as for Figure 7a, but for slopes fit to z-scored Hit and False Alarm rates calculated from encodings by our unadapted Baseline model. **c)** Details are as for Figure 7a, but for data calculated for random samples taken from normally-shaped distributions with mean values and S.D.s set to mimic the performance of our human participants when adapted to +40°. **d)** Details are as for Figure 7b, but for data calculated from encodings by our +40° adapted model.

### Stage 4: What impact do different levels of distribution skew and excess kurtosis have on SDT-based analyses of confidence?

To this point our simulated analyses, describing the impact of non-normal distributions on SDT-based analyses of confidence, have related to analyses of model-generated encodings. This has a benefit, in that our model provides a good approximation of human performance in an actual experiment (see Figure 3), so the degrees of post-adaptation skew, and pre- and post-adaptation kurtosis, suggested by our model might resemble those produced by operations of the human mind. However, this treatment limits analyses to 2 model states (Baseline and +40° adapted), that are likely characterised by a singular differential skew and average level of excess kurtosis. Our analyses to this point are also tied to the performance of our model, so it seems worth making a more general point using analyses that do not depend on the operations of our model.

We isolate and depict the effects of excess kurtosis and differential skew across a range of plausible values in a final set of simulations. For these, we selected the maximal average kurtosis produced by our model (∼0.4) and the maximal level of a distributional skew (∼0.16). To describe the effects of excess kurtosis, we use the Matlab pearsrnd command to create 51 pairs of distributions of signal and noise values with a common average offset (+/-2° from a central perceptual category boundary of 0°), and a common standard deviation (of 3°), but with different levels of excess kurtosis – ranging from 0 to 0.4 (i.e. levels of kurtosis range from 3, which describes a normal distribution, to 3.4). We use a confidence criterion value of 7 to classify generated values as noise presentations with high or low confidence, or as signal presentations with low or high confidence – and we estimate both d’ and meta d’ (code for this analysis is provided as Supplemental material code #3). As can be seen in Figure 8a, as excess kurtosis increases, so too does the underestimation of meta d’ relative to

We conducted a final set of analyses, to describe the impact of differential signal and noise distribution skews. These were similar to our simulations describing the impact of excess kurtosis – but for these simulations we used the Matlab pearsrnd command to set excess kurtosis to 0 and Noise distributions to skews ranging from +0.16 to -0.16, and Signal distribution skews to a range from -0.16 to +0.16. As depicted in Figure 8a, when noise distributions are positively skewed and signal distributions are negatively skewed, meta d’ tends to be overestimated relative to d’, and when noise distributions are negatively skewed and signal distributions are positively skewed, meta d’ tends to be underestimated relative to d’.

## Discussion

Our behavioural experimental data describe how direction perception, the precision of perceptual decisions, and confidence (i.e. the spread of uncertainty) are all impacted by adaptation to different direction differences (Stage 1 of our study; see Figure 3f-g datapoints and error bars). The key qualitative features of these changes could all be explained by a labelled-line observer model, which describes how mappings between inputs, direction perception and confidence might be changed by adaptation (Stage 2 of our study, see Figure 3f-g smoothed lines). We reasoned that the success of this approach could be challenging for SDT-based analyses of confidence, as these assume that repeated exposures to an input should elicit a normally-shaped distribution of different perceptual experiences (Green & Swets, 1966; Fleming & Lau, 2014; Maniscalco & Lau, 2012; Yarrow et al., 2011), whereas visual adaptation could encourage abnormally-shaped distributions.

To assess the possibility of our model might not encode normally-shaped distributions, in Stage 3 we conducted simulated experiments, to find out what shapes would describe the distributions of values encoded by our model for repeated Signal and Noise inputs. We found that these had excess kurtosis both before and after adaptation (i.e. had more extreme values, relative to the predictions of a normal distribution), and were differently skewed post-adaptation (see Figure 5). Moreover, SDT-based analyses of confidence informed by these distributions were undermined – with meta d’ tending to be systematically underestimated relative to d’ – even though these two summary statistics were calculated from a common dataset (see Figure 5).

In Stage 4 of our study, we conducted simulations to examine the impact of distribution skews and excess kurtosis on SDT-based analyses of confidence per se – independent of the operations of our model (see Figure 9). For these simulations we used simple matlab routines to generate data that conformed to differently shaped distributions, and quantified the effect these had on SDT-based analyses of confidence, which commit to the normality assumption (Fleming & Lau, 2014; Maniscalco & Lau, 2012). We established that these analyses are systematically undermined by either type of deviance from the normality assumption, with meta d’ systematically over (when Signal distributions were positively skewed relatively to Noise distributions; see Figure 9b) and underestimated (when Signal distributions were negatively skewed relative to Noise distributions, and when distributions had excess kurtosis; see Figure 9a-b) relative to estimates of d’ – even though both estimates had been informed by a common dataset. Miyoshi et al (2022) have also recently demonstrated this adverse contingency, between excess kurtosis and the performance of SDT-based analyses of confidence. While our analyses were inspired by the first three stages of our study, this particular set of results does not rest on the veracity of the earlier stages of our study. The contingencies we have demonstrated in Stage 4 are factual, regardless of the motivations that led us to test for these.

**Figure 9.**
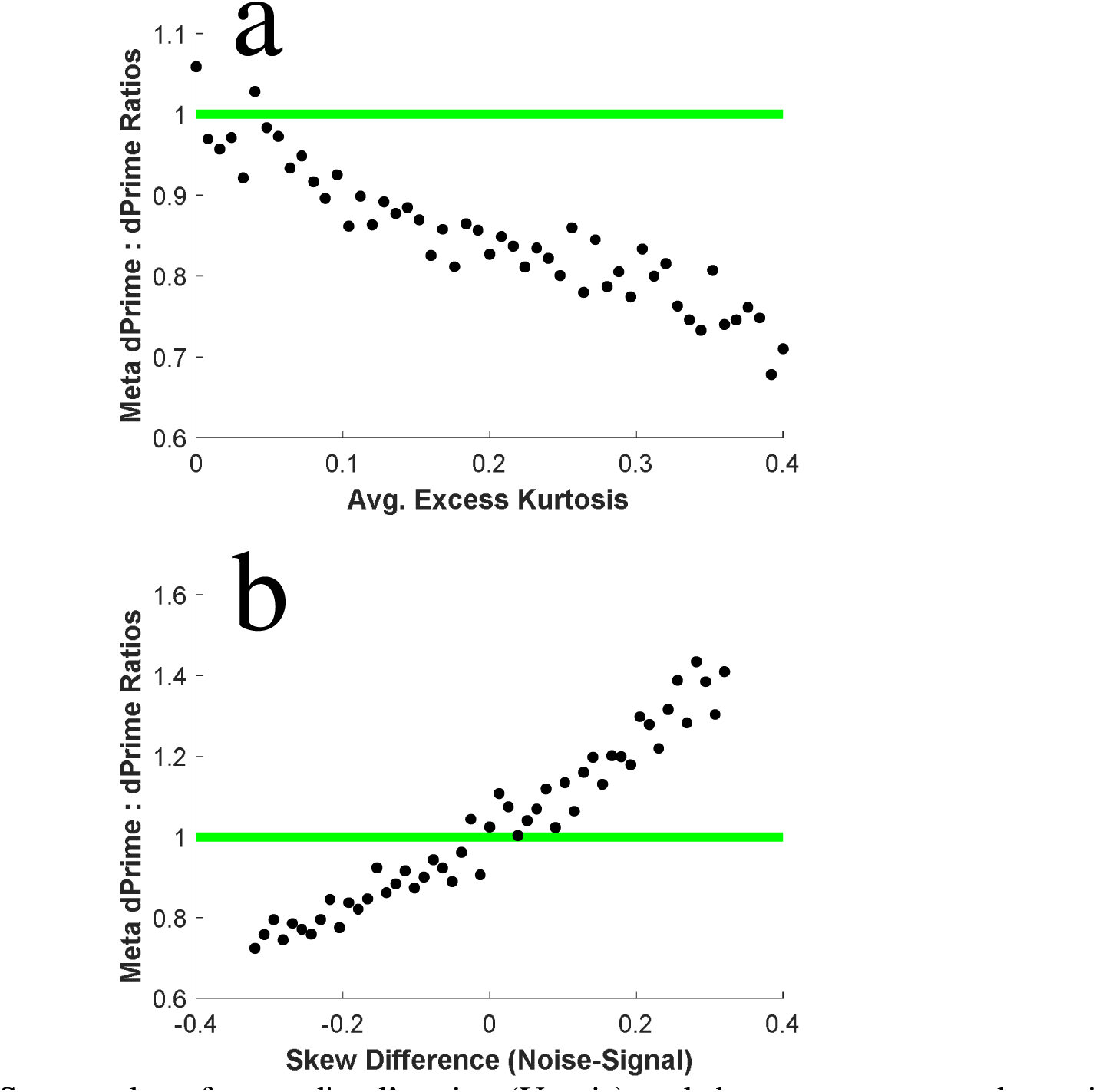
**a)** Scatter plot of meta d’ : d’ ratios (Y-axis) and the average excess kurtosis of signal and noise distributions (X-axis). As kurtosis levels increase from the level of kurtosis that is consistent with a normal distribution (i.e. 3), Meta d’ estimates decrease as a ratio relative to d’ estimates. **b)** Scatter plot of meta d’ : d’ ratios (Y-axis) and the differences in the skew of signal and noise distributions (X-axis). Meta d’ estimates transition from exceeding d’ estimates when Noise distributions are positively skewed and Signal distributions are negatively skewed, to being reduced relative to d’ estimates when Noise distributions are negatively skewed and Signal distributions are positively skewed.

We regard our data as an existence proof – that if SDT-based analyses of confidence are inadvertently applied to data that are informed by abnormally-shaped experiential distributions (that either have excess kurtosis, are skewed, or both), results can encourage flawed conclusions. In our simulated experiments analyses of confidence and perception were informed by the same data – a situation that is usually understood to be metacognitively ideal (Fleming & Lau, 2014; Maniscalco & Lau, 2012). Our results, however, show that confidence analyses informed by these data can result in smaller meta d’ than d’ estimates (see Figures 5 and 10), a situation that is usually regarded as evidence that a participant is metacognitively inefficient, and that their estimates of confidence must be subject to an additional source of noise relative to perceptual judgments (Fleming & Lau, 2014; Maniscalco & Lau, 2012). Our analyses show that this need not be true – that meta d’ and d’ differences can instead result from applying analyses that assume normally-shaped experiential distributions to data that deviate from normality.

### Our observer model is an unrealistic oversimplification – with some redeeming features

A reasonable criticism of our study is that our observer model is an oversimplification, that does not capture key features of the decisional processes related to confidence. One glaring example of this is that we have made the simplifying assumption that all noise within the model resides within encoding stages – as opposed to the criteria used when categorizing encoded values for perceptual decisions and confidence ratings. Other studies have enjoyed considerable success in explaining human behaviour, in studies of confidence, by describing how confidence criteria might be subject to noise (e.g. Adler & Ma, 2018; Desender et al., 2022; Mamassian & de Gardelle, 2021; Miyoshi & Lau, 2020; Shekhar & Rahnev, 2021). Indeed, two of the authors have previously highlighted, in the context of time perception, that effects attributed to encoding changes can often equally, or better be explained by changes to decisional criteria (Yarrow et al., 2011). Our choice here, to assume a noiseless application of perceptual and confidence criteria, was not made because we think these are realistic or inconsequential assumptions, but rather because these are assumptions underlying popular SDT-based analyses of confidence (Fleming & Lau, 2014; Maniscalco & Lau, 2012) – and we ultimately wanted to subject our model data to this form of analysis.

While limited, our model has served as motivation to examine how SDT-based analyses of confidence perform when informed by experiential distributions that are skewed or have excess kurtosis. Our model also demonstrates that an experimental manipulation (in this case adaptation) can have a disproportionate impact on summary statistics describing confidence (in this case FWHH aftereffects, see red data Figure 3g) relative to a summary statistic describing the precision of perceptual decisions (JND aftereffects, see blue data Figure 3g) – merely because these two summary statistics rely on different magnitudes of the same type of information (also see Arnold et al., 2021). This highlights that apparent dissociations between measures of confidence and perceptual precision are possible, even when these map onto the same type of information.

### What does this mean for SDT-based analyses of confidence?

Our data point to a dilemma – even in a context where we know that analyses of confidence have been informed by non-normal distributions, that fact could not reliably be detected by the standard procedure experimenters rely on to assess if a SDT-based analysis might have been informed by non-normal experiential distributions. This is to check for deviance from a 1:1 slope for z-scored hit and false alarm rates (Green & Swets, 1966; Pastore & Scheirer, 1974; Wixted & Stretch, 2004). This shows that when a SDT-based analysis of confidence suggests a participant is metacognitively inefficient, there will be a degree of ambiguity – that rather than inefficiency, the meta d’ - d’ difference might instead be driven by analyses having been inadvertently informed by non-normal experiential distributions.

While our data serve as an existence proof, of what can happen when SDT-based analyses of confidence are informed by non-normal experiential distributions (also see Miyoshi et al, 2022), the distributions that informed the analyses of Stages 3 and 4 of this study were computer generated – not a product of a human mind. Researchers do not have ready access to the experiential distributions that inform our internal decision processes. These can be estimated, from experiments involving a great number of reproductions of an input by human participants (e.g. Bae et al., 2015; Bobko et al., 1977; de Gardelle et al., 2010), but these reproductions could be contaminated by noise that is additional to perception (i.e. by short term memory noise, e.g. Bays et al., 2011; Yarrow et al., 2020) and possibly by reporting biases, such as a tendency to reproduce inputs as an exemplar experience (see Bae et al., 2015), or a tendency to exaggerate differences between a perceived input and exemplar experiences (see de Gardelle et al., 2010). With these reservations in mind, there is evidence that human experiential distributions, for tilt and time, might be subject to excess kurtosis (Acerbi et al., 2012; Anderson, 2014; Jabar & Anderson, 2015).

While researchers usually do not have ready access to a ground truth estimate of the shape of the experiential distributions they are investigating, these are assumed to have a normal shape (Green & Swets, 1966; Fleming & Lau, 2014; Maniscalco & Lau, 2012; Yarrow et al., 2011). If experiential distributions were normally-shaped in fact, our demonstration that SDT-based analyses of confidence can be undermined if they are applied to non-normal distributions would have no consequence. So, it is reasonable to ask if there is any evidence that experiential distributions might be abnormally shaped? We would argue yes.

### Evidence that experiential distributions might be abnormally shaped

It seems reasonable to presume that measures of perceptual sensitivity and precision will inversely scale with the magnitude of trial-by-trial changes in how inputs are encoded and experienced. Across many visual dimensions, sensitivity and precision are not uniform. Humans are more sensitive to direction and to orientation differences about cardinal (vertical and horizontal) as opposed to oblique angles (Appelle, 1972; Dakin et al., 2005; Girshick et al., 2011; Storrs & Arnold, 2015). Spatial acuity scales with distance from fixation (Virsu & Rovamo, 1979), and people are more sensitive to differences between slower speeds than to differences between faster speeds (e.g. Stocker & Simoncelli, 2006). In each of these cases, experiences triggered by repeated exposures to a given input might be associated with a greater range of experiences extending into regions of the perceptual dimension marked by less sensitivity/precision, and with a smaller range of experiences extending into regions associated with higher sensitivity/precision – in sum producing skewed experiential distributions. In human vision, skewed experiential distributions might be more the rule than an exception.

We believe there is also good evidence to suggest that human experiential distributions, from repeated exposures to a common input, might be characterized by excess kurtosis. We have already mentioned evidence from reproduction experiments, suggesting human experiential distributions might be characterized by excess kurtosis (Acerbi et al., 2012; Anderson, 2014; Bays, 2016; Jabar & Anderson, 2015). In addition, the distribution of responses amongst cells tuned to different instances of a perceptual dimension in vision (i.e. to different orientations) is often marked by excess kurtosis – particularly for sections of a dimension that have a learnt relevance (e.g. Failor et al., 2021) and when inputs are natural images (e.g. Field, 1987; Olshausen and Field, 1996).

### Consequences of performing SDT-based analyses of confidence that assume normality on abnormally-shaped experiential distributions

Our analyses show that if SDT-based analyses of confidence are inadvertently informed by differentially skewed experiential distributions, meta d’ can either be underestimated relative to d’ (when Signal distributions are more Left-skewed) or overestimated (when Signal distributions are more Right-skewed, see Figure 8b). Results characterized by greater meta d’ than d’ estimates have been a source of some speculation as to probable cause (e.g. Rahnev & Fleming, 2019). Our analyses reveal one cause of this could be that SDT-based analyses of confidence have inadvertently been performed on data informed by skewed experiential distributions (see Figure 8b).

We have also shown that if SDT-based analyses of confidence are inadvertently conducted on experiential distributions that have excess kurtosis, meta d’ will be systematically underestimated relative to d’ (see Figure 8a), and it is likely this could happen without the experimenter being able to detect evidence of the non-normal distributions (see Figure 8).

This is particularly troubling, as the dominant finding of studies that have used SDT-based analyses of confidence is that estimates of meta d’ are less than estimates of d’, and this has been interpreted as evidence that confidence judgments are subject to an additional source of noise (Fleming & Lau, 2014; Maniscalco & Lau, 2012). Our analyses reveal that this conclusion need not follow from this type of finding. We suspect that there will always be some ambiguity as to the underlying cause for this type of finding.

### SDT does not need to assume Gaussian-shaped experiential distributions – but assuming some other shape might not solve the problem

While popular instantiations of SDT commit to the normality assumption, including popular SDT-based analyses of confidence (e.g. Fleming & Lau, 2014; Maniscalco & Lau, 2012), SDT is overall agnostic about the precise shape of experiential distributions. Provided a particular shape of distribution can be assumed, areal overlaps can be estimated from behavioural data – it does not matter if the distributions are assumed to have a normal or an abnormal shape (Green & Swets, 1966). Indeed, some researchers have advocated for performing SDT-based analyses of confidence using calculations that assume non-normal experiential distributions (e.g. Medha & Dobromir, 2021; Miyoshi et al., 2022; Winter & Peters, 2022), or which have a differently shaped distribution to describe confidence (Boundy-Singer et al., 2022) or the variance of confidence criteria (Shekhar & Rahnev, 2021). The problem with this approach is that the criticisms we have made about the impact of erroneously assuming a normally-shaped experiential distribution could apply equally to erroneously assuming any other distribution shape, and we doubt the precise shape of distributions will ever be known in sufficient detail to preclude the possibility that results might differ from ground truth due to the wrong shape(s) having been assumed in calculations. Using this general approach, we believe there will always be some doubt as to whether evidence has revealed that confidence is truly subject to additional noise, or if the wrong shaped distributions have been assumed.

### Why do abnormally-shaped distributions have a disproportionate impact on SDT-based analyses of confidence? The problem is with the tails

A lot of our analyses have been based on calculations that assume normally-shaped distributions. For instance, our modelling involved fitting cumulative Gaussian functions to modelled categorical perceptual decisions (see Figure 4b). As can be seen, this approach can often result in a very good description of data – even in this context where we know the distributions have a shape that is sufficiently abnormal to undermine SDT-based analyses of confidence (see Figures 5 – 6).

The problem for popular SDT-based analyses of confidence (Fleming & Lau, 2014; Maniscalco & Lau, 2012) is that these are exclusively based on trials that are informed by data that would reside within the far right-side tails of experiential distributions – categorical perceptual decisions that have resulted in high-confidence target categorizations (see Figure 1b). This emphasis makes these analyses particularly susceptible to the impact of any deviance from a normally-shaped distribution, as the impact of an abnormal shape, in terms of areal overlap, is exaggerated at the tails relative to the entirety of a distribution.

One consequence of the over reliance of popular SDT-based analyses of confidence on the extremes of experiential distributions is that the ramifications of our analyses might be specific to these popular SDT-based analyses of confidence (Fleming & Lau, 2014; Maniscalco & Lau, 2012). Our data are less challenging to standard implementations of the SDT framework (Green & Swets, 1966) that use entire datasets when trying to infer areal overlaps between experiential distributions.

### What’s the solution?

Researchers should be aware that popular SDT-based analyses of confidence (Fleming & Lau, 2014; Maniscalco & Lau, 2012) do not provide a ground truth measure of human metacognitive sensitivity. They provide an approximation of what this might be, accompanied by some degree of unavoidable ambiguity, as to whether metacognitive sensitivity might really have differed, or if analyses might have inadvertently been informed by experiential distributions that are abnormally shaped.

To avoid interpretive ambiguity, estimates of metacognitive sensitivity should not be disproportionally impacted by any inadvertent difference between the assumed and actual shapes of experiential distributions. A possible candidate is a recent modelling approach outlined by Mamassian & de Gardelle (2021). While this approach is informed by the SDT framework, it involves a modelling of the entirety of psychometric functions, and so it may be less susceptible to the influences we have identified. This, however, will need to be formally evaluated.

Until a better means of estimating metacognitive sensitivity can be developed and proven, researchers are urged to acknowledge ambiguities when interpreting the results of perceptual confidence experiments, in order to avoid over interpretations of data.

## Supporting information

Supplemental Matlab files

## Acknowledgements

This research was supported by a Discovery Project Grant DP200102227, funded by the Australian Research Council, awarded to D.H.A. & A.J.

## Conflict of interest

The authors declare no competing financial interests.

## Data and materials availability

All data and analysis scripts for this project will be made available via UQeSpace https://espace.library.uq.edu.au

## Author contributions

D.H.A & A.J. conceived of the study. D.H.A. programmed experiments, analysed data and wrote the first draft of the manuscript. J.A. tested participants. All authors edited successive versions of the manuscript.

## Funding

This research was supported by an ARC Discovery Project Grant awarded to DHA and AJ.

## Notes

### Competing Interest Statement

The authors have declared no competing interest.

