## Supplemental Matlab files for "On why we lack confidence in signal-detection-based analyses of confidence": Supp_1_MAE_Confidence_Population_Code.m

```
%% This code implements a labelled-line observer model, which runs 
% simulated experiments that mimic the effects of adapting to different 
% directions of motion, and determines PSE estimates calculated from 
% perceptual decisions and the peak of raised Gaussian distributions fit to 
% uncertainty (low-confidence decisions) both as a function of test  
% direction. The model also determines JND estimates for perceptual  
% decisions, and FWHH estimates associated with distributions of  
% low-confidence - and calculates aftereffects for all of these estimates  
% relative to baseline (unadapted model) encodings. Trial-by-trial model 
% encoding noise is implemented by randomly modulating each model channel's 
% response potential (between 0 and 1). Model performance is compared to  
% functions collected for human participants. 

clear all % Clear memory
close all % close all figures

% If psignifit has not already been added to your path, do that now
addpath /Users/uqdarnol/Dropbox/Research_Projects/Toolbox/Psignifit/psignifit-master

%% Model parameters

Channel_Spacing = 10;       % Spacing between peak tunings of model channels

Gain_Factor = 1.8;          % Controls the spread of reductions in channel gain potentials post-adaptation
Shift_Factor = 4.5;         % Controls the spread of channel tuning shifts post-adaptation

Ad_Amplitude = 0.95;        % The maximal amount of post-adaptation channel gain reductions 
Shift_Amplitude = 40;       % The maximal post-adaptation channel tuning shift

Confidence_Criterion = 7;   % The decoded angular difference beyond the category 
                            % criterion (0) needed to evoke a 'high' 
                            % confidence response
                            
Attenuate_0_Adaptor = 0.5;  % Amount to reduce adaptation by when you have adapted to the average test value

sigma = 20;                 % The S.D. of model channels 

%% Pre-Generate adaptation induced channel gain reductions and tuning shifts

Xs = -180:Channel_Spacing:180-Channel_Spacing; % An array of direction values, for use in calculating adaptation multiples

% Pre-determine adaptation induced changes
for n = 1 : size(Xs,2)
    
    if abs(Xs(n)*Gain_Factor)<=180
        % Multiple to reduce channel gain by, determined by angular
        % difference from the adaptor (0 degrees)
        Post_Ad_Gain_Reductions(n)=cosd(abs(Xs(n)*Gain_Factor)/2);
    else
        Post_Ad_Gain_Reductions(n)=0;
    end
    
    if sind(Xs(n)*Shift_Factor) < 0 & abs(Xs(n)*Shift_Factor)<=180
        % Multiple to shift channel tuning by, for negative angular
        % differences from the adaptor (0 degrees, so a repulsive tuning
        % shift)
        Tuning_Shift_Multiples(n)=-sind(abs(Xs(n)*Shift_Factor)/1);
    elseif sind(Xs(n)*Shift_Factor) > 0 & abs(Xs(n)*Shift_Factor)<=180
        % Multiple to shift channel tuning by, for positive angular
        % differences from the adaptor (0 degrees, so a repulsive tuning
        % shift)
        Tuning_Shift_Multiples(n)=sind(abs(Xs(n)*Shift_Factor)/1);
    else
        Tuning_Shift_Multiples(n) = 0;
    end
    
end

%% Parameters for calculating an unadapted population code

x = -180:.1:180; % Create an array of x indicies, finely sampling the full gamut of possible directions
s = size(x,2);  % Record the size of the X axis indices array

Population_N = size(Tuning_Shift_Multiples,2); % Number of channels in population code
end1 = -180; % Specify the tuning of the first channel
end2 = 180-Channel_Spacing;  % Specify the tuning of the last channel
Chn_Labels=end1:((end2-end1)/(Population_N-1)):end2; % Determine ALL channel labels / tunings 
                                                     % (i.e. what value they are voting for when active)

%% Simulate Experiments

% Specify what values the model will adapt to
Adaptors = 0:Channel_Spacing:180;

% Create an array of test direction values
T1s= -28.5:4:-12.5;     % Begin with large negative values and step size
T2s= -11.5:2:-3.5;      % More finely sampled negative test values
T3s = -2.5:0.5:2.5;     % Even more finely sampled negative/positive test values
T4s = 3.5:2:11.5;       % Finely sampled positive test values
T5s = 12.5:4:28.5;      % Large positive test values and big step size
Test_Vals = cat(2,T1s,T2s,T3s,T4s,T5s); % Here we combine the five parts of the test array

N_Repeats = 500;        % Specify the number of times each test value will 
                        % be sampled. As the computer will not complain, 
                        % simulate A LOT of trials, to get an accuarate 
                        % estimate of modelled psychometric functions
                        
Tests = []; % Create an array that we will store a randomised order of test values in
for r = 1 : N_Repeats 
    Tests(:,r) = Test_Vals(randperm(length(Test_Vals))); % Store a random permutation of the test values in the test array
end
Tests = reshape(Tests,1,size(Tests,1)*size(Tests,2)); % reshape the test array into a 1 x 1320 sized array


Decoded_Vals=[]; % Create an array to store values decoded from unadapted population code
Adapted_Decoded_Vals = []; % Create a second array to store values decoded from adapted population codes

Baseline_Categorisation_Array = zeros(size(Test_Vals,2),3); % Create a 27 x 3 column array, to store unadapted 
Baseline_Categorisation_Array(:,1)=Test_Vals;               % categorisations (i.e. tilted left / right of vertical).
                                                            % First column will store test values, second number of
                                                            % tilted right categorisations, third number of times that
                                                            % test value was sampled
                                                            
% Create a matching array to store adapted categorisation data
Adapted_Categorisation_Array = zeros(size(Test_Vals,2),3,size(Adaptors,2));
for n = 1 : size(Adaptors,2)
    Adapted_Categorisation_Array(:,1,n)=Test_Vals';
end

% Create matching arrays to store confidence data in
Baseline_Confidence_Array = Baseline_Categorisation_Array;
Adapted_Confidence_Array = Adapted_Categorisation_Array;    


%% Simulated experiment

close all    
% Begin simulating experiment... 
for Trial = 1 : size(Tests,2) 
    
    % Loop to create baseline channels for population code
    for Ch_Num = 1 : Population_N 
        
        % If we are simulating trial-by-trial encoding noise generate a
        % random gain multiple for each model channel
        Channel_Tuning_Noise(Ch_Num) = 1;%rand(1);
        
        % Generate baseline model channels - normal probability density
        % functions, centered on different physical test values
        % (Chn_Labels) with a standard deviation (sigma)
        Trial_Baseline_Channels(Ch_Num,:) = normpdf(x,Chn_Labels(Ch_Num),sigma);
        
        % Make sure that the baseline channels start scaled 0 to 1
        Trial_Baseline_Channels(Ch_Num,:) = Trial_Baseline_Channels(Ch_Num,:)./max(Trial_Baseline_Channels(Ch_Num,:));
        
        % Now multiply the channel's gain potential by the encoding noise 
        % level set for this channel on this trial...
        Trial_Baseline_Channels(Ch_Num,:) = Trial_Baseline_Channels(Ch_Num,:)*(Channel_Tuning_Noise(Ch_Num));
        
    end

    % Find the position on X axis that corresponds with the Test for this trial...
    Test_idx = find(x==Tests(Trial));

    % Determine what row of the results arrays to store data in...
    [col Results_Row] = find(Test_Vals==Tests(Trial));

    Votes = [];     % Prepare an empty array to store channel votes in
    Vote_Labels = Chn_Labels+180;
    
    % Calculate model encoding, by taking a weighted sum of channel 'votes'
    % (e.g. channel responses to the test). Channel labels are multiplied 
    % by channel response magnitudes (which are divided by the total
    % response of ALL channels)
    Votes(1:Population_N) = Vote_Labels(:).*((Trial_Baseline_Channels(:,Test_idx))./sum(Trial_Baseline_Channels(:,Test_idx)));

    % The decoded input value is the weighted sum of all the channel
    % 'votes'
    Decoded_Vals(Trial) = sum(Votes)-180; 
    
    % Categorise input as Negative or Positive - by recording when trials
    % have been encoded as a positive value
    if Decoded_Vals(Trial) >= 0
        Baseline_Categorisation_Array(Results_Row,2)=Baseline_Categorisation_Array(Results_Row,2)+1;
        Category_Resp = 1;
    else
        Category_Resp = 0;
    end
    % Record the number of times that a test value has been sampled
    Baseline_Categorisation_Array(Results_Row,3)=Baseline_Categorisation_Array(Results_Row,3)+1;
    

    % Categorise confidence as low or high, relative to the confidence
    % criterion (the unsigned encoded value needed to elicit a high-level of 
    % confidence) by recording when trials have exceeded the criterion 
    if abs(Decoded_Vals(Trial)) >= Confidence_Criterion
        Baseline_Confidence_Array(Results_Row,2)=Baseline_Confidence_Array(Results_Row,2)+1;
        Confidence_Resp = 1;
    else
        Confidence_Resp = 0;
    end
    Baseline_Confidence_Array(Results_Row,3)=Baseline_Confidence_Array(Results_Row,3)+1;
    

    % Loop to simulate the consequences of adaptation. 1 loop per adaptor
    for Ad_Num = 1 : size(Adaptors,2)
        
        Adapted_Labels=Chn_Labels; % Initial labels for each channel

        % Identify which channel is tuned to 0 - which we have used as the
        % 'adapted' channel...
        Ad_Idx = find(Xs==min(abs(Xs)))-1;
        
        % Now we create an array of indices that we use to determine how
        % much to reduce the response potential of each model channel, and
        % how much (and in which direction) to shift each model channels
        % tuning by for this adaptor...
        Ad_Index(1:Population_N) =  1 + mod( Ad_Idx+floor((Chn_Labels(1:Population_N)-Adaptors(Ad_Num))/Channel_Spacing) , Population_N);
        
        % Reduce this channel's response potential by the reduction gain
        % for this channel after adapting to this adaptor...
        Adapt_Mag(1:Population_N) = Post_Ad_Gain_Reductions(Ad_Index);
        
        % Reduce the impact of adaptation after adapting to the average
        % test value, in recognition of the fact that adaptation is
        % cumulative across trials, and consequently will be reduced if the 
        % adaptor is the same as the average test value... 
        % see... 
        % Á Kristjánsson (2011). The functional benefits of tilt adaptation. 
        % Seeing and perceiving 24 (1), 37-51.
        % (and)
        % Fischer & Whitney (2014). Serial dependence in visual perception.
        % Nature Neuroscience 17, 738 - 743.         
        if Ad_Num == 1
            Shift_Mag(1:Population_N) = (Tuning_Shift_Multiples(Ad_Index)*Shift_Amplitude*(1-Attenuate_0_Adaptor));
        elseif Ad_Num == 2 | Ad_Num == size(Adaptors,2)
            Shift_Mag(1:Population_N) = (Tuning_Shift_Multiples(Ad_Index)*Shift_Amplitude*(1-Attenuate_0_Adaptor*0.75));
        elseif Ad_Num == 3 | Ad_Num == size(Adaptors,2)-1
            Shift_Mag(1:Population_N) = (Tuning_Shift_Multiples(Ad_Index)*Shift_Amplitude*(1-Attenuate_0_Adaptor*0.5));
        elseif Ad_Num == 4 | Ad_Num == size(Adaptors,2)-2
            Shift_Mag(1:Population_N) = (Tuning_Shift_Multiples(Ad_Index)*Shift_Amplitude*(1-Attenuate_0_Adaptor*0.25));
        else
            Shift_Mag(1:Population_N) = (Tuning_Shift_Multiples(Ad_Index)*Shift_Amplitude);
        end
        Shift_Mag(1:Population_N) = (Tuning_Shift_Multiples(Ad_Index)*Shift_Amplitude);
        

        
         for Ch_Num = 1 : Population_N
            % Generate adapted model channels - normal probability density
            % functions, centered on different physical test values
            % (Chn_Labels) with a standard deviation (sigma)
            Adapted_Channels(Ch_Num,:) = normpdf(x,Chn_Labels(Ch_Num)+Shift_Mag(Ch_Num),sigma);
            
            % Make sure that the baseline channels start scaled 0 to 1
            Adapted_Channels(Ch_Num,:) = Adapted_Channels(Ch_Num,:)./max(Adapted_Channels(Ch_Num,:));
            
             % Reduce channel gains, to simulate adaptation. Mitigate this
             % impact if the adaptor is matched to the average test value
            if Ad_Num == 1 
                % 0 Adaptation
                Adapted_Channels(Ch_Num,:) = Adapted_Channels(Ch_Num,:).*(1-Adapt_Mag(Ch_Num)*Ad_Amplitude*(1-Attenuate_0_Adaptor));
            elseif Ad_Num == 2 | Ad_Num == size(Adaptors,2)
                % 10 Adaptation
                Adapted_Channels(Ch_Num,:) = Adapted_Channels(Ch_Num,:).*(1-Adapt_Mag(Ch_Num)*Ad_Amplitude*(1-Attenuate_0_Adaptor*0.8));
            elseif Ad_Num == 3 | Ad_Num == size(Adaptors,2)-1
                % 20 Adaptation
                Adapted_Channels(Ch_Num,:) = Adapted_Channels(Ch_Num,:).*(1-Adapt_Mag(Ch_Num)*Ad_Amplitude*(1-Attenuate_0_Adaptor*0.6));
            elseif Ad_Num == 4 | Ad_Num == size(Adaptors,2)-2
                % 30 Adaptation
                Adapted_Channels(Ch_Num,:) = Adapted_Channels(Ch_Num,:).*(1-Adapt_Mag(Ch_Num)*Ad_Amplitude*(1-Attenuate_0_Adaptor*0.4));
            elseif Ad_Num == 5 | Ad_Num == size(Adaptors,2)-3
                % 40 Adaptation
                Adapted_Channels(Ch_Num,:) = Adapted_Channels(Ch_Num,:).*(1-Adapt_Mag(Ch_Num)*Ad_Amplitude*(1-Attenuate_0_Adaptor*0.2));
            else
                Adapted_Channels(Ch_Num,:) = Adapted_Channels(Ch_Num,:).*(1-Adapt_Mag(Ch_Num)*Ad_Amplitude);
            end
            
            % Now multiply the channel's gain potential by encoding noise 
            % (same noise levels as for baseline) 
            Adapted_Channels(Ch_Num,:) = Adapted_Channels(Ch_Num,:)*(Channel_Tuning_Noise(Ch_Num));
            
         end
         
        Adapted_Votes = [];     % Prepare an empty array to store channel votes in
        Vote_Labels = Chn_Labels+180;
        % Calculate adapted model encoding, by taking a weighted sum of 
        % channel 'votes' (e.g. adapted channel responses to the test). 
        % Channel labels are multiplied by channel response magnitudes 
        % (which are divided by the total response of ALL channels)
        Adapted_Votes(1:Population_N) = Vote_Labels(:).*((Adapted_Channels(:,Test_idx))./sum(Adapted_Channels(:,Test_idx)));
        
        % Sum the weighted votes of each channel, to determine the encoded value for this trial
        Adapted_Decoded_Vals(Trial,Ad_Num) = sum(Adapted_Votes)-180; 
        
        % Categorise input as Negative or Positive - by recording when trials
        % have been encoded as a positive value
        if Adapted_Decoded_Vals(Trial,Ad_Num) >= 0
            Adapted_Categorisation_Array(Results_Row,2,Ad_Num)=Adapted_Categorisation_Array(Results_Row,2,Ad_Num)+1;
            Category_Resp=1;
        else
            Category_Resp=0;
        end
        % Record the number of times a test value has been sampled
        Adapted_Categorisation_Array(Results_Row,3,Ad_Num)=Adapted_Categorisation_Array(Results_Row,3,Ad_Num)+1;

    
        % Categorise confidence as low or high, relative to the confidence
        % criterion (the unsigned encoded value needed to elicit a high-level of 
        % confidence) by recording when trials have exceeded the criterion 
        if abs(Adapted_Decoded_Vals(Trial,Ad_Num)) >= Confidence_Criterion
            Adapted_Confidence_Array(Results_Row,2,Ad_Num)=Adapted_Confidence_Array(Results_Row,2,Ad_Num)+1;
            Confidence_Resp=1;
        else
            Confidence_Resp=0;
        end
        % Record the number of times a test value has been sampled
        Adapted_Confidence_Array(Results_Row,3,Ad_Num)=Adapted_Confidence_Array(Results_Row,3,Ad_Num)+1; 
        
    end

end


%% Calculate and plot results of simulated experiment(s)

% Calculate the proportion of times each input resulted in a
% low-confidence categorisation at baseline (1-Certainty estimates)
Baseline_Confidence_Ys = 1-(Baseline_Confidence_Array(:,2)./Baseline_Confidence_Array(:,3));
% Transpose test values for plotting / curve fitting
Test_Vals=Test_Vals';

% Fit a Guassian distribution to proportions of low-confidence trials at
% baseline (uncertainty), as a function of baseline inputs
Baseline_Fit = fit(Test_Vals,Baseline_Confidence_Ys,'gauss1');

figure(1) % Prepare to plot results in Figure 1
clf % Clear the figure
subplot(2,1,1)

% Plot the results
p = plot(Baseline_Fit,Test_Vals,Baseline_Confidence_Ys);
set(p,{'LineWidth'},{3}); % Set the plot line width to 3
set(p,{'MarkerSize'},{20}); % Set the marker size
legend boxoff % Get rid of the legend
ax=gca; % Target the current access

% Target the fitted curve
TheFittedLine = (ax.Children(1).YData);
% Find the half max value of curve fit
halfMax = (min(TheFittedLine) + max(TheFittedLine)) / 2;
% Find where the data first drops below half the max.
index1 = find(TheFittedLine >= halfMax, 1, 'first');
% Find where the data last rises above half the max.
index2 = find(TheFittedLine >= halfMax, 1, 'last');
FWHH = index2-index1 + 1; % calculate FWHH in indices
% Convert FWHH (full width at half height) to angular degrees 
Baseline_Confidence_FWHH = (max(Test_Vals)-min(Test_Vals)) *(FWHH/size(TheFittedLine,2));

% Add some labels to the figure
Title_Text = strcat('Baseline Confidence');
title(Title_Text)
hold on % Then mark the limits of the FWHH on the plot
plot([Baseline_Fit.b1-(Baseline_Confidence_FWHH*0.5),Baseline_Fit.b1-(Baseline_Confidence_FWHH*0.5)],[halfMax,0],'k','LineWidth',3);
plot([Baseline_Fit.b1+(Baseline_Confidence_FWHH*0.5),Baseline_Fit.b1+(Baseline_Confidence_FWHH*0.5)],[halfMax,0],'k','LineWidth',3);

% Calculate Baseline confidence PSE - from peak uncertainty (in degrees)
Baseline_Confidence_PSE = Baseline_Fit.b1;


options = struct;   % initialize as an empty structure, to store curve fitting options
options.sigmoidName = 'norm';   % choose a cumulative Gaussian as the sigmoid
options.expType = "YesNo"       % Specify the design of the 'experiment' - a forced choice categorisation task 
options.widthalpha     = 0.25;   % Specify value for JND
options.estimateType   = 'mean';
options.stepN=[20,20,10,10,10];

% Fit a cumulative Gaussian psychmetric curve to categorisation data
Psychometric = psignifit(Baseline_Categorisation_Array,options);
% Record Baseline category PSE
Baseline_PSE = Psychometric.Fit(1)


%% Plot results of Psychometric function fit to Baseline data
figure(3)
clf
subplot(2,1,1)
box off
Title_Text = strcat('Baseline Categorisations:');
title(Title_Text)
plotOptions.dataColor = [0,0,0];  % color of the data
plotOptions.yLabel = 'Pr. Right Categorisations';
plotOptions.xLabel = 'Input Directions';
plotPsych(Psychometric,plotOptions);
ax=gca;
TheFittedLine = (ax.Children(6).YData);
PSE_idx = find(TheFittedLine >= 1/4, 1, 'first');
JND_idx = find(TheFittedLine >= 1-(1/4), 1, 'first');
% Record Baseline category JND (0.75% point of fitted function - 50% point)
Baseline_JND = ax.Children(6).XData(JND_idx) - ax.Children(6).XData(PSE_idx);
hold on % Later, we will plot the result of adapting to 0 below this

Baseline_Lower_JND = ax.Children(6).XData(PSE_idx)
Baseline_Upper_JND = ax.Children(6).XData(JND_idx)

%% Loop for calculating and plotting some adapted function fits
Model_PSE_AEs= []; % Array to store categorisation PSE aftereffects in
Local_Adapted_JND_AEs= []; % Array to store categorisation JND aftereffects in
Local_Adapted_Confidence_PSEs = []; % Array to store confidence PSE aftereffects in
Local_Adapted_Confidence_FWHHs = [];% Array to store confidence FWHH aftereffects in

for Ad_Num = 1 : size(Adaptors,2)

    % Fit a psychometric function to categorisation data for each adaptor
    Local_Ad_Psychometric = psignifit(Adapted_Categorisation_Array(:,:,Ad_Num),options);
    % Record PSE
    Local_Adapted_PSE(Ad_Num) = Local_Ad_Psychometric.Fit(1);

    figure(8)
    clf
    plotPsych(Local_Ad_Psychometric,plotOptions);
    ax=gca;
    TheFittedLine = (ax.Children(6).YData);
    PSE_idx = find(TheFittedLine >= 1/4, 1, 'first');
    JND_idx = find(TheFittedLine >= 1-(1/4), 1, 'first');
    % Record JND
    Local_Adapted_JND(Ad_Num) = ax.Children(6).XData(JND_idx) - ax.Children(6).XData(PSE_idx);


    % If you have adapted to 40, plot the results of that BELOW baseline
    % plot
    if Ad_Num == 5
        figure(3)
        hold on
        subplot(2,1,2)
        box off
        plotOptions.dataColor = [0,0,0];  % color of the data
        plotOptions.yLabel = 'Pr. Right Categorisations';
        plotOptions.xLabel = 'Input Directions';
        plotPsych(Local_Ad_Psychometric,plotOptions);
        Title_Text = strcat('Adapted Categorisations: Adaptor...',num2str(Adaptors(Ad_Num)),' degs');
        title(Title_Text)
        
        ax=gca;
        TheFittedLine = (ax.Children(6).YData);
        PSE_idx = find(TheFittedLine >= 1/4, 1, 'first');
        JND_idx = find(TheFittedLine >= 1-(1/4), 1, 'first');
        Ad_Lower_JND = ax.Children(6).XData(PSE_idx)
        Ad_Upper_JND = ax.Children(6).XData(JND_idx)
    end

    % Calculate categorisation PSE aftereffect (Adapted - Baseline PSE)
    Model_PSE_AEs(Ad_Num) = Local_Adapted_PSE(Ad_Num)-Baseline_PSE;

    % Calculate categorisation JND aftereffect (Adapted - Baseline JND)
    Model_JND_AEs(Ad_Num) = Baseline_JND-Local_Adapted_JND(Ad_Num);


    %% Fit a Guassian distribution to proportions of low-confidence trials 
    % (uncertainty) for each adaptor, as a function of baseline inputs
    Local_Adapted_Confidence_Ys = 1-(Adapted_Confidence_Array(:,2,Ad_Num)./Adapted_Confidence_Array(:,3,Ad_Num));
    Local_Fit = fit(Test_Vals,Local_Adapted_Confidence_Ys,'gauss1');

    % If you have adapted to 0, plot the results of that BELOW baseline
    % plot
    if Ad_Num == 1
        figure(1)
        hold on
        subplot(2,1,2)
    else
        figure(2)
        clf
    end
    p = plot(Local_Fit,Test_Vals,Local_Adapted_Confidence_Ys);
    set(p,{'LineWidth'},{3});
    set(p,{'MarkerSize'},{20});
    legend boxoff
    ax=gca;
    % Target the fitted curve
    TheFittedLine = (ax.Children(1).YData);
    % Find the half max value 
    halfMax = (min(TheFittedLine) + max(TheFittedLine)) / 2;
    % Find where the data first drops below half the max.
    index1 = find(TheFittedLine >= halfMax, 1, 'first');
    % Find where the data last rises above half the max.
    index2 = find(TheFittedLine >= halfMax, 1, 'last');
    FWHH = index2-index1 + 1; % FWHH in indices
    % Calculate and record FWHH in degrees
    Local_Adapted_FWHH_Degs(Ad_Num) = (max(Test_Vals)-min(Test_Vals)) *(FWHH/size(TheFittedLine,2));

    % Add some labels to plot
    Title_Text = strcat('Local adapted Confidence: Adaptor...',num2str(Adaptors(Ad_Num)),' degs');
    title(Title_Text)
    hold on
    % Plot limits of FWHH 
    plot([Local_Fit.b1-(Local_Adapted_FWHH_Degs(Ad_Num)*0.5),Local_Fit.b1-(Local_Adapted_FWHH_Degs(Ad_Num)*0.5)],[halfMax,0],'k','LineWidth',3);
    plot([Local_Fit.b1+(Local_Adapted_FWHH_Degs(Ad_Num)*0.5),Local_Fit.b1+(Local_Adapted_FWHH_Degs(Ad_Num)*0.5)],[halfMax,0],'k','LineWidth',3);

    % Record Uncertainty PSE (peak uncertainty - in degrees)
    Local_Adapted_Confidence_PSEs(Ad_Num) = Local_Fit.b1;    

    % Calculate Confidence PSE aftereffect (Adapted - Baseline)
    Model_Adapted_Confidence_PSE_AEs(Ad_Num) = Local_Adapted_Confidence_PSEs(Ad_Num)-Baseline_Confidence_PSE;
    % Calculate Confidence FWHH aftereffect (Adapted - Baseline)
    Model_Adapted_Confidence_FWHH_AEs(Ad_Num) = Baseline_Confidence_FWHH-Local_Adapted_FWHH_Degs(Ad_Num);    

end

temp1 = -flip(Model_Adapted_Confidence_PSE_AEs);
temp1(1) = Model_Adapted_Confidence_PSE_AEs(size(Model_Adapted_Confidence_PSE_AEs,2));
temp1(size(Model_Adapted_Confidence_PSE_AEs,2):-1+size(Model_Adapted_Confidence_PSE_AEs,2)*2)=Model_Adapted_Confidence_PSE_AEs;
Model_Adapted_Confidence_PSE_AEs = temp1;

temp2 = flip(Model_Adapted_Confidence_FWHH_AEs);
temp2(1) = Model_Adapted_Confidence_FWHH_AEs(size(Model_Adapted_Confidence_FWHH_AEs,2));
temp2(size(Model_Adapted_Confidence_FWHH_AEs,2):-1+size(Model_Adapted_Confidence_FWHH_AEs,2)*2)=Model_Adapted_Confidence_FWHH_AEs;
Model_Adapted_Confidence_FWHH_AEs = temp2;

temp3 = -flip(Model_PSE_AEs);
temp3(1) = Model_PSE_AEs(size(Model_PSE_AEs,2));
temp3(size(Model_PSE_AEs,2):-1+size(Model_PSE_AEs,2)*2)=Model_PSE_AEs;
Model_PSE_AEs = temp3;

temp4 = flip(Model_JND_AEs);
temp4(1) = Model_JND_AEs(size(Model_JND_AEs,2));
temp4(size(Model_JND_AEs,2):-1+size(Model_JND_AEs,2)*2)=Model_JND_AEs;
Model_JND_AEs = temp4;

Adaptors = -180:Channel_Spacing:180;


%% Tilt AEs, JND AEs, Confidence PSEs and Confidence FWHHs from our data...
Confidence_PSE_AEs = [ 4.05 	 14.62 	 16.84 	 13.35 	 6.58 	 2.35 	 1.43 	-0.48 	 1.77 	 1.81 
 0.34 	 6.27 	 8.96 	 12.00 	 3.19 	-0.78 	 0.47 	 0.15 	-1.03 	-0.30 
-2.00 	 12.69 	 14.31 	 7.68 	 6.28 	-2.19 	-2.55 	-4.28 	-1.63 	 0.72 
-0.15 	 7.32 	 10.01 	 9.43 	 5.13 	 3.33 	 1.21 	 1.40 	-0.38 	 2.08 
 0.18 	 10.92 	 11.60 	 10.23 	 8.03 	 0.82 	-0.80 	 0.89 	-1.77 	-2.16 
 0.44 	 7.02 	 10.25 	 6.24 	 4.55 	 1.35 	-0.39 	-1.62 	-1.02 	-2.23 
-0.27 	 11.26 	 11.93 	 11.39 	 2.53 	 3.47 	 1.83 	-3.18 	-3.02 	-1.18 
 4.49 	 10.96 	 14.01 	 11.56 	 11.58 	 7.46 	 4.31 	 3.25 	 4.60 	 5.84 
-0.28 	 7.63 	 10.84 	 11.66 	 8.55 	 3.29 	 1.54 	 0.00 	-0.88 	-0.88 
 0.54 	 8.54 	 14.27 	 9.67 	 6.18 	 3.12 	-1.36 	 0.21 	-1.58 	-1.49 
 7.10 	 12.94 	 18.03 	 16.09 	 7.99 	 7.82 	 7.94 	 5.77 	 8.00 	 9.19 
 1.76 	 7.50 	 7.41 	 5.56 	 2.77 	 1.61 	-0.57 	-1.70 	-1.64 	-1.20 
 NaN 	 NaN 	 3.03 	 9.81 	 0.28 	 9.74 	 NaN 	 NaN 	 NaN 	-3.23 
 0.37 	 6.50 	 10.40 	 4.24 	 1.29 	-0.83 	-7.41 	-7.65 	-7.46 	-5.52 
 0.12 	 5.39 	 5.53 	 4.16 	 2.31 	-0.15 	 1.30 	 0.15 	-0.30 	 1.18 
 0.36 	 4.29 	 8.69 	 6.86 	 3.60 	-0.69 	-0.54 	-0.18 	 1.70 	 0.14 
 0.68 	 9.43 	 16.36 	 11.87 	 5.66 	 4.04 	 0.71 	-0.11 	-1.46 	 1.04 
 4.64 	 8.36 	 10.15 	 11.04 	 8.90 	 4.15 	 1.30 	 6.08 	 3.36 	 1.83 
-5.75 	 0.07 	 NaN 	-4.20 	 NaN 	 5.40 	-17.69 	-9.54 	-9.61 	-13.18 
];

Our_Confidence_PSE_AEs = nanmean(Confidence_PSE_AEs,1);
Our_Confidence_PSE_AEs = -flip(Our_Confidence_PSE_AEs);
Our_Confidence_PSE_AEs(size(Our_Confidence_PSE_AEs,2):size(Our_Confidence_PSE_AEs,2)+size(Our_Confidence_PSE_AEs,2)-1)=-flip(Our_Confidence_PSE_AEs)

Our_Confidence_PSE_SDs = nanstd(Confidence_PSE_AEs,1,1);
Our_Confidence_PSE_SDs = Our_Confidence_PSE_SDs./sqrt(size(Our_Confidence_PSE_AEs,1));
Our_Confidence_PSE_SDs = flip(Our_Confidence_PSE_SDs);
Our_Confidence_PSE_SDs(size(Our_Confidence_PSE_SDs,2):size(Our_Confidence_PSE_SDs,2)+size(Our_Confidence_PSE_SDs,2)-1)= flip(Our_Confidence_PSE_SDs)

Perception_PSE_AEs = [ 3.21 	 15.68 	 20.24 	 14.80 	 5.82 	 3.64 	 0.98 	-0.65 	 0.33 	 2.14 
 0.27 	 9.61 	 8.29 	 9.46 	 3.03 	-1.15 	-0.90 	-0.50 	-2.21 	-0.17 
 0.28 	 10.84 	 15.28 	 11.57 	 6.41 	 0.20 	-5.27 	-3.72 	-2.08 	 2.13 
 2.73 	 9.27 	 13.78 	 13.36 	 6.02 	 5.31 	 4.31 	 2.93 	 2.35 	 2.47 
 1.85 	 11.80 	 13.40 	 13.70 	 8.68 	 0.66 	 0.84 	-0.59 	-2.29 	 0.11 
 0.07 	 6.20 	 10.22 	 5.61 	 3.22 	-0.39 	-1.26 	-3.33 	-1.50 	-2.88 
-0.49 	 11.98 	 12.42 	 11.39 	 3.61 	 3.87 	 0.62 	-1.75 	-2.29 	-0.00 
 4.59 	 10.30 	 17.88 	 14.69 	 14.02 	 9.11 	 3.63 	 3.40 	 2.73 	 7.08 
 1.50 	 10.69 	 10.43 	 12.23 	 9.66 	 5.10 	 3.39 	-0.50 	-0.13 	-0.80 
-0.88 	 12.65 	 12.94 	 8.47 	 5.27 	 1.06 	-0.99 	-1.70 	-4.29 	-3.05 
 5.13 	 6.07 	 16.05 	 16.27 	 2.21 	 9.62 	 7.02 	 6.11 	 2.22 	 6.79 
-1.68 	 7.43 	 4.96 	 7.48 	 4.18 	 0.28 	-0.63 	-4.18 	-2.08 	-4.76 
 9.13 	 NaN 	 8.04 	 8.46 	 1.47 	 4.42 	-21.30 	 NaN 	 NaN 	-2.28 
-0.22 	 4.75 	 9.60 	 6.30 	 1.04 	 0.22 	-7.93 	-6.10 	-5.71 	-6.53 
 1.22 	 8.37 	 4.47 	 4.43 	 2.19 	 0.23 	 1.26 	 1.26 	 1.69 	 0.91 
-0.54 	 2.34 	 7.36 	 8.50 	 0.93 	-2.41 	-2.88 	 0.15 	 5.07 	 1.27 
 3.45 	 12.66 	 12.85 	 10.53 	 6.08 	 2.36 	 2.97 	-1.62 	 0.34 	 2.04 
 2.64 	 6.17 	 9.89 	 8.21 	 11.24 	 2.72 	 2.23 	 6.45 	 5.31 	 1.69 
-3.57 	-3.68 	 NaN 	 NaN 	 NaN 	-2.18 	-13.26 	-7.00 	-9.40 	-9.30 
];

Our_Tilt_AEs = nanmean(Perception_PSE_AEs,1);
Our_Tilt_AEs = -flip(Our_Tilt_AEs);
Our_Tilt_AEs(size(Our_Tilt_AEs,2):size(Our_Tilt_AEs,2)+size(Our_Tilt_AEs,2)-1)=-flip(Our_Tilt_AEs)

Our_Tilt_SDs = nanstd(Perception_PSE_AEs,1,1);
Our_Tilt_SDs = Our_Tilt_SDs./sqrt(size(Our_Tilt_AEs,1));
Our_Tilt_SDs = flip(Our_Tilt_SDs);
Our_Tilt_SDs(size(Our_Tilt_SDs,2):size(Our_Tilt_SDs,2)+size(Our_Tilt_SDs,2)-1)=flip(Our_Tilt_SDs)

Confidence_FWHH_AEs = [ 8.11 	-3.08 	-7.02 	-10.29 	-0.13 	 0.79 	 3.27 	-5.68 	-3.87 	-1.36 
-1.49 	 8.03 	-6.43 	-10.13 	-2.00 	-3.39 	-1.05 	-2.79 	-2.51 	-1.84 
 6.78 	 3.59 	-11.42 	-1.60 	-2.89 	 0.45 	-3.87 	 5.26 	 4.63 	 1.49 
 2.54 	-5.61 	-7.51 	-0.81 	-0.48 	-2.38 	 0.48 	-0.86 	 2.41 	 0.19 
 3.11 	 0.69 	-8.55 	-0.27 	 0.62 	 0.25 	-2.47 	-3.90 	 1.30 	-0.18 
 0.57 	-2.89 	-6.72 	-6.86 	-3.11 	-0.66 	-2.66 	-4.63 	-3.33 	 1.07 
 3.51 	-1.59 	-17.83 	-11.96 	-9.92 	 0.41 	 0.81 	-6.46 	-4.69 	-0.92 
 1.59 	 3.82 	-8.80 	-0.33 	-2.90 	 1.59 	-4.36 	-0.63 	-0.80 	-1.90 
 4.50 	 0.34 	-6.28 	 4.16 	-9.01 	-0.89 	-3.65 	-3.17 	-0.03 	-2.38 
 1.59 	-0.66 	-13.93 	-8.58 	-7.39 	-4.85 	-5.17 	 0.29 	-7.54 	-0.26 
 8.98 	 4.14 	-9.52 	-4.50 	 4.28 	 2.73 	 1.45 	 0.60 	 6.55 	 6.03 
 4.39 	 5.27 	-4.77 	-2.62 	-2.41 	-2.09 	-5.20 	-3.39 	-3.43 	-3.17 
 7.17 	 NaN 	-15.50 	 0.55 	-13.82 	-0.60 	 NaN 	 NaN 	 NaN 	-2.42 
 0.79 	 1.49 	-4.16 	-2.25 	-2.54 	-2.79 	-0.10 	-7.97 	-8.40 	 1.08 
 4.15 	 2.16 	-2.80 	-2.57 	-4.12 	-5.76 	-15.83 	-2.73 	-5.33 	-4.28 
 2.73 	 2.49 	-4.67 	-7.46 	-4.36 	 1.84 	-0.57 	 0.44 	-0.69 	 0.77 
 0.51 	 2.59 	-9.85 	-3.29 	-3.20 	 0.13 	-3.26 	-3.95 	 0.10 	 1.71 
 18.18 	 1.29 	-12.04 	-12.11 	-5.69 	 3.97 	 1.26 	 2.36 	-1.57 	 0.55 
-3.74 	-7.38 	-5.22 	-5.26 	 NaN 	-7.55 	-9.54 	 2.02 	-15.18 	 NaN 
];

Our_Confidence_FWHH_AEs = nanmean(Confidence_FWHH_AEs,1);
Our_Confidence_FWHH_AEs = flip(Our_Confidence_FWHH_AEs);
Our_Confidence_FWHH_AEs(size(Our_Confidence_FWHH_AEs,2):size(Our_Confidence_FWHH_AEs,2)+size(Our_Confidence_FWHH_AEs,2)-1)=flip(Our_Confidence_FWHH_AEs);

Our_Confidence_FWHH_SDs = nanstd(Confidence_FWHH_AEs,1,1);
Our_Confidence_FWHH_SDs = Our_Confidence_FWHH_SDs./sqrt(size(Our_Confidence_FWHH_AEs,1));
Our_Confidence_FWHH_SDs = flip(Our_Confidence_FWHH_SDs);
Our_Confidence_FWHH_SDs(size(Our_Confidence_FWHH_SDs,2):size(Our_Confidence_FWHH_SDs,2)+size(Our_Confidence_FWHH_SDs,2)-1)=flip(Our_Confidence_FWHH_SDs);

Perception_JND_AEs = [-2.14 	-0.77 	-1.74 	-1.14 	 0.91 	-1.51 	 0.97 	-0.26 	-0.60 	-1.39 
 0.91 	 1.37 	-1.50 	-0.32 	-2.13 	 0.46 	 0.21 	 0.86 	 0.72 	 0.27 
 0.07 	 0.14 	-2.89 	-3.46 	-0.95 	 0.00  	-0.84 	-1.65 	 0.63 	 0.61 
 0.09 	-2.94 	-3.45 	-2.62 	-3.58 	-1.31 	-1.52 	-3.53 	-1.89 	-1.09 
 0.42 	 0.00  	-1.29 	-1.63 	-0.04 	-1.59 	-1.07 	-0.86 	-0.29 	 0.56 
-0.16 	 0.51 	-4.27 	-3.81 	-2.07 	-0.86 	-0.36 	 0.77 	-0.69 	-1.08 
 0.69 	-0.64 	-4.36 	-0.11 	-0.61 	-2.79 	-0.17 	-0.00 	-1.57 	-1.98 
-1.50 	-0.23 	-2.26 	-3.25 	-0.47 	-0.16 	 0.01 	-0.59 	 1.99 	-2.92 
-0.56 	-2.35 	-9.92 	 0.29 	 0.85 	 0.17 	-3.47 	-1.66 	 0.12 	-0.43 
-2.32 	 0.13 	-6.59 	-5.38 	-1.68 	-1.03 	-2.42 	 0.68 	 0.18 	 1.51 
-0.07 	-2.39 	-1.19 	-4.17 	-0.80 	-0.42 	-1.79 	-0.80 	-2.59 	-1.26 
-0.34 	 0.59 	-4.60 	-3.30 	-1.22 	-0.17 	-1.79 	-2.78 	-1.59 	-0.81 
 1.73 	 NaN 	 3.69 	-0.62 	 0.49 	-4.55 	 NaN 	 NaN 	 NaN 	 0.73 
 0.15 	-2.07 	-1.40 	-0.22 	 0.04 	-0.74 	-0.50 	-2.83 	-1.61 	 0.93 
-0.64 	 2.60 	-1.04 	-0.81 	-2.18 	 0.12 	-1.04 	-1.39 	 0.08 	-1.09 
 0.17 	-0.04 	-1.07 	-0.84 	 0.68 	 1.50 	 0.74 	 2.36 	-0.26 	 2.79 
 0.76 	-1.89 	-5.69 	-2.15 	-0.78 	-0.16 	-0.26 	-3.91 	-1.83 	-0.38 
-1.13 	-1.78 	-0.35 	 0.09 	 1.01 	-2.00 	-0.35 	 0.51 	-1.96 	 0.71 
 1.62 	 2.40 	-2.60 	-2.91 	-5.11 	-0.08 	 1.39 	-0.18 	 0.12 	 0.57 
];

Our_Tilt_JND_AEs = nanmean(Perception_JND_AEs,1);
Our_Tilt_JND_AEs = flip(Our_Tilt_JND_AEs);
Our_Tilt_JND_AEs(size(Our_Tilt_JND_AEs,2):size(Our_Tilt_JND_AEs,2)+size(Our_Tilt_JND_AEs,2)-1)=flip(Our_Tilt_JND_AEs);

Our_Tilt_JND_SDs = nanstd(Perception_JND_AEs,1,1);
Our_Tilt_JND_SDs = Our_Tilt_JND_SDs./sqrt(size(Perception_JND_AEs,1));
Our_Tilt_JND_SDs = flip(Our_Tilt_JND_SDs);
Our_Tilt_JND_SDs(size(Our_Tilt_JND_SDs,2):size(Our_Tilt_JND_SDs,2)+size(Our_Tilt_JND_SDs,2)-1)=flip(Our_Tilt_JND_SDs);

Our_Adaptors = [-180 -160 -140 -120 -100 -80 -60 -40 -20 0	20	40	60	80	100	120	140 160	180];


%% Plot Local Adapted Aftereffect functions...

figure(6)
clf
title('Local Adapted Functions') 
subplot(2,2,1)
% Plot categorical PSE aftereffect function...
plot(Adaptors,Model_PSE_AEs,'k','LineWidth',3);
hold on
%plot(GandR_Adaptors,GandR_Tilt_AEs,'b*');
plot(Our_Adaptors,Our_Tilt_AEs,'r*');
plot([-180,180],[0,0],'k','LineWidth',2);

% Plot categorical JND aftereffect function...
subplot(2,2,2)
plot(Adaptors,Model_JND_AEs,'k','LineWidth',3);
hold on
%plot(RandB_Adaptors,RandB_TD_Changes,'b*');
plot(Our_Adaptors,Our_Tilt_JND_AEs,'r*');
plot([-180,180],[0,0],'k','LineWidth',2);

% Plot Uncertainty PSE aftereffect function...
subplot(2,2,3)
plot(Adaptors,Model_Adapted_Confidence_PSE_AEs,'k','LineWidth',3);
hold on
plot(Our_Adaptors,Our_Confidence_PSE_AEs,'r*');
plot([-180,180],[0,0],'k','LineWidth',2);

% Plot Uncertainty FWHH aftereffect function...
subplot(2,2,4)
plot(Adaptors,Model_Adapted_Confidence_FWHH_AEs,'k','LineWidth',3);
hold on
plot(Our_Adaptors,Our_Confidence_FWHH_AEs,'r*');
plot([-180,180],[0,0],'k','LineWidth',2);
        


%% Plot Local Adapted Aftereffect functions...

figure(11)
clf
subplot(1,2,1)
% Plot smoothed categorical PSE aftereffect function...
samplingRateIncrease = 10;
lengthX = length(Adaptors);
newXSamplePoints = linspace(min(Adaptors), max(Adaptors), lengthX * samplingRateIncrease);
smoothed_PSE_AEs = spline(Adaptors, Model_PSE_AEs, newXSamplePoints);
smoothed_Adaptors = newXSamplePoints;
plot1 = plot(smoothed_Adaptors,smoothed_PSE_AEs,'b','LineWidth',3);
plot1.Color(4) = 0.2;
hold on
plot(Our_Adaptors,Our_Tilt_AEs,'b.','MarkerSize',24);
for ErrBars = 1 : size(Our_Adaptors,2)
    plot([Our_Adaptors(ErrBars),Our_Adaptors(ErrBars)],[Our_Tilt_AEs(ErrBars)-Our_Tilt_SDs(ErrBars),Our_Tilt_AEs(ErrBars)+Our_Tilt_SDs(ErrBars)],'color',[0 0 1 0.2],'LineWidth',2)
end
plot([-180,180],[0,0],'k','LineWidth',2);
hold on

% Plot categorical JND aftereffect function...
smoothed_Model_Adapted_Confidence_PSE_AEs = spline(Adaptors, Model_Adapted_Confidence_PSE_AEs, newXSamplePoints);
%subplot(1,3,2)
plot2 = plot(smoothed_Adaptors,smoothed_Model_Adapted_Confidence_PSE_AEs,'r','LineWidth',3);
plot2.Color(4) = 0.2;
hold on
plot(Our_Adaptors,Our_Confidence_PSE_AEs,'r.','MarkerSize',24);
for ErrBars = 1 : size(Our_Adaptors,2)
    plot([Our_Adaptors(ErrBars),Our_Adaptors(ErrBars)],[Our_Confidence_PSE_AEs(ErrBars)-Our_Confidence_PSE_SDs(ErrBars),Our_Confidence_PSE_AEs(ErrBars)+Our_Confidence_PSE_SDs(ErrBars)],'color',[1 0 0 0.2],'LineWidth',2)
end
plot([-180,180],[0,0],'k','LineWidth',2);
xlabel('Adapted Orientation Difference (degs)','FontSize',16,'FontWeight','bold') 
ylabel({'Perceptual / Confidence';'PSE Aftereffects (degs)'},'FontSize',16,'FontWeight','bold')  
set(gca,'fontsize',14)

box off


% Plot Uncertainty PSE aftereffect function...
smoothed_Model_JND_AEs = spline(Adaptors, Model_JND_AEs, newXSamplePoints);
subplot(1,2,2)
plot3 = plot(smoothed_Adaptors,smoothed_Model_JND_AEs,'b','LineWidth',3);
plot3.Color(4) = 0.2;
hold on
plot(Our_Adaptors,Our_Tilt_JND_AEs,'b.','MarkerSize',24);
for ErrBars = 1 : size(Our_Adaptors,2)
    plot([Our_Adaptors(ErrBars),Our_Adaptors(ErrBars)],[Our_Tilt_JND_AEs(ErrBars)-Our_Tilt_JND_SDs(ErrBars),Our_Tilt_JND_AEs(ErrBars)+Our_Tilt_JND_SDs(ErrBars)],'color',[0 0 1 0.2],'LineWidth',2)
end
plot([-180,180],[0,0],'b','LineWidth',2);

% Plot Uncertainty FWHH aftereffect function...
smoothed_Model_Adapted_Confidence_FWHH_AEs = spline(Adaptors, Model_Adapted_Confidence_FWHH_AEs, newXSamplePoints);

plot4 = plot(smoothed_Adaptors,smoothed_Model_Adapted_Confidence_FWHH_AEs,'r','LineWidth',3);
plot4.Color(4) = 0.2;
hold on
plot(Our_Adaptors,Our_Confidence_FWHH_AEs,'r.','MarkerSize',24);
for ErrBars = 1 : size(Our_Adaptors,2)
    plot([Our_Adaptors(ErrBars),Our_Adaptors(ErrBars)],[Our_Confidence_FWHH_AEs(ErrBars)-Our_Confidence_FWHH_SDs(ErrBars),Our_Confidence_FWHH_AEs(ErrBars)+Our_Confidence_FWHH_SDs(ErrBars)],'color',[1 0 0 0.2],'LineWidth',2)
end
plot([-180,180],[0,0],'k','LineWidth',2);

xlabel('Adapted Orientation Difference (degs)','FontSize',16,'FontWeight','bold') 
ylabel({'Perceptual JND / Confidence FWHH';'Aftereffects (degs)'},'FontSize',16,'FontWeight','bold')  
set(gca,'fontsize',14)

box off
        

%%

figure(4)
subplot(1,3,1)
xlim([-40 40])
ylim([-40 40])
x1 = -40; x2 = 40;
y1 = 5/2; y2 = -5/2;
fill([x1 x1 x2 x2],[y1 y2 y2 y1],'m','LineStyle','none');
hold on
y1 = 40; y2 = 5/2;
fill([x1 x1 x2 x2],[y1 y2 y2 y1],[0.5 0.5 0.5],'LineStyle','none');
y1 = -5/2; y2 = -40;
fill([x1 x1 x2 x2],[y1 y2 y2 y1],[0.5 0.5 0.5],'LineStyle','none');
scatter(Tests(:),Decoded_Vals(:),'k','filled')
plot([-40,40],[0,0],'--g','LineWidth',3)
plot([-40,40],[-40,40],'r','LineWidth',3)
xlabel('Physical Input Orientations','FontSize',16,'FontWeight','bold')  
ylabel('Baseline Decoded Orientations','FontSize',16,'FontWeight','bold')  

subplot(1,3,2)
xlim([-40 40])
ylim([-40 40])
x1 = -40; x2 = 40;
y1 = 5/2; y2 = -5/2;
fill([x1 x1 x2 x2],[y1 y2 y2 y1],'m','LineStyle','none');
hold on
y1 = 40; y2 = 5/2;
fill([x1 x1 x2 x2],[y1 y2 y2 y1],[0.5 0.5 0.5],'LineStyle','none');
y1 = -5/2; y2 = -40;
fill([x1 x1 x2 x2],[y1 y2 y2 y1],[0.5 0.5 0.5],'LineStyle','none');
scatter(Tests(:),Adapted_Decoded_Vals(:,1),'k','filled')
xlim([-40 40])
ylim([-40 40])
plot([-40,40],[0,0],'--g','LineWidth',3)
plot([-40,40],[-40,40],'r','LineWidth',3)
xlabel('Physical Input Orientations','FontSize',16,'FontWeight','bold') 
ylabel('0-deg adapted Decoded Orientations','FontSize',16,'FontWeight','bold')  


subplot(1,3,3)
xlim([-40 40])
ylim([-40 40])
x1 = -40; x2 = 40;
y1 = 5/2; y2 = -5/2;
fill([x1 x1 x2 x2],[y1 y2 y2 y1],'m','LineStyle','none');
hold on
y1 = 40; y2 = 5/2;
fill([x1 x1 x2 x2],[y1 y2 y2 y1],[0.5 0.5 0.5],'LineStyle','none');
y1 = -5/2; y2 = -40;
fill([x1 x1 x2 x2],[y1 y2 y2 y1],[0.5 0.5 0.5],'LineStyle','none');
scatter(Decoded_Vals(:),Adapted_Decoded_Vals(:,1),'k','filled')
xlim([-40 40])
ylim([-40 40])
plot([-40,40],[0,0],'--g','LineWidth',3)
plot([-40,40],[-40,40],'r','LineWidth',3)
xlabel('Baseline Decoded Orientations','FontSize',16,'FontWeight','bold')  
ylabel('0-deg adapted Decoded Orientations','FontSize',16,'FontWeight','bold')  


%% Plots to highlight JND vs Confidence Differences
[vals idndices] = find(abs(Decoded_Vals)<=1);

figure(6)
clf
subplot(1,2,1)
scatter(Decoded_Vals(idndices),Adapted_Decoded_Vals(idndices,1),'k','filled')
h1=lsline;
h1.Color = 'r';
h1.LineWidth = 4;
yl = ylim
xlim([yl(1) yl(2)])

Correlate_Base_Data = Decoded_Vals;
[vals idndices] = find(Correlate_Base_Data<=4);
Correlate_Base_Data(idndices)=NaN;
[vals idndices] = find(Correlate_Base_Data>=6);
Correlate_Base_Data(idndices)=NaN;
[vals idndices] = find(Correlate_Base_Data>=4);

subplot(1,2,2)
scatter(abs(Decoded_Vals(idndices)),abs(Adapted_Decoded_Vals(idndices,1)),'k','filled')
h2=lsline;
h2.Color = 'r';
h2.LineWidth = 4;
yl = ylim
xlim([yl(1) yl(2)])


%%

figure(7)
clf
subplot(1,3,1)
xlim([-40 40])
ylim([-40 40])
x1 = -40; x2 = 40;
y1 = 20; y2 = -20;
fill([x1 x1 x2 x2],[y1 y2 y2 y1],'m','LineStyle','none');
hold on
y1 = 40; y2 = 7;
fill([x1 x1 x2 x2],[y1 y2 y2 y1],[0.5 0.5 0.5],'LineStyle','none');
y1 = -7; y2 = -40;
fill([x1 x1 x2 x2],[y1 y2 y2 y1],[0.5 0.5 0.5],'LineStyle','none');
scatter(Tests(:),Decoded_Vals(:),'k','filled')
xlim([-40 40])
ylim([-40 40])

plot([-40,40],[0,0],'--g','LineWidth',3)
plot([-40,40],[-40,40],'r','LineWidth',3)
xlabel('Physical Input Orientations','FontSize',16,'FontWeight','bold')  
ylabel('Baseline Decoded Orientations','FontSize',16,'FontWeight','bold')  

subplot(1,3,2)
xlim([-40 40])
ylim([-40 40])
x1 = -40; x2 = 40;
y1 = 7; y2 = -7;
fill([x1 x1 x2 x2],[y1 y2 y2 y1],'m','LineStyle','none');
hold on
y1 = 40; y2 = 7;
fill([x1 x1 x2 x2],[y1 y2 y2 y1],[0.5 0.5 0.5],'LineStyle','none');
y1 = -7; y2 = -40;
fill([x1 x1 x2 x2],[y1 y2 y2 y1],[0.5 0.5 0.5],'LineStyle','none');
scatter(Tests(:),Adapted_Decoded_Vals(:,5),'k','filled')
xlim([-40 40])
ylim([-40 40])

xlim([-40 40])
ylim([-40 40])
plot([-40,40],[0,0],'--g','LineWidth',3)
plot([-40,40],[-40,40],'r','LineWidth',3)
xlabel('Physical Input Orientations','FontSize',16,'FontWeight','bold') 
ylabel('40-deg adapted Decoded Orientations','FontSize',16,'FontWeight','bold')  


subplot(1,3,3)
xlim([-40 40])
ylim([-40 40])
x1 = -40; x2 = 40;
y1 = 7; y2 = -7;
fill([x1 x1 x2 x2],[y1 y2 y2 y1],'m','LineStyle','none');
hold on
y1 = 40; y2 = 7;
fill([x1 x1 x2 x2],[y1 y2 y2 y1],[0.5 0.5 0.5],'LineStyle','none');
y1 = -7; y2 = -40;
fill([x1 x1 x2 x2],[y1 y2 y2 y1],[0.5 0.5 0.5],'LineStyle','none');
scatter(Decoded_Vals(:),Adapted_Decoded_Vals(:,5),'k','filled')
xlim([-40 40])
ylim([-40 40])

xlim([-40 40])
ylim([-40 40])
plot([-40,40],[0,0],'--g','LineWidth',3)
plot([-40,40],[-40,40],'r','LineWidth',3)
xlabel('Baseline Decoded Orientations','FontSize',16,'FontWeight','bold')  
ylabel('40-deg adapted Decoded Orientations','FontSize',16,'FontWeight','bold')  


%% Find out what 40 degree adapted tests have been encoded as +/- 1 JND from 0
Popn = Adapted_Decoded_Vals(:,5);
% Lower_JND = Local_Adapted_PSE(5)-Local_Adapted_JND(5)/2;
% Upper_JND = Local_Adapted_PSE(5)+Local_Adapted_JND(5)/2;
Lower_JND = -Local_Adapted_JND(5)*0.75;
Upper_JND = +Local_Adapted_JND(5)*0.75;

[Lower_idxs val] = find(Popn > Lower_JND-0.5 & Popn < Lower_JND+0.5);
[Upper_idxs val] = find(Popn > Upper_JND-0.5 & Popn < Upper_JND+0.5);

Avg_40_Ad_Lower_JND = round(mean(Tests(Lower_idxs)),2)
Avg_40_Ad_Upper_JND = round(mean(Tests(Upper_idxs)),2)


%Popn = Decoded_Vals;
% Lower_JND = Baseline_PSE-Baseline_JND/2;
% Upper_JND = Baseline_PSE+Baseline_JND/2;
Lower_JND = -Baseline_JND*0.75;
Upper_JND = +Baseline_JND*0.75;

[val Lower_idxs] = find(Decoded_Vals > Lower_JND-0.25 & Decoded_Vals < Lower_JND+0.25);
[val Upper_idxs] = find(Decoded_Vals > Upper_JND-0.25 & Decoded_Vals < Upper_JND+0.25);

Avg_Baseline_Lower_JND = mean(Tests(Lower_idxs))
Avg_Baseline_Upper_JND = mean(Tests(Upper_idxs))


%% Plots to highlight JND vs Confidence Differences
[~, idndices] = find(abs(Decoded_Vals)<=1);

figure(8)
clf
subplot(1,2,1)
scatter(Decoded_Vals(idndices),Adapted_Decoded_Vals(idndices,5),'k','filled')
h3=lsline;
h3.Color = 'r';
h3.LineWidth = 4;
yl = ylim
xlim([yl(1) yl(2)])


Correlate_Base_Data = Decoded_Vals;
[vals idndices] = find(Correlate_Base_Data<=4);
Correlate_Base_Data(idndices)=NaN;
[vals idndices] = find(Correlate_Base_Data>=6);
Correlate_Base_Data(idndices)=NaN;
[vals idndices] = find(Correlate_Base_Data>=4);

subplot(1,2,2)
scatter(abs(Decoded_Vals(idndices)),abs(Adapted_Decoded_Vals(idndices,5)),'k','filled')
h4=lsline;
h4.Color = 'r';
h4.LineWidth = 4;
yl = ylim
xlim([yl(1) yl(2)])
```
