## Supplemental Matlab files for "On why we lack confidence in signal-detection-based analyses of confidence": Supp_2_Confidence_Population_Code_Model_SDT.m

```
%% This code runs simulated SDT-based experiments, to determine what shape 
% distributions of values are encoded by our labelled line observer model - 
% which is designed to account for MAE datasets. The code will either 
% simulate experiments where 101 different decisional criteria (bias   
% levels) are adopted for probability distributions (signal and noise)  
% that have a fixed offset (i.e. fixed sensitivity), or where 101 different
% different levels of sensitivity (distribution offsets) are sampled for an
% unbiased observer. 
%
% In each case, two model states are sampled - baseline (unadapted) and +40
% degree adapted.
%
% The model mimics human performance, by simulating levels of
% trial-by-trial encoding noise that match human behaviour, before and after
% adaptation. You can specify the number of simulated trials (500, 5000 or
% 50000). The code will output plots where gamma functions are fit to noise
% (red) and signal (green) distributions either generated from model output 
% (lower plots in Figures 1 & 2) or from random samples of normal 
% probability distributions with means and SDs set to match behavioural data 
% from our experiemnt (top plots in Figures 1 & 2). In these plots central
% black vertical  bars mark the average decision criterion value, and blue 
% bars mark confidence criteria.
%
% The code will also output XY scatter plots of d' and meta d' values 
% calculated based on encoded data from repeated (signal and noise) inputs
% via our model (red data), or from random samples of a normal distribution 
% (black data - Figure 3).
%
% If you simulate 50000 trials, the code will also plot d' : meta d'
% ratios and differences as a function of decision criteria distribution
% offsets (sensitivity), and perform an analysis to see if slopes of plots
% of z-scored Hit and Flase Alarm rates are different for (skewed) model 
% distributions of encoded values vs normally distributed distributions 
%
% The code needs psignifit to be installed, as this is used to fit
% functions to the results of simulated experiments...

clear all % Clear memory
close all % close all figures
% If psignifit has not already been added to your path, do that now
addpath /Users/uqdarnol/Dropbox/Research_Projects/Toolbox/Psignifit/psignifit-master

%% Specify the total number of trials to simulate
N_Repeats = 50000;      % Specify the number of times each test value will 
                        % be sampled. As the computer will not complain, 
                        % simulate A LOT of trials, to get an accuarate 
                        % estimate of modelled psychometric functions


%% Model parameters...
Baseline_JND = 4.5;
Confidence_Crit = 7;

Lower_Input = -2;
Upper_Input = 2;

Baseline_Lower_JNDs = Lower_Input-0.6:0.05:Lower_Input+0.6;
Baseline_Upper_JNDs = Upper_Input-0.6:0.05:Upper_Input+0.6;
Baseline_Upper_JNDs = flip(Baseline_Upper_JNDs);


Ad_PSE = 11.5;
Ad_JND = 6.7;

Ad_Lower_JNDs = (Ad_PSE+Lower_Input)-0.6:0.05:(Ad_PSE+Lower_Input)+0.6;
Ad_Upper_JNDs = (Ad_PSE+Upper_Input)-0.6:0.05:(Ad_PSE+Upper_Input)+0.6;
Ad_Upper_JNDs = flip(Ad_Upper_JNDs);

Lower_Input = Baseline_Lower_JNDs(1);
Upper_Input = Baseline_Upper_JNDs(1);

%% Model parameters

Channel_Spacing = 10;       % Spacing in-between the peak tunings of model channels

Gain_Factor = 1.8;          % Controls spread of reductions in channel gain potential post-adaptation
Shift_Factor = 4.5;         % Controls spread of channel tuning shifts post-adaptation

Ad_Amplitude =     0.95;    % Maximal amount of post-adaptation channel gain reduction 
Shift_Amplitude =   40;     % Maximal post-adaptation channel tuning shift

Confidence_Criterion = 7;   % Decoded angular difference beyond category 
                            % criterion (0) needed to evoke a 'high' 
                            % confidence response
                            
Attenuate_0_Adaptor = 0.5;  % Maximum attenuation of adaptation-driven gain reductions

sigma = 20;                 % Bandwidth of model channels

%% Pre-Generate adaptation induced channel gain reductions and tuning shifts
Xs = -180:Channel_Spacing:180; % An array of direction values, for use in calculating adaptation multiples
%Xs = -180:Channel_Spacing:180-Channel_Spacing; % An array of direction values, for use in calculating adaptation multiples

for n = 1 : size(Xs,2)
    
    if abs(Xs(n)*Gain_Factor)<=180
        % Multiple to reduce channel gain by, determined by angular
        % difference from the adaptor (0 degrees)
        Post_Ad_Gain_Reductions(n)=cosd(abs(Xs(n)*Gain_Factor)/2);
    else
        Post_Ad_Gain_Reductions(n)=0;
    end
    
    if sind(Xs(n)*Shift_Factor) < 0 & abs(Xs(n)*Shift_Factor)<=180
        % Multiple to shift channel tuning by, for negative angular
        % differences from the adaptor (0 degrees, so a repulsive tuning
        % shift)
        Tuning_Shift_Multiples(n)=-sind(abs(Xs(n)*Shift_Factor)/1);
        % Multiple to shift channel tuning by, for positive angular
        % differences from the adaptor (0 degrees, so a repulsive tuning
        % shift)
    elseif sind(Xs(n)*Shift_Factor) > 0 & abs(Xs(n)*Shift_Factor)<=180
        Tuning_Shift_Multiples(n)=sind(abs(Xs(n)*Shift_Factor)/1);
    else
        Tuning_Shift_Multiples(n) = 0;
    end
    
end

%% Parameters for calculating an unadapted population code

x = -180:0.05:180; % Create an array of x indicies, finely sampling the full gamut of possible directions
%x = round(x,2);
s = size(x,2);  % Record the size of the X axis indices array

Population_N = size(Tuning_Shift_Multiples,2); % Number of channels in population code
end1 = -180; % Specify the tuning of the first channel
end2 = 180-Channel_Spacing;  % Specify the tuning of the last channel
Chn_Labels=end1:((end2-end1)/(Population_N-1)):end2; % Determine ALL channel labels / tunings 
                                                     % (i.e. what value they are voting for when active)

%% Simulate SDT Experiments


Meta_D_Array = zeros(2,4,2,size(Baseline_Lower_JNDs,2));

% Array to keep a trial-by-trial record of results, so we can do multiple 
% simulations, sampling random half datasets, to see if we can distinguish 
% between skewed and normal experiential PDs based on z-scored HR : FAR 
% slopes... 
Z_score_Array = zeros(4,N_Repeats/2,size(Baseline_Lower_JNDs,2));

Decoded_Vals=[]; % Create an array to store values decoded from unadapted population code
Adapted_Decoded_Vals = []; % Create a second array to store values decoded from adapted population codes

Crit_Noise = normrnd(0,0,1,N_Repeats*size(Baseline_Lower_JNDs,2));

Norm_Baseline_Low_Dist = normrnd(Lower_Input,Baseline_JND,1,N_Repeats/2);
Norm_Baseline_High_Dist = normrnd(Upper_Input,Baseline_JND,1,N_Repeats/2);
Norm_Baseline_Dists = Norm_Baseline_Low_Dist;
Norm_Baseline_Dists(2,:)=Norm_Baseline_High_Dist;

Norm_Adapted_Low_Dist = normrnd(Lower_Input,Ad_JND,1,N_Repeats/2);
Norm_Adapted_High_Dist = normrnd(Upper_Input,Ad_JND,1,N_Repeats/2);
Norm_Adapted_Dists = Norm_Adapted_Low_Dist;
Norm_Adapted_Dists(2,:) = Norm_Adapted_High_Dist;

Normal_Meta_D_Array = Meta_D_Array;

%% Prepare Baseline and Adapted channels...

%% Parameter Search Iterations
Vote_Labels = Chn_Labels+180;
Adapted_Labels=Chn_Labels; % Initial labels for each channel

% Identify which channel is tuned to 0 - which we have used as the
% 'adapted' channel...
Ad_Idx = find(Xs==min(abs(Xs)))-1;

% Now we create an array of indices that we use to determine how
% much to reduce the response potential of each model channel, and
% how much (and in which direction) to shift each model channels
% tuning by for this adaptor...
Ad_Index(1:Population_N) =  1 + mod( Ad_Idx+floor((Chn_Labels(1:Population_N)-40)/Channel_Spacing) , Population_N);

% Reduce this channel's response potential by the reduction gain
% for this channel after adapting to this adaptor...
Adapt_Mag(1:Population_N) = Post_Ad_Gain_Reductions(Ad_Index);
Shift_Mag(1:Population_N) = (Tuning_Shift_Multiples(Ad_Index)*Shift_Amplitude);


% If we are simulating trial-by-trial encoding noise generate a
% random gain multiple for each model channel
Channel_Tuning_Noise(1:Population_N) = rand(1,Population_N);
for Ch_Num = 1 : Population_N 

    % Generate baseline model channels - normal probability density
    % functions, centered on different physical test values
    % (Chn_Labels) with a standard deviation (sigma)
    Baseline_Channels(Ch_Num,:) = normpdf(x,Chn_Labels(Ch_Num),sigma);

    % Make sure that the baseline channels start scaled 0 to 1
    Baseline_Channels(Ch_Num,:) = Baseline_Channels(Ch_Num,:)./max(Baseline_Channels(Ch_Num,:));
    
    % Generate adapted model channels - normal probability density
    % functions, centered on different physical test values
    % (Chn_Labels) with a standard deviation (sigma)
    Adapted_Channels(Ch_Num,:) = normpdf(x,Chn_Labels(Ch_Num)+Shift_Mag(Ch_Num),sigma);

    % Make sure that the adapted channels start scaled 0 to 1
    Adapted_Channels(Ch_Num,:) = Adapted_Channels(Ch_Num,:)./max(Adapted_Channels(Ch_Num,:));

    % Reduce channel gains, to simulate adaptation. Mitigate this
    % impact if the adaptor is matched to the average test value
    Adapted_Channels(Ch_Num,:) = Adapted_Channels(Ch_Num,:).*(1-Adapt_Mag(Ch_Num)*Ad_Amplitude*(1-Attenuate_0_Adaptor*0.2));

end

 


close all
    
Trial_Counts = [0 0];
Crit_Noise_Count = 0;
% Begin simulated experiment... 
for Trial = 1 : N_Repeats   

    for Adjust = 1 : size(Baseline_Lower_JNDs,2)
        
        Crit_Noise_Count=Crit_Noise_Count+1;
        
        % Simulating trial-by-trial encoding noise by generating a
        % random gain multiple for each model channel
       Channel_Tuning_Noise(1:Population_N) = rand(1,Population_N);

        Norm_Baseline_Low_Dist = normrnd(Lower_Input,Baseline_JND,1,N_Repeats/2);
        Norm_Baseline_High_Dist = normrnd(Upper_Input,Baseline_JND,1,N_Repeats/2);
        Norm_Baseline_Dists = Norm_Baseline_Low_Dist;
        Norm_Baseline_Dists(2,:)=Norm_Baseline_High_Dist;

        Norm_Adapted_Low_Dist = normrnd(Lower_Input,Ad_JND,1,N_Repeats/2);
        Norm_Adapted_High_Dist = normrnd(Upper_Input,Ad_JND,1,N_Repeats/2);
        Norm_Adapted_Dists = Norm_Adapted_Low_Dist;
        Norm_Adapted_Dists(2,:) = Norm_Adapted_High_Dist;
       
       for Ch_Num = 1 : Population_N 
            % Now multiply the channel's gain potential by the encoding noise 
            % level set for this channel on this trial...
            Trial_Baseline_Channels_1(Ch_Num,:) = Baseline_Channels(Ch_Num,:)*(Channel_Tuning_Noise(Ch_Num));
            
            % Now multiply the channel's gain potential by encoding noise 
            % (same noise levels as for baseline)
            Trial_Adapted_Channels_1(Ch_Num,:) = Adapted_Channels(Ch_Num,:)*(Channel_Tuning_Noise(Ch_Num));
       end
    
        if mod(Trial,2) == 1
            % Sample Low Test values
            Baseline_Test = Baseline_Lower_JNDs(Adjust);
            Adjust_Normal_Baseline_Dists = Baseline_Test-Baseline_Lower_JNDs(1);
            Results_Row = 1;
        else
            % Sample High Test values
            Baseline_Test = Baseline_Upper_JNDs(Adjust);
            Adjust_Normal_Baseline_Dists = Baseline_Test-Baseline_Upper_JNDs(1);
            Results_Row = 2;
        end
        
        
        if Adjust == 1
            Trial_Counts(Results_Row)=Trial_Counts(Results_Row)+1;
        end


        % Find the position on X axis that corresponds with the Test for this trial...
        Test_Differences = abs(x-Baseline_Test);
        [val Test_idx] = min(Test_Differences);

        Votes = [];     % Prepare an empty array to store channel votes in

        % Calculate model encoding, by taking a weighted sum of channel 'votes'
        % (e.g. channel responses to the test). Channel labels are multiplied 
        % by channel response magnitudes (which are divided by the total
        % response of ALL channels)
        Votes(1:Population_N) = Vote_Labels(:).*((Trial_Baseline_Channels_1(:,Test_idx))./sum(Trial_Baseline_Channels_1(:,Test_idx)));

        Decoded_Val = sum(Votes)-180; 
        Base_Decoded_Vals(Results_Row,Trial_Counts(Results_Row),Adjust) = sum(Votes)-180;

        % Categorise input as Negative or Positive - by recording when trials
        % have been encoded as a positive value
        if Decoded_Val > 0 %Crit_Noise(Crit_Noise_Count)%Decoded_Val_2
            Category_Resp = 1;
        elseif Decoded_Val < 0 %Crit_Noise(Crit_Noise_Count)%Decoded_Val_2
            Category_Resp = 0;
        else
            Category_Resp = round(rand(1));
        end

        % Categorise confidence as low or high, relative to the confidence
        % criterion (the unsigned encoded value needed to elicit a high-level of 
        % confidence) by recording when trials have exceeded the criterion 
        if abs(Decoded_Val) >= Confidence_Crit %| Decoded_Val >= 0+Confidence_Crit
            Confidence_Resp = 1;
        else
            Confidence_Resp = 0;
        end

        if Category_Resp == 1
            if Confidence_Resp == 1
                Meta_D_Array(Results_Row,1,1,Adjust)=Meta_D_Array(Results_Row,1,1,Adjust)+1;
            else
                Meta_D_Array(Results_Row,2,1,Adjust)=Meta_D_Array(Results_Row,2,1,Adjust)+1;
            end
        else
            if Confidence_Resp == 1
                Meta_D_Array(Results_Row,4,1,Adjust)=Meta_D_Array(Results_Row,4,1,Adjust)+1;
            else
                Meta_D_Array(Results_Row,3,1,Adjust)=Meta_D_Array(Results_Row,3,1,Adjust)+1;
            end
        end
        Z_score_Array_Baseline(Results_Row+2,Trial_Counts(Results_Row),Adjust)=Category_Resp;
        
        %%        
        % Categorise input as Negative or Positive - by recording when trials
        % have been encoded as a positive value
        if Norm_Baseline_Dists(Results_Row,Trial_Counts(Results_Row))+Adjust_Normal_Baseline_Dists > 0
            Category_Resp = 1;
        elseif Norm_Baseline_Dists(Results_Row,Trial_Counts(Results_Row))+Adjust_Normal_Baseline_Dists < 0
            Category_Resp = 0;
        else
            Category_Resp = round(rand(1));
        end
        Base_Norm_Decoded_Vals(Results_Row,Trial_Counts(Results_Row),Adjust)=Norm_Baseline_Dists(Results_Row,Trial_Counts(Results_Row))+Adjust_Normal_Baseline_Dists;


        % Categorise confidence as low or high, relative to the confidence
        % criterion (the unsigned encoded value needed to elicit a high-level of 
        % confidence) by recording when trials have exceeded the criterion 
        if abs(Norm_Baseline_Dists(Results_Row,Trial_Counts(Results_Row))+Adjust_Normal_Baseline_Dists) >= Confidence_Crit % | Norm_Baseline_Dists(Results_Row,Trial_Counts(Results_Row))+Adjust_Normal_Baseline_Dists >= 0+Confidence_Crit
            Confidence_Resp = 1;
        else
            Confidence_Resp = 0;
        end

        if Category_Resp == 1
            if Confidence_Resp == 1
                Normal_Meta_D_Array(Results_Row,1,1,Adjust)=Normal_Meta_D_Array(Results_Row,1,1,Adjust)+1;
            else
                Normal_Meta_D_Array(Results_Row,2,1,Adjust)=Normal_Meta_D_Array(Results_Row,2,1,Adjust)+1;
            end
        else
            if Confidence_Resp == 1
                Normal_Meta_D_Array(Results_Row,4,1,Adjust)=Normal_Meta_D_Array(Results_Row,4,1,Adjust)+1;
            else
                Normal_Meta_D_Array(Results_Row,3,1,Adjust)=Normal_Meta_D_Array(Results_Row,3,1,Adjust)+1;
            end
        end
        Z_score_Array_Baseline(Results_Row+2,Trial_Counts(Results_Row),Adjust)=Category_Resp;
    
        %%
        
        if mod(Trial,2) == 1
            Adapted_Test = Ad_Lower_JNDs(Adjust);
            Adjust_Normal_Ad_Dists = Adapted_Test-Ad_Lower_JNDs(1);
        else
            Adapted_Test = Ad_Upper_JNDs(Adjust);
            Adjust_Normal_Ad_Dists = Adapted_Test-Ad_Upper_JNDs(1);
        end
        Test_Differences = abs(x-Adapted_Test);
        [val Test_idx] = min(Test_Differences);

        Adapted_Votes = [];     % Prepare an empty array to store channel votes in

        % Calculate adapted model encoding, by taking a weighted sum of 
        % channel 'votes' (e.g. adapted channel responses to the test). 
        % Channel labels are multiplied by channel response magnitudes 
        % (which are divided by the total response of ALL channels)
        Adapted_Votes(1:Population_N) = Vote_Labels(:).*((Trial_Adapted_Channels_1(:,Test_idx))./sum(Trial_Adapted_Channels_1(:,Test_idx)));

        Ad_Decoded_Vals(Results_Row,Trial_Counts(Results_Row),Adjust) = sum(Adapted_Votes)-180;
        Adapted_Decoded_Val = sum(Adapted_Votes)-180;

        % Categorise input as Negative or Positive - by recording when trials
        % have been encoded as a positive value
        if Adapted_Decoded_Val > 0
            Category_Resp=1;
        elseif Adapted_Decoded_Val < 0
            Category_Resp=0;
        else
            Category_Resp = round(rand(1));
        end

        % Categorise confidence as low or high, relative to the confidence
        % criterion (the unsigned encoded value needed to elicit a high-level of 
        % confidence) by recording when trials have exceeded the criterion 
        if abs(Adapted_Decoded_Val) >= Confidence_Crit %| Adapted_Decoded_Val >= 0+Confidence_Crit
            Confidence_Resp=1;
        else
            Confidence_Resp=0;
        end

        if Category_Resp == 1
            if Confidence_Resp == 1
                Meta_D_Array(Results_Row,1,2,Adjust)=Meta_D_Array(Results_Row,1,2,Adjust)+1;
            else
                Meta_D_Array(Results_Row,2,2,Adjust)=Meta_D_Array(Results_Row,2,2,Adjust)+1;
            end
        elseif Adapted_Decoded_Val < 0
            if Confidence_Resp == 1
                Meta_D_Array(Results_Row,4,2,Adjust)=Meta_D_Array(Results_Row,4,2,Adjust)+1;
            else
                Meta_D_Array(Results_Row,3,2,Adjust)=Meta_D_Array(Results_Row,3,2,Adjust)+1;
            end
        end
        Z_score_Array(Results_Row,Trial_Counts(Results_Row),Adjust)=Category_Resp;

        %%
        % Categorise input as Negative or Positive - by recording when trials
        % have been encoded as a positive value
        if Norm_Adapted_Dists(Results_Row,Trial_Counts(Results_Row))+Adjust_Normal_Ad_Dists > 0
            Category_Resp=1;
        elseif Norm_Adapted_Dists(Results_Row,Trial_Counts(Results_Row))+Adjust_Normal_Ad_Dists < 0
            Category_Resp=0;
        else
            Category_Resp = round(rand(1));
        end
        Ad_Norm_Decoded_Vals(Results_Row,Trial_Counts(Results_Row),Adjust)=Norm_Adapted_Dists(Results_Row,Trial_Counts(Results_Row))+Adjust_Normal_Ad_Dists;

        % Categorise confidence as low or high, relative to the confidence
        % criterion (the unsigned encoded value needed to elicit a high-level of 
        % confidence) by recording when trials have exceeded the criterion 
        if abs(Norm_Adapted_Dists(Results_Row,Trial_Counts(Results_Row))+Adjust_Normal_Ad_Dists) >= Confidence_Crit % | Norm_Adapted_Dists(Results_Row,Trial_Counts(Results_Row))+Adjust_Normal_Ad_Dists >= 0+Confidence_Crit
            Confidence_Resp=1;
        else
            Confidence_Resp=0;
        end

        if Category_Resp == 1
            if Confidence_Resp == 1
                Normal_Meta_D_Array(Results_Row,1,2,Adjust)=Normal_Meta_D_Array(Results_Row,1,2,Adjust)+1;
            else
                Normal_Meta_D_Array(Results_Row,2,2,Adjust)=Normal_Meta_D_Array(Results_Row,2,2,Adjust)+1;
            end
        else
            if Confidence_Resp == 1
                Normal_Meta_D_Array(Results_Row,4,2,Adjust)=Normal_Meta_D_Array(Results_Row,4,2,Adjust)+1;
            else
                Normal_Meta_D_Array(Results_Row,3,2,Adjust)=Normal_Meta_D_Array(Results_Row,3,2,Adjust)+1;
            end
        end
        Z_score_Array(Results_Row+2,Trial_Counts(Results_Row),Adjust)=Category_Resp;
            
    end
        
    if mod(Trial,100) == 0
        Pr_Experiment_Completed = Trial/N_Repeats
    end
end


%%
Shape_Params=[];

Rounded_Baseline_Vals = round(Base_Decoded_Vals(:,:,1),2);
Baseline_Means=mean(Rounded_Baseline_Vals,2);

The_Min_Min = min((min(Rounded_Baseline_Vals(1,:))),(min(Rounded_Baseline_Vals(2,:))));

figure(1)
subplot(2,1,2)
box off
histfit(Rounded_Baseline_Vals(1,:)+abs(The_Min_Min),100,'Gamma');
hold on
histfit(Rounded_Baseline_Vals(2,:)+abs(The_Min_Min),100,'Gamma');

h = findobj(gca);
h(2).Color=[0 0.5 0];
h(3).FaceColor='g';
h(3).EdgeColor='none';
h(3).FaceAlpha=0.5;

h(4).Color=[0.75 0 0];
h(5).FaceColor='r';
h(5).EdgeColor='none';
h(5).FaceAlpha=0.5;

h(3).XData = h(3).XData+The_Min_Min;
h(2).XData = h(2).XData+The_Min_Min;
h(5).XData = h(5).XData+The_Min_Min;
h(4).XData = h(4).XData+The_Min_Min;

ax=gca;
plot([0,0],[0,ax.YLim(2)],'-k','LineWidth',2)
plot([-Confidence_Crit,-Confidence_Crit],[0,ax.YLim(2)],'-b','LineWidth',2)
plot([Confidence_Crit,Confidence_Crit],[0,ax.YLim(2)],'-b','LineWidth',2)

TheLimits_1 = xlim;

Popn = Rounded_Baseline_Vals(1,:);
Difference = 100-mean(Popn);
Popn=Popn+Difference;
Popn=Popn';

n = size(Popn,1);  % number of observations
n1 = 1/n; % used with moments in most equations 
ma = mean(Popn(:,1)); % mean

% Moments 
Popn(:,2) = (Popn(:,1)-ma).^2;
Popn(:,3) = (Popn(:,1)-ma).^3;
Popn(:,4) = (Popn(:,1)-ma).^4;
m2 = sum(Popn(:,2));
m3 = sum(Popn(:,3));
m4 = sum(Popn(:,4));
% Skewness
skew = (n1*m3)/((n1*m2)^(3/2));
skg1 = @(Popn)((1/numel(Popn))*sum((Popn-mean(Popn)).^3))/(((1/numel(Popn))*sum((Popn-mean(Popn)).^2))^(3/2));
skg1_bootci = bootci(1000, skg1, Popn(:,1));
Shape_Params(1,1) = skew;
Shape_Params(1,2) = skg1_bootci(1);
Shape_Params(1,3) = skg1_bootci(2);
% Kurtosis
kurt = (sum((Popn-mean(Popn)).^4)./length(Popn)) ./ (var(Popn,1).^2);
kur1 = @(Popn)((1/numel(Popn))*sum((Popn-mean(Popn)).^4))/(((1/numel(Popn))*sum((Popn-mean(Popn)).^2))^(4/2));
kur1_bootci = bootci(1000, kur1, Popn(:,1));
Shape_Params(1,4) = kurt(1);
Shape_Params(1,5) = kur1_bootci(1);
Shape_Params(1,6) = kur1_bootci(2);

Popn = Rounded_Baseline_Vals(2,:);
Difference = 100-mean(Popn);
Popn=Popn+Difference;
Popn=Popn';

ma = mean(Popn(:,1)); % mean
% Moments 
Popn(:,2) = (Popn(:,1)-ma).^2;
Popn(:,3) = (Popn(:,1)-ma).^3;
Popn(:,4) = (Popn(:,1)-ma).^4;
m2 = sum(Popn(:,2));
m3 = sum(Popn(:,3));
m4 = sum(Popn(:,4));
% Skewness
skew = (n1*m3)/((n1*m2)^(3/2));
skg1 = @(Popn)((1/numel(Popn))*sum((Popn-mean(Popn)).^3))/(((1/numel(Popn))*sum((Popn-mean(Popn)).^2))^(3/2));
skg1_bootci = bootci(1000, skg1, Popn(:,1));
Shape_Params(2,1) = skew;
Shape_Params(2,2) = skg1_bootci(1);
Shape_Params(2,3) = skg1_bootci(2);
% Kurtosis
kurt = (sum((Popn-mean(Popn)).^4)./length(Popn)) ./ (var(Popn,1).^2);
kur1 = @(Popn)((1/numel(Popn))*sum((Popn-mean(Popn)).^4))/(((1/numel(Popn))*sum((Popn-mean(Popn)).^2))^(4/2));
kur1_bootci = bootci(1000, kur1, Popn(:,1));
Shape_Params(2,4) = kurt(1);
Shape_Params(2,5) = kur1_bootci(1);
Shape_Params(2,6) = kur1_bootci(2);

Baseline_Test = round(Baseline_Lower_JNDs(1),2);
Adjust_Normal_Baseline_Lower = Baseline_Test-Baseline_Lower_JNDs(round(size(Baseline_Lower_JNDs,2)/2));
Baseline_Test = round(Baseline_Upper_JNDs(1),2);
Adjust_Normal_Baseline_Upper = Baseline_Test-Baseline_Upper_JNDs(round(size(Baseline_Lower_JNDs,2)/2));

Norm_Baseline_Low_Dist=Norm_Baseline_Low_Dist+Adjust_Normal_Baseline_Lower;
Norm_Baseline_High_Dist=Norm_Baseline_High_Dist+Adjust_Normal_Baseline_Upper;

subplot(2,1,1)
box off
histfit(Norm_Baseline_Low_Dist,100,'normal');
hold on
histfit(Norm_Baseline_High_Dist,100,'normal');

h = findobj(gca);
h(2).Color=[0 0.5 0];
h(3).FaceColor='g';
h(3).EdgeColor='none';
h(3).FaceAlpha=0.5;

h(4).Color=[0.75 0 0];
h(5).FaceColor='r';
h(5).EdgeColor='none';
h(5).FaceAlpha=0.5;

ax=gca;
plot([0,0],[0,ax.YLim(2)],'-k','LineWidth',2)
plot([-Confidence_Crit,-Confidence_Crit],[0,ax.YLim(2)],'-b','LineWidth',2)
plot([Confidence_Crit,Confidence_Crit],[0,ax.YLim(2)],'-b','LineWidth',2)

TheLimits_2 = xlim;

Low_Limit = min([TheLimits_1(1) TheLimits_2(1)]);
High_Limit = max([TheLimits_1(2) TheLimits_2(2)]);

xlim([Low_Limit High_Limit]);

subplot(2,1,2)
xlim([Low_Limit High_Limit]);

Popn = Norm_Baseline_Low_Dist;
Difference = 100-mean(Popn);
Popn=Popn+Difference;
Popn=Popn';

ma = mean(Popn(:,1)); % mean
% Moments 
Popn(:,2) = (Popn(:,1)-ma).^2;
Popn(:,3) = (Popn(:,1)-ma).^3;
Popn(:,4) = (Popn(:,1)-ma).^4;
m2 = sum(Popn(:,2));
m3 = sum(Popn(:,3));
m4 = sum(Popn(:,4));
% Skewness
skew = (n1*m3)/((n1*m2)^(3/2));
skg1 = @(Popn)((1/numel(Popn))*sum((Popn-mean(Popn)).^3))/(((1/numel(Popn))*sum((Popn-mean(Popn)).^2))^(3/2));
skg1_bootci = bootci(1000, skg1, Popn(:,1));
Shape_Params(3,1) = skew;
Shape_Params(3,2) = skg1_bootci(1);
Shape_Params(3,3) = skg1_bootci(2);
% Kurtosis
kurt = (sum((Popn-mean(Popn)).^4)./length(Popn)) ./ (var(Popn,1).^2);
kur1 = @(Popn)((1/numel(Popn))*sum((Popn-mean(Popn)).^4))/(((1/numel(Popn))*sum((Popn-mean(Popn)).^2))^(4/2));
kur1_bootci = bootci(1000, kur1, Popn(:,1));
Shape_Params(3,4) = kurt(1);
Shape_Params(3,5) = kur1_bootci(1);
Shape_Params(3,6) = kur1_bootci(2);

Popn = Norm_Baseline_High_Dist;
Difference = 100-mean(Popn);
Popn=Popn+Difference;
Popn=Popn';
ma = mean(Popn(:,1)); % mean
% Moments 
Popn(:,2) = (Popn(:,1)-ma).^2;
Popn(:,3) = (Popn(:,1)-ma).^3;
Popn(:,4) = (Popn(:,1)-ma).^4;
m2 = sum(Popn(:,2));
m3 = sum(Popn(:,3));
m4 = sum(Popn(:,4));
% Skewness
skew = (n1*m3)/((n1*m2)^(3/2));
skg1 = @(Popn)((1/numel(Popn))*sum((Popn-mean(Popn)).^3))/(((1/numel(Popn))*sum((Popn-mean(Popn)).^2))^(3/2));
skg1_bootci = bootci(1000, skg1, Popn(:,1));
Shape_Params(4,1) = skew;
Shape_Params(4,2) = skg1_bootci(1);
Shape_Params(4,3) = skg1_bootci(2);
% Kurtosis
kurt = (sum((Popn-mean(Popn)).^4)./length(Popn)) ./ (var(Popn,1).^2);
kur1 = @(Popn)((1/numel(Popn))*sum((Popn-mean(Popn)).^4))/(((1/numel(Popn))*sum((Popn-mean(Popn)).^2))^(4/2));
kur1_bootci = bootci(1000, kur1, Popn(:,1));
Shape_Params(4,4) = kurt(1);
Shape_Params(4,5) = kur1_bootci(1);
Shape_Params(4,6) = kur1_bootci(2);

%%
figure(2)
subplot(2,1,2)
box off
Rounded_Adapted_Vals = round(Ad_Decoded_Vals(:,:,1),2);

Adapted_Means=mean(Rounded_Adapted_Vals,2);

TheMin = min(min(Rounded_Adapted_Vals(1,:)),min(Rounded_Adapted_Vals(2,:)))

histfit(Rounded_Adapted_Vals(1,:)+abs(TheMin),100,'gamma');
hold on
histfit(Rounded_Adapted_Vals(2,:)+abs(TheMin),100,'gamma');
h = findobj(gca);
h(3).FaceColor='g';
h(3).EdgeColor='none';
h(3).FaceAlpha=0.5;
h(2).Color=[0 0.5 0];
h(5).FaceColor='r';
h(5).EdgeColor='none';
h(5).FaceAlpha=0.5;
h(4).Color=[0.75 0 0];

h(3).XData = h(3).XData+TheMin;
h(2).XData = h(2).XData+TheMin;
h(5).XData = h(5).XData+TheMin;
h(4).XData = h(4).XData+TheMin;

ax=gca;
plot([0,0],[0,ax.YLim(2)],'-k','LineWidth',2)
plot([-Confidence_Crit,-Confidence_Crit],[0,ax.YLim(2)],'-b','LineWidth',2)
plot([Confidence_Crit,Confidence_Crit],[0,ax.YLim(2)],'-b','LineWidth',2)

TheLimits_1 = xlim;

Adapted_Test = round(Ad_Lower_JNDs(1),2);
Adjust_Normal_Ad_Lower = Adapted_Test-Ad_Lower_JNDs(round(size(Baseline_Lower_JNDs,2)/2));
Adapted_Test = round(Ad_Upper_JNDs(1),2);
Adjust_Normal_Ad_Upper = Adapted_Test-Ad_Upper_JNDs(round(size(Baseline_Lower_JNDs,2)/2));

Norm_Adapted_Low_Dist=Norm_Adapted_Low_Dist+Adjust_Normal_Ad_Lower;
Norm_Adapted_High_Dist=Norm_Adapted_High_Dist+Adjust_Normal_Ad_Upper;

subplot(2,1,1)
box off
histfit(Norm_Adapted_Low_Dist,100,'normal');
hold on
histfit(Norm_Adapted_High_Dist,100,'normal');
h = findobj(gca);
h(2).Color=[0 0.5 0];
h(3).FaceColor='g';
h(3).EdgeColor='none';
h(3).FaceAlpha=0.5;

h(4).Color=[0.75 0 0];
h(5).FaceColor='r';
h(5).EdgeColor='none';
h(5).FaceAlpha=0.5;

ax=gca;
plot([0,0],[0,ax.YLim(2)],'-k','LineWidth',2)
plot([-Confidence_Crit,-Confidence_Crit],[0,ax.YLim(2)],'-b','LineWidth',2)
plot([Confidence_Crit,Confidence_Crit],[0,ax.YLim(2)],'-b','LineWidth',2)

TheLimits_2 = xlim;

Low_Limit = min([TheLimits_1(1) TheLimits_2(1)]);
High_Limit = max([TheLimits_1(2) TheLimits_2(2)]);

Popn = Rounded_Adapted_Vals(1,:);
Difference = 100-mean(Popn);
Popn=Popn+Difference;
Popn=Popn';

ma = mean(Popn(:,1)); % mean
% Moments 
Popn(:,2) = (Popn(:,1)-ma).^2;
Popn(:,3) = (Popn(:,1)-ma).^3;
Popn(:,4) = (Popn(:,1)-ma).^4;
m2 = sum(Popn(:,2));
m3 = sum(Popn(:,3));
m4 = sum(Popn(:,4));
% Skewness
skew = (n1*m3)/((n1*m2)^(3/2));
skg1 = @(Popn)((1/numel(Popn))*sum((Popn-mean(Popn)).^3))/(((1/numel(Popn))*sum((Popn-mean(Popn)).^2))^(3/2));
skg1_bootci = bootci(1000, skg1, Popn(:,1));
Shape_Params(5,1) = skew;
Shape_Params(5,2) = skg1_bootci(1);
Shape_Params(5,3) = skg1_bootci(2);
% Kurtosis
kurt = (sum((Popn-mean(Popn)).^4)./length(Popn)) ./ (var(Popn,1).^2);
kur1 = @(Popn)((1/numel(Popn))*sum((Popn-mean(Popn)).^4))/(((1/numel(Popn))*sum((Popn-mean(Popn)).^2))^(4/2));
kur1_bootci = bootci(1000, kur1, Popn(:,1));
Shape_Params(5,4) = kurt(1);
Shape_Params(5,5) = kur1_bootci(1)
Shape_Params(5,6) = kur1_bootci(2)


Popn = Rounded_Adapted_Vals(2,:);
Difference = 100-mean(Popn);
Popn=Popn+Difference;
Popn=Popn';
ma = mean(Popn(:,1)); % mean
% Moments 
Popn(:,2) = (Popn(:,1)-ma).^2;
Popn(:,3) = (Popn(:,1)-ma).^3;
Popn(:,4) = (Popn(:,1)-ma).^4;
m2 = sum(Popn(:,2));
m3 = sum(Popn(:,3));
m4 = sum(Popn(:,4));
% Skewness
skew = (n1*m3)/((n1*m2)^(3/2));
skg1 = @(Popn)((1/numel(Popn))*sum((Popn-mean(Popn)).^3))/(((1/numel(Popn))*sum((Popn-mean(Popn)).^2))^(3/2));
skg1_bootci = bootci(1000, skg1, Popn(:,1));
Shape_Params(6,1) = skew;
Shape_Params(6,2) = skg1_bootci(1);
Shape_Params(6,3) = skg1_bootci(2);
% Kurtosis
kurt = (sum((Popn-mean(Popn)).^4)./length(Popn)) ./ (var(Popn,1).^2);
kur1 = @(Popn)((1/numel(Popn))*sum((Popn-mean(Popn)).^4))/(((1/numel(Popn))*sum((Popn-mean(Popn)).^2))^(4/2));
kur1_bootci = bootci(1000, kur1, Popn(:,1));
Shape_Params(6,4) = kurt(1);
Shape_Params(6,5) = kur1_bootci(1);
Shape_Params(6,6) = kur1_bootci(2);


Popn=Norm_Adapted_Low_Dist;
Difference = 100-mean(Popn);
Popn=Popn+Difference;
Popn=Popn';

ma = mean(Popn(:,1)); % mean
% Moments 
Popn(:,2) = (Popn(:,1)-ma).^2;
Popn(:,3) = (Popn(:,1)-ma).^3;
Popn(:,4) = (Popn(:,1)-ma).^4;
m2 = sum(Popn(:,2));
m3 = sum(Popn(:,3));
m4 = sum(Popn(:,4));
% Skewness
skew = (n1*m3)/((n1*m2)^(3/2));
skg1 = @(Popn)((1/numel(Popn))*sum((Popn-mean(Popn)).^3))/(((1/numel(Popn))*sum((Popn-mean(Popn)).^2))^(3/2));
skg1_bootci = bootci(1000, skg1, Popn(:,1));
Shape_Params(7,1) = skew;
Shape_Params(7,2) = skg1_bootci(1);
Shape_Params(7,3) = skg1_bootci(2);
% Kurtosis
kurt = (sum((Popn-mean(Popn)).^4)./length(Popn)) ./ (var(Popn,1).^2);
kur1 = @(Popn)((1/numel(Popn))*sum((Popn-mean(Popn)).^4))/(((1/numel(Popn))*sum((Popn-mean(Popn)).^2))^(4/2));
kur1_bootci = bootci(1000, kur1, Popn(:,1));
Shape_Params(7,4) = kurt(1);
Shape_Params(7,5) = kur1_bootci(1);
Shape_Params(7,6) = kur1_bootci(2);


Popn=Norm_Adapted_High_Dist;
Difference = 100-mean(Popn);
Popn=Popn+Difference;
Popn=Popn';

ma = mean(Popn(:,1)); % mean
% Moments 
Popn(:,2) = (Popn(:,1)-ma).^2;
Popn(:,3) = (Popn(:,1)-ma).^3;
Popn(:,4) = (Popn(:,1)-ma).^4;
m2 = sum(Popn(:,2));
m3 = sum(Popn(:,3));
m4 = sum(Popn(:,4));
% Skewness
skew = (n1*m3)/((n1*m2)^(3/2));
skg1 = @(Popn)((1/numel(Popn))*sum((Popn-mean(Popn)).^3))/(((1/numel(Popn))*sum((Popn-mean(Popn)).^2))^(3/2));
skg1_bootci = bootci(1000, skg1, Popn(:,1));
Shape_Params(8,1) = skew;
Shape_Params(8,2) = skg1_bootci(1);
Shape_Params(8,3) = skg1_bootci(2)
% Kurtosis
kurt = (sum((Popn-mean(Popn)).^4)./length(Popn)) ./ (var(Popn,1).^2);
kur1 = @(Popn)((1/numel(Popn))*sum((Popn-mean(Popn)).^4))/(((1/numel(Popn))*sum((Popn-mean(Popn)).^2))^(4/2));
kur1_bootci = bootci(1000, kur1, Popn(:,1));
Shape_Params(8,4) = kurt(1);
Shape_Params(8,5) = kur1_bootci(1);
Shape_Params(8,6) = kur1_bootci(2);

    
%% Meta D-Prime Analyses...

% To prevent analyses being based on 0 values...
% SDT_Adjust = 0;%1/N_Repeats;
% Meta_D_Array = Meta_D_Array+SDT_Adjust;
% Normal_Meta_D_Array = Normal_Meta_D_Array+SDT_Adjust;

Analysis=[];
dPrimes=[];
Meta_dPrimes = [];
Meta_d_Ratio_Corrs = [];
Norm_Meta_d_Ratio_Corrs = [];
for t = 1 : size(Meta_D_Array,3)
    for SDT_Adjust = 1 : size(Meta_D_Array,4)
        
        Analysis = fit_meta_d_MLE(Meta_D_Array(2,:,t,SDT_Adjust),Meta_D_Array(1,:,t,SDT_Adjust),1);
        dPrimes(t,SDT_Adjust)=Analysis.da;
        Meta_dPrimes(t,SDT_Adjust)=Analysis.meta_da;
        
        if t == 1
            The_Skew_1 = skewness(Base_Decoded_Vals(2,:,SDT_Adjust));
            The_Skew_2 = skewness(Base_Decoded_Vals(1,:,SDT_Adjust));
            The_Kurtosis_1 = kurtosis(Base_Decoded_Vals(2,:,SDT_Adjust))-3;
            The_Kurtosis_2 = kurtosis(Base_Decoded_Vals(1,:,SDT_Adjust))-3;
        else
            The_Skew_1 = skewness(Ad_Decoded_Vals(2,:,SDT_Adjust));
            The_Skew_2 = skewness(Ad_Decoded_Vals(1,:,SDT_Adjust));
            The_Kurtosis_1 = kurtosis(Ad_Decoded_Vals(2,:,SDT_Adjust))-3;
            The_Kurtosis_2 = kurtosis(Ad_Decoded_Vals(1,:,SDT_Adjust))-3;
        end
        
        Meta_d_Ratio_Corrs(t,SDT_Adjust,1) = (Analysis.meta_da / Analysis.da);
        Meta_d_Ratio_Corrs(t,SDT_Adjust,2) = mean([The_Skew_1  The_Skew_2]);
        Meta_d_Ratio_Corrs(t,SDT_Adjust,3) = mean([The_Kurtosis_1  The_Kurtosis_2]);
        
        Analysis = fit_meta_d_MLE(Normal_Meta_D_Array(2,:,t,SDT_Adjust),Normal_Meta_D_Array(1,:,t,SDT_Adjust),1);
        Normal_dPrimes(t,SDT_Adjust)=Analysis.da;
        Normal_Meta_dPrimes(t,SDT_Adjust)=Analysis.meta_da;

        if t == 1
            The_Skew_1 = skewness(Base_Norm_Decoded_Vals(2,:,SDT_Adjust));
            The_Skew_2 = skewness(Base_Norm_Decoded_Vals(1,:,SDT_Adjust));
            The_Kurtosis_1 = kurtosis(Base_Norm_Decoded_Vals(2,:,SDT_Adjust))-3;
            The_Kurtosis_2 = kurtosis(Base_Norm_Decoded_Vals(1,:,SDT_Adjust))-3;
        else
            The_Skew_1 = skewness(Ad_Norm_Decoded_Vals(2,:,SDT_Adjust));
            The_Skew_2 = skewness(Ad_Norm_Decoded_Vals(1,:,SDT_Adjust));
            The_Kurtosis_1 = kurtosis(Ad_Norm_Decoded_Vals(2,:,SDT_Adjust))-3;
            The_Kurtosis_2 = kurtosis(Ad_Norm_Decoded_Vals(1,:,SDT_Adjust))-3;
        end
        Meta_d_Ratio_Corrs(t,size(Meta_D_Array,4)+SDT_Adjust,1) = (Analysis.meta_da / Analysis.da);
        Meta_d_Ratio_Corrs(t,size(Meta_D_Array,4)+SDT_Adjust,2) = mean([The_Skew_1 - The_Skew_2]);
        Meta_d_Ratio_Corrs(t,size(Meta_D_Array,4)+SDT_Adjust,3) = mean([The_Kurtosis_1  The_Kurtosis_2]);

        Norm_Meta_d_Ratio_Corrs(t,SDT_Adjust,1) = Analysis.meta_da / Analysis.da;
        Norm_Meta_d_Ratio_Corrs(t,SDT_Adjust,2) = mean([The_Skew_1 - The_Skew_2]);
        Norm_Meta_d_Ratio_Corrs(t,SDT_Adjust,3) = mean([The_Kurtosis_1  The_Kurtosis_2]);
    end    
end
Meta_D_Ratios = Meta_dPrimes./dPrimes;
Normal_Meta_D_Ratios = Normal_Meta_dPrimes./Normal_dPrimes;

%%
TheMin_dPrimes = min(min(dPrimes));
TheMin_Norm_dPrimes = min(min(Normal_dPrimes));
TheMin_MdPrimes = min(min(Meta_dPrimes));
TheMin_Norm_MdPrimes = min(min(Normal_Meta_dPrimes));
TheMin = 0.45;%min([TheMin_dPrimes,TheMin_Norm_dPrimes,TheMin_MdPrimes,TheMin_Norm_MdPrimes]);


TheMax_dPrimes = max(max(dPrimes));
TheMax_Norm_dPrimes = max(max(Normal_dPrimes));
TheMax_MdPrimes = max(max(Meta_dPrimes));
TheMax_Norm_MdPrimes = max(max(Normal_Meta_dPrimes));
TheMax = 1.7;%max([TheMax_dPrimes TheMax_Norm_dPrimes TheMax_MdPrimes TheMax_Norm_MdPrimes]);


figure(3)
clf
subplot(3,2,1)

scatter(Normal_dPrimes(1,:),Normal_Meta_dPrimes(1,:),'k','filled')
hold on
scatter(dPrimes(1,:),Meta_dPrimes(1,:),'r','filled')

xlabel('Base dPrimes','FontSize',16,'FontWeight','bold')  
ylabel('Base meta dPrimes','FontSize',16,'FontWeight','bold')  

Xlimits = xlim;
Ylimits = ylim;

plot([TheMin,TheMax],[TheMin,TheMax],'-g','LineWidth',2)
xlim([TheMin TheMax]);
ylim([TheMin TheMax]);

Xlimits = xlim;
Ylimits = ylim;

subplot(3,2,3)    

[Rval,Pval] = corr(Meta_d_Ratio_Corrs(1,:,1)',Meta_d_Ratio_Corrs(1,:,2)')
scatter(Meta_d_Ratio_Corrs(1,:,1),Meta_d_Ratio_Corrs(1,:,2),'r','filled')
lsline
h4 = lsline
h4.Color = 'r';
h4.LineWidth = 3;
if round(Pval,3) == 0
    FigText = strcat({'R-val = ','R-Sq = ','P-val '},{num2str(round(Rval,2)),num2str(round((Rval*Rval),2)),'< 0.001'})
else
    FigText = strcat({'R-val = ','R-Sq = ','P-val: '},{num2str(round(Rval,2)),num2str(round((Rval*Rval),2)),num2str(round(Pval,2))})
end
text(0.71,-0.03,FigText{1},'Color', 'b','FontSize', 12);
text(0.71,-0.04,FigText{2},'Color', 'b','FontSize', 12);
text(0.71,-0.05,FigText{3},'Color', 'b','FontSize', 12);
xlabel('Base dPrime : meta dPrime','FontSize',16,'FontWeight','bold')  
ylabel('Base Skew Diff.','FontSize',16,'FontWeight','bold')  
hold on
scatter(Norm_Meta_d_Ratio_Corrs(1,:,1),Norm_Meta_d_Ratio_Corrs(1,:,2),'k','filled')


subplot(3,2,5)    

[Rval,Pval] = corr(Meta_d_Ratio_Corrs(1,:,1)',Meta_d_Ratio_Corrs(1,:,3)')
scatter(Meta_d_Ratio_Corrs(1,:,1),Meta_d_Ratio_Corrs(1,:,3),'r','filled')
lsline
h4 = lsline
h4.Color = 'r';
h4.LineWidth = 3;
if round(Pval,3) == 0
    FigText = strcat({'R-val = ','R-Sq = ','P-val '},{num2str(round(Rval,2)),num2str(round((Rval*Rval),2)),'< 0.001'})
else
    FigText = strcat({'R-val = ','R-Sq = ','P-val: '},{num2str(round(Rval,2)),num2str(round((Rval*Rval),2)),num2str(round(Pval,2))})
end
text(0.72,0.1,FigText{1},'Color', 'b','FontSize', 12);
text(0.72,0.025,FigText{2},'Color', 'b','FontSize', 12);
text(0.72,-0.05,FigText{3},'Color', 'b','FontSize', 12);
xlabel('Base dPrime : meta dPrime','FontSize',16,'FontWeight','bold')  
ylabel('Base Avg. Kurtosis','FontSize',16,'FontWeight','bold')  
hold on
scatter(Norm_Meta_d_Ratio_Corrs(1,:,1),Norm_Meta_d_Ratio_Corrs(1,:,3),'k','filled')


subplot(3,2,2)
scatter(Normal_dPrimes(2,:),Normal_Meta_dPrimes(2,:),'k','filled')
hold on
scatter(dPrimes(2,:),Meta_dPrimes(2,:),'r','filled')
xlabel('Ad. dPrimes','FontSize',16,'FontWeight','bold')  
ylabel('Ad. meta dPrimes','FontSize',16,'FontWeight','bold')  

plot([TheMin,TheMax],[TheMin,TheMax],'-g','LineWidth',2)
xlim([TheMin TheMax]);
ylim([TheMin TheMax]);

Xlimits = xlim;
Ylimits = ylim;

subplot(3,2,4)
[Rval,Pval] = corr(Meta_d_Ratio_Corrs(2,:,1)',Meta_d_Ratio_Corrs(2,:,2)')
scatter(Meta_d_Ratio_Corrs(2,:,1),Meta_d_Ratio_Corrs(2,:,2),'r','filled')
h6 = lsline
h6.Color = 'r';
h6.LineWidth = 3;
if round(Pval,3) == 0
    FigText = strcat({'R-val = ','R-Sq = ','P-val '},{num2str(round(Rval,2)),num2str(round((Rval*Rval),2)),'< 0.001'})
else
    FigText = strcat({'R-val = ','R-Sq = ','P-val: '},{num2str(round(Rval,2)),num2str(round((Rval*Rval),2)),num2str(round(Pval,2))})
end
text(0.82,0.05,FigText{1},'Color', 'b','FontSize', 12);
text(0.82,0.025,FigText{2},'Color', 'b','FontSize', 12);
text(0.82,0.0,FigText{3},'Color', 'b','FontSize', 12);
xlabel('Ad. dPrime : meta dPrime','FontSize',16,'FontWeight','bold')  
ylabel('Ad. Skew Diff.','FontSize',16,'FontWeight','bold')  
hold on
scatter(Norm_Meta_d_Ratio_Corrs(2,:,1),Norm_Meta_d_Ratio_Corrs(2,:,2),'k','filled')


subplot(3,2,6)
[Rval,Pval] = corr(Meta_d_Ratio_Corrs(2,:,1)',Meta_d_Ratio_Corrs(2,:,3)')
scatter(Meta_d_Ratio_Corrs(2,:,1),Meta_d_Ratio_Corrs(2,:,3),'r','filled')
h6 = lsline
h6.Color = 'r';
h6.LineWidth = 3;
if round(Pval,3) == 0
    FigText = strcat({'R-val = ','R-Sq = ','P-val '},{num2str(round(Rval,2)),num2str(round((Rval*Rval),2)),'< 0.001'})
else
    FigText = strcat({'R-val = ','R-Sq = ','P-val: '},{num2str(round(Rval,2)),num2str(round((Rval*Rval),2)),num2str(round(Pval,2))})
end
text(0.82,0.15,FigText{1},'Color', 'b','FontSize', 12);
text(0.82,0.075,FigText{2},'Color', 'b','FontSize', 12);
text(0.82,0.0,FigText{3},'Color', 'b','FontSize', 12);
xlabel('Ad. dPrime : meta dPrime','FontSize',16,'FontWeight','bold')  
ylabel('Ad. Avg. Kurtosis','FontSize',16,'FontWeight','bold')  
hold on
scatter(Norm_Meta_d_Ratio_Corrs(2,:,1),Norm_Meta_d_Ratio_Corrs(2,:,3),'k','filled')

%% Determine if d' meta d' ratios have exceeded a 5% difference, and check 
% to see if plots of z-scored Hit and False Alarm rates have a slope that 
% is discernably different from 1:1

Test_Diffs = (Ad_Upper_JNDs-Ad_Lower_JNDs);

Meta_Ratios = Meta_dPrimes./dPrimes;
Meta_Diffs = Meta_dPrimes-dPrimes;

Norm_Meta_Ratios = Normal_Meta_dPrimes./Normal_dPrimes;
Norm_Meta_Diffs = Normal_Meta_dPrimes-Normal_dPrimes;

figure(4)
clf
subplot(1,2,1)
plot([3,5.4],[1,1],'-g','LineWidth',75)
hold on
scatter(Test_Diffs,Norm_Meta_Ratios(1,:),'k','filled')
scatter(Test_Diffs,Meta_Ratios(1,:),'r','filled')

xlabel('Input Differences (degs)','FontSize',16,'FontWeight','bold')  
ylabel('Meta dPrime Ratios - Baseline','FontSize',16,'FontWeight','bold')  
ax=gca;
ax.FontSize = 14; 
box off
xlim([3 5.4])
ylim([0.7 1.05])
yticks([0.75 0.85 0.95 1.05])

subplot(1,2,2)
plot([3,5.4],[1,1],'-g','LineWidth',75)
hold on
scatter(Test_Diffs,Norm_Meta_Ratios(2,:),'k','filled')
scatter(Test_Diffs,Meta_Ratios(2,:),'r','filled')
xlabel('Input Differences (degs)','FontSize',16,'FontWeight','bold')  
ylabel('Meta dPrime Ratios - Adapted','FontSize',16,'FontWeight','bold')
ax=gca;
ax.FontSize = 14; 
box off
xlim([3 5.4])
ylim([0.7 1.05])
yticks([0.75 0.85 0.95 1.05])


%% Analysis to determine if results informed by a skewed distribution could be detected    
[vals Potential_Indicies_Ad] = find(Meta_Ratios(2,:) < 0.95);
[vals Potential_Indicies_Baseline] = find(Meta_Ratios(1,:) < 0.95);

N_Simulations = 1000;
slopes = [];
figure(10)
clf
for trial_N = 1 : N_Simulations

    Random_Trial_Order = randperm(N_Repeats/2);
    Random_Levels_Order_Baseline = randperm(size(Potential_Indicies_Baseline,2));
    Random_Levels_Order_Adapted = randperm(size(Potential_Indicies_Ad,2));

    Data_Baseline = Z_score_Array_Baseline(:,Random_Trial_Order(1:size(Z_score_Array_Baseline,2)/2),Potential_Indicies_Baseline(Random_Levels_Order_Baseline(1:5)));
    Data = Z_score_Array(:,Random_Trial_Order(1:size(Z_score_Array,2)/2),Potential_Indicies_Ad(Random_Levels_Order_Adapted(1:5)));

    SD_FARs = squeeze(mean(Data_Baseline(1,:,:),2));
    SD_HRs= squeeze(mean(Data_Baseline(2,:,:),2));
    Xs = icdf('normal',SD_FARs,0,1);
    Ys = icdf('normal',SD_HRs,0,1);    
    for rows = 1 : 3
        figure(7)
        clf
        scatter(Xs(1:2+rows),Ys(1:2+rows),24,'g','filled');
        hold on
        info = lsline;
        slopes_Baseline(rows,2,trial_N) = (max(info.YData)-min(info.YData)) / (max(info.XData)-min(info.XData));
    end

    SD_FARs = squeeze(mean(Data_Baseline(3,:,:),2));
    SD_HRs= squeeze(mean(Data_Baseline(4,:,:),2));
    Xs = icdf('normal',SD_FARs,0,1);
    Ys = icdf('normal',SD_HRs,0,1);
    for rows = 1 : 3
        figure(7)
        clf
        scatter(Xs(1:2+rows),Ys(1:2+rows),24,'g','filled');
        hold on
        info = lsline;
        slopes_Baseline(rows,1,trial_N) = (max(info.YData)-min(info.YData)) / (max(info.XData)-min(info.XData));
    end

    SD_FARs = squeeze(mean(Data(1,:,:),2));
    SD_HRs= squeeze(mean(Data(2,:,:),2));
    Xs = icdf('normal',SD_FARs,0,1);
    Ys = icdf('normal',SD_HRs,0,1);    
    for rows = 1 : 3
        figure(7)
        clf
        scatter(Xs(1:2+rows),Ys(1:2+rows),24,'g','filled');
        hold on
        info = lsline;
        slopes(rows,2,trial_N) = (max(info.YData)-min(info.YData)) / (max(info.XData)-min(info.XData));
    end

    SD_FARs = squeeze(mean(Data(3,:,:),2));
    SD_HRs= squeeze(mean(Data(4,:,:),2));
    Xs = icdf('normal',SD_FARs,0,1);
    Ys = icdf('normal',SD_HRs,0,1);
    for rows = 1 : 3
        figure(7)
        clf
        scatter(Xs(1:2+rows),Ys(1:2+rows),24,'g','filled');
        hold on
        info = lsline;
        slopes(rows,1,trial_N) = (max(info.YData)-min(info.YData)) / (max(info.XData)-min(info.XData));
    end


end


%%
AVG_Slopes_Baseline = mean(slopes_Baseline,3);
SD_Slopes_Baseline  = std(slopes_Baseline,1,3);

AVG_Slopes = mean(slopes,3);
SD_Slopes  = std(slopes,1,3);

Low_Crit = AVG_Slopes(:,1)-(SD_Slopes(:,1)*3)
High_Crit = AVG_Slopes(:,1)+(SD_Slopes(:,1)*3)

Low_Crit_Baseline = AVG_Slopes_Baseline(:,1)-(SD_Slopes_Baseline(:,1)*3)
High_Crit_Baseline = AVG_Slopes_Baseline(:,1)+(SD_Slopes_Baseline(:,1)*3)

Failed_Slopes = slopes;
Failed_Slopes_Baseline = slopes_Baseline;
for n = 1 : 3
    Temp_Slopes = Failed_Slopes(n,1,:);
    Temp_Slopes(Temp_Slopes<Low_Crit(n))=10;
    Temp_Slopes(Temp_Slopes>High_Crit(n))=10;
    Temp_Slopes(Temp_Slopes<10)=0;
    Temp_Slopes(Temp_Slopes==10)=1;
    Failed_Slopes(n,1,:)=Temp_Slopes;

    Temp_Slopes = Failed_Slopes(n,2,:);
    Temp_Slopes(Temp_Slopes<Low_Crit(n))=10;
    Temp_Slopes(Temp_Slopes>High_Crit(n))=10;
    Temp_Slopes(Temp_Slopes<10)=0;
    Temp_Slopes(Temp_Slopes==10)=1;
    Failed_Slopes(n,2,:)=Temp_Slopes;

    Temp_Slopes = Failed_Slopes_Baseline(n,1,:);
    Temp_Slopes(Temp_Slopes<Low_Crit_Baseline(n))=10;
    Temp_Slopes(Temp_Slopes>High_Crit_Baseline(n))=10;
    Temp_Slopes(Temp_Slopes<10)=0;
    Temp_Slopes(Temp_Slopes==10)=1;
    Failed_Slopes_Baseline(n,1,:)=Temp_Slopes;

    Temp_Slopes = Failed_Slopes(n,2,:);
    Temp_Slopes(Temp_Slopes<Low_Crit_Baseline(n))=10;
    Temp_Slopes(Temp_Slopes>High_Crit_Baseline(n))=10;
    Temp_Slopes(Temp_Slopes<10)=0;
    Temp_Slopes(Temp_Slopes==10)=1;
    Failed_Slopes_Baseline(n,2,:)=Temp_Slopes;
end
Slope_Fail_Rates = mean(Failed_Slopes,3);
Slope_Fail_Rates_Baseline = mean(Failed_Slopes_Baseline,3);
Sample_Pts = [3 4 5];

figure(5)
clf
subplot(2,2,1)
bp1 = bar(Sample_Pts,Slope_Fail_Rates_Baseline(:,1));
bp1.FaceColor = 'k';
ylim([0 1])
hold on
plot([2,6],[0.05,0.05],'-g','LineWidth',3)
xlabel('# Sample Points','FontSize',16,'FontWeight','bold')  
ylabel('Pr. Failed Baseline Slope Tests','FontSize',16,'FontWeight','bold')  
title('Baseline Normal PD samples','FontSize',16,'FontWeight','bold')
box off


subplot(2,2,2)
bp2 = bar(Sample_Pts,Slope_Fail_Rates_Baseline(:,2));
bp2.FaceColor = 'r';
ylim([0 1])
hold on
plot([2,6],[0.05,0.05],'-g','LineWidth',3)
xlabel('# Sample Points','FontSize',16,'FontWeight','bold')  
ylabel('Pr. Failed Baseline Slope Tests','FontSize',16,'FontWeight','bold')  
title('Baseline Model PD samples','FontSize',16,'FontWeight','bold')
box off

subplot(2,2,3)
bp1 = bar(Sample_Pts,Slope_Fail_Rates(:,1));
bp1.FaceColor = 'k';
ylim([0 1])
hold on
plot([2,6],[0.05,0.05],'-g','LineWidth',3)
xlabel('# Sample Points','FontSize',16,'FontWeight','bold')  
ylabel('Pr. Failed Adapted Slope Tests','FontSize',16,'FontWeight','bold')  
title('Adapted Normal PD samples','FontSize',16,'FontWeight','bold')
box off


subplot(2,2,4)
bp2 = bar(Sample_Pts,Slope_Fail_Rates(:,2));
bp2.FaceColor = 'r';
ylim([0 1])
hold on
plot([2,6],[0.05,0.05],'-g','LineWidth',3)
xlabel('# Sample Points','FontSize',16,'FontWeight','bold')  
ylabel('Pr. Failed Adapted Slope Tests','FontSize',16,'FontWeight','bold')  
title('Adapted Model PD samples','FontSize',16,'FontWeight','bold')
box off

Shape_Params
```
