## Supplemental Matlab files for "On why we lack confidence in signal-detection-based analyses of confidence": Supp_3_SDT_Kurtosis_Skew.m

```
%% This code runs simulated SDT based experiments, to determine how a 
% popular SDT-based analysis of confidence is impacted by being informed 
% by experiential distributions that are either differently skewed, or that
% have a uniform level of excess kurtosis. Simulations are of an unbiased
% observer with a fixed level of sensitivity (i.e. the means of Signal and  
% noise distributions have a constant offset), but in the first set of
% simulations different levels of excess kurtosis are sampled, and in the
% other different levels of opposite, signal and noise, skews are sampled.
%
% 51 simulated experiements are condutced degree to test each scenario,
% using the pearsrnd command to create distributions with specified meansm
% S.D.s, skews and levels of kurtosis. These inform SDT-based analyses of
% confidence, and meta d' : d' ratios are plotted as a function of
% Signal/Noise skew differences, or as a function of levels of excess
% kurtosis


%%
clear all % Clear memory
close all % close all figures
% If psignifit has not already been added to your path, do that now
addpath /Users/uqdarnol/Dropbox/Research_Projects/Toolbox/Psignifit/psignifit-master


%%
Max_Kutosis = 0.4;  % Specifies the maximal level of excess kurtosis
Max_Skew = 0.16;    % Specifies the maximal skew of a Signal or Noise distribution

N_Levels = 51;      % Number of levels of skew difference or excess kurtosis to simulate

Noise_Mu = -2;      % Mean of noise distribution (determines sensitivity)
Signal_Mu = 2;      % Mean of signal distribution
The_SD = 3;         % The S.D. of signal and nose distributions

Dec_Crit = 0;       % The category decision criterion - 0 to simulate ad unbiased observer
Conf_Crit = 7;      % The absolute (unsigned) sampled value needed to elicit high-confidence

N_Trials = 50000;   % Number of trials to simulate across Signal and Noise distributions


%% Loop to run simulations...
for N_Simulation = 1 : N_Levels
    
    % Specify the level of excess kurtosis (from 0 to Max_Kutosis)
    Curt = 3 + (((N_Simulation-1)/(N_Levels-1))*Max_Kutosis);

    % Create pseudo random distributions of Signal and Noise values
    Noise_Dist = pearsrnd(Noise_Mu,The_SD,0,Curt,1,N_Trials/2);
    Signal_Dist = pearsrnd(Signal_Mu,The_SD,0,Curt,1,N_Trials/2);
    
    % Convert Signal and Noise values into (unsigned) absolute values for
    % confidence classifications
    Noise_Confidence_Responses = abs(Noise_Dist);
    Signal_Confidence_Responses = abs(Signal_Dist);
    
    
    % Find Noise values to classify as having elicited high- or
    % low-confidence
    [vals idxs_Noise_High_Confidence] = find(Noise_Confidence_Responses >= Conf_Crit);
    [vals idxs_Noise_Low_Confidence] = find(Noise_Confidence_Responses < Conf_Crit);
    % Find Noise values to categorise as a Noise (values below category 
    % criterion) or signal (values equal to or above category criterion)
    [vals idxs_Noise_Cat1] = find(Noise_Dist < Dec_Crit);
    [vals idxs_Noise_Cat2] = find(Noise_Dist >= Dec_Crit);
    
    % Find Noise values classified as Noise with high-confidence
    [idxs_High_Conf_Noise_N vals]=intersect(idxs_Noise_High_Confidence,idxs_Noise_Cat1);
    % Find Noise values classified as Noise with low-confidence
    [idxs_Low_Conf_Noise_N vals]=intersect(idxs_Noise_Low_Confidence,idxs_Noise_Cat1); 
    % Find Noise values classified as Signal with low-confidence
    [idxs_Low_Conf_Signal_N vals]=intersect(idxs_Noise_Low_Confidence,idxs_Noise_Cat2); 
    % Find Noise values classified as Signal with high-confidence
    [idxs_High_Conf_Signal_N vals]=intersect(idxs_Noise_High_Confidence,idxs_Noise_Cat2); 
    
    
    % Find Signal values to classify as having elicited high- or
    % low-confidence
    [vals idxs_Signal_High_Confidence] = find(Signal_Confidence_Responses >= Conf_Crit);
    [vals idxs_Signal_Low_Confidence] = find(Signal_Confidence_Responses < Conf_Crit);
    % Find Signal values to categorise as a Noise (values below category 
    % criterion) or signal (values equal to or above category criterion)
    [vals idxs_Signal_Cat1] = find(Signal_Dist < Dec_Crit);
    [vals idxs_Signal_Cat2] = find(Signal_Dist >= Dec_Crit);
    
    % Find Signal values classified as Noise with high-confidence
    [idxs_High_Conf_Noise_S vals]=intersect(idxs_Signal_High_Confidence,idxs_Signal_Cat1);
    % Find Signal values classified as Noise with low-confidence
    [idxs_Low_Conf_Noise_S vals]=intersect(idxs_Signal_Low_Confidence,idxs_Signal_Cat1); 
    % Find Signal values classified as Signal with low-confidence
    [idxs_Low_Conf_Signal_S vals]=intersect(idxs_Signal_Low_Confidence,idxs_Signal_Cat2); 
    % Find Signal values classified as Signal with high-confidence
    [idxs_High_Conf_Signal_S vals]=intersect(idxs_Signal_High_Confidence,idxs_Signal_Cat2); 
    
    % Create meta-d array, and fill the first row with Noise
    % classifications X Confidence
    Meta_D_Array=[];
    Meta_D_Array(1,1) = size(idxs_High_Conf_Noise_N,2);
    Meta_D_Array(1,2) = size(idxs_Low_Conf_Noise_N,2);
    Meta_D_Array(1,3) = size(idxs_Low_Conf_Signal_N,2);
    Meta_D_Array(1,4) = size(idxs_High_Conf_Signal_N,2);
    
    % Fill second row with Signal classifications X Confidence
    Meta_D_Array(2,1) = size(idxs_High_Conf_Noise_S,2);
    Meta_D_Array(2,2) = size(idxs_Low_Conf_Noise_S,2);
    Meta_D_Array(2,3) = size(idxs_Low_Conf_Signal_S,2);
    Meta_D_Array(2,4) = size(idxs_High_Conf_Signal_S,2);
    
    % Conduct SDT-based analysis of confidence
    Analysis = fit_meta_d_MLE(Meta_D_Array(1,:),Meta_D_Array(2,:));
    % Store estimated d' values for excess kurtosis simmulations
    dPrimes_Kurt(N_Simulation)=Analysis.da;
    % Store estimated meta d' values for excess kurtosis simmulations
    Meta_dPrimes_Kurt(N_Simulation)=Analysis.meta_da;
    % Store the simulated level of excess kurtosis   
    Curts(N_Simulation) = Curt;
    
    %% Now we run simulations testing for the impact of different Signal and Noise skews
    
    % Set excess kurtosis to 0 (a kurtosis of 3 is consistent with a normal
    % distribution)
    Curt = 3;
    
    % Set skew to a level ranging from a maximal Left (-'ve) to a maximal
    % right (+'ve) skew (of Max_Skew)
    % distribution)
    Skew = -Max_Skew + (((N_Simulation-1)/(N_Levels-1))*(Max_Skew*2));
    
    % Create pseudo random distributions of Signal and Noise values, with
    % opposite skews
    Noise_Dist = pearsrnd(Noise_Mu,The_SD,-Skew,Curt,1,N_Trials/2);
    Signal_Dist = pearsrnd(Signal_Mu,The_SD,Skew,Curt,1,N_Trials/2);
    
    % Convert Signal and Noise values into (unsigned) absolute values for
    % confidence classifications    
    Noise_Confidence_Responses = abs(Noise_Dist);
    Signal_Confidence_Responses = abs(Signal_Dist);
    
    % Find Noise values to classify as having elicited high- or
    % low-confidence
    [vals idxs_Noise_High_Confidence] = find(Noise_Confidence_Responses >= Conf_Crit);
    [vals idxs_Noise_Low_Confidence] = find(Noise_Confidence_Responses < Conf_Crit);
    % Find Noise values to categorise as a Noise (values below category 
    % criterion) or signal (values equal to or above category criterion)
    [vals idxs_Noise_Cat1] = find(Noise_Dist < Dec_Crit);
    [vals idxs_Noise_Cat2] = find(Noise_Dist >= Dec_Crit);
    
    % Find Noise values classified as Noise with high-confidence
    [idxs_High_Conf_Noise_N vals]=intersect(idxs_Noise_High_Confidence,idxs_Noise_Cat1);
    % Find Noise values classified as Noise with low-confidence
    [idxs_Low_Conf_Noise_N vals]=intersect(idxs_Noise_Low_Confidence,idxs_Noise_Cat1); 
    % Find Noise values classified as Signal with low-confidence
    [idxs_Low_Conf_Signal_N vals]=intersect(idxs_Noise_Low_Confidence,idxs_Noise_Cat2); 
    % Find Noise values classified as Signal with high-confidence
    [idxs_High_Conf_Signal_N vals]=intersect(idxs_Noise_High_Confidence,idxs_Noise_Cat2); 

    
    % Find Signal values to classify as having elicited high- or
    % low-confidence
    [vals idxs_Signal_High_Confidence] = find(Signal_Confidence_Responses >= Conf_Crit);
    [vals idxs_Signal_Low_Confidence] = find(Signal_Confidence_Responses < Conf_Crit);
    % Find Signal values to categorise as a Noise (values below category 
    % criterion) or signal (values equal to or above category criterion)
    [vals idxs_Signal_Cat1] = find(Signal_Dist < Dec_Crit);
    [vals idxs_Signal_Cat2] = find(Signal_Dist >= Dec_Crit);
    
    % Find Signal values classified as Noise with high-confidence
    [idxs_High_Conf_Noise_S vals]=intersect(idxs_Signal_High_Confidence,idxs_Signal_Cat1);
    % Find Signal values classified as Noise with low-confidence
    [idxs_Low_Conf_Noise_S vals]=intersect(idxs_Signal_Low_Confidence,idxs_Signal_Cat1); 
    % Find Signal values classified as Signal with low-confidence
    [idxs_Low_Conf_Signal_S vals]=intersect(idxs_Signal_Low_Confidence,idxs_Signal_Cat2); 
    % Find Signal values classified as Signal with high-confidence
    [idxs_High_Conf_Signal_S vals]=intersect(idxs_Signal_High_Confidence,idxs_Signal_Cat2); 
    
    % Create meta-d array, and fill the first row with Noise
    % classifications X Confidence
    Meta_D_Array=[];
    Meta_D_Array(1,1) = size(idxs_High_Conf_Noise_N,2);
    Meta_D_Array(1,2) = size(idxs_Low_Conf_Noise_N,2);
    Meta_D_Array(1,3) = size(idxs_Low_Conf_Signal_N,2);
    Meta_D_Array(1,4) = size(idxs_High_Conf_Signal_N,2);
    
    % Fill second row with Signal classifications X Confidence
    Meta_D_Array(2,1) = size(idxs_High_Conf_Noise_S,2);
    Meta_D_Array(2,2) = size(idxs_Low_Conf_Noise_S,2);
    Meta_D_Array(2,3) = size(idxs_Low_Conf_Signal_S,2);
    Meta_D_Array(2,4) = size(idxs_High_Conf_Signal_S,2);
    
    % Conduct SDT-based analysis of confidence
    Analysis = fit_meta_d_MLE(Meta_D_Array(1,:),Meta_D_Array(2,:));
    
    % Store estimated d' values for different skew simulations
    dPrimes_Skew(N_Simulation)=Analysis.da;
    % Store estimated meta d' values for different skew simulations
    Meta_dPrimes_Skew(N_Simulation)=Analysis.meta_da;    

    % Store the Signal skew value that was simulated
    Skews(N_Simulation) = Skew;
end

% Calculate meta d' : d' ratios
Kurts_Ratios = Meta_dPrimes_Kurt./dPrimes_Kurt;
Skews_Ratios = Meta_dPrimes_Skew./dPrimes_Skew;

% Create figure, and plot the results of excess kurtosis simulations in
% subplot 1 
figure
subplot(2,1,1)
plot([0,0.4],[1,1],'-g','LineWidth',5)
hold on
scatter(Curts-3,Kurts_Ratios,'k','filled')
xlabel('Avg. Excess Kurtosis','FontSize',16,'FontWeight','bold')  
ylabel('Meta dPrime : dPrime Ratios','FontSize',16,'FontWeight','bold')  
box off
ax=gca;
ax.FontSize = 14; 

% Plot results of different skew simulations in subplot 2. Note that we
% double the individual skew to get the sampled difference
subplot(2,1,2)
plot([-0.4,0.4],[1,1],'-g','LineWidth',5)
hold on
scatter(2*Skews,Skews_Ratios,'k','filled')
xlabel('Skew Difference (Noise-Signal)','FontSize',16,'FontWeight','bold')  
ylabel('Meta dPrime : dPrime Ratios','FontSize',16,'FontWeight','bold')  
box off
ax=gca;
ax.FontSize = 14; 
```
